# Epigenetic variations are accompanying landmarks of freshwater adaptation in threespine sticklebacks

**DOI:** 10.1101/2022.08.22.504457

**Authors:** Artemiy Golden, Alexey Starshin, Alexandr Mazur, Nikolai Mugue, Daria Kaplun, Artem Artemov, Ekaterina Khrameeva, Egor Prokhortchouk

## Abstract

For evolutionary biology, the phenotypic consequences of epigenetic variations and their potential contribution to adaptation and diversification are pressing issues. Marine and freshwater sticklebacks represent an ideal model for studying both genetic and epigenetic components of phenotypic plasticity that allow fish to inhabit water with different salinity. Here, we applied single-cell genomics (scRNA-seq and scATAC-seq) and whole-genome bisulfite sequencing to characterize intercellular variability in transcription, the abundance of open chromatin regions, and CpG methylation level in gills of marine and freshwater stickleback morphs. We found little difference in overall transcriptional variance between the morphs but observed significant changes in chromatin openness variance. In addition, genomic divergence islands (DIs) coincided with regions of increased methylation entropy in freshwater fish. Moreover, analysis of transcription factor binding sites within DIs revealed that СTCF motifs around marker SNPs were significantly enriched within the region. Altogether, our data show that increased epigenetic variance accompanies the adaptation of marine sticklebacks to freshwater.

## INTRODUCTION

Any kind of phenotype in complex organisms results from the action of two forces - genetic and environmental. However, even external conditions exert their influence on the organism through modulation of genome functioning. The main mechanisms of gene regulation in vertebrates involve DNA methylation, combinatorial histone code, noncoding RNAs. If the environment suddenly changes, these mechanisms are the immediate frontline to adopt the genotype to the new surroundings.

Any kind of naturally observed parameter can be described in terms of statistics: expected value, mean, dispersion etc. For instance, each cell in a certain population for each gene produce a digital value of transcriptional strength. Thus, a population of cell is characterized by transcription of each gene in statistical terms. Single cell genomics open possibilities to consider transcription, chromatin openness, histone modifications as a random variable that has a particular distribution function for each gene/genomic region within the population. While “bulk” genomics mainly operate with the mean values, analysis of individual cells may add a concept of dispersion or variation of particular epigenetic parameter. NGS revolution generated a new methodology based on the analysis of genetic and epigenetic information of single cells. Today it is possible to analyze not only DNA methylation but information on histone modifications (Bartosovic et al., 2021) and open chromatin structures (ATAC-seq), allowing to evaluate the role of epigenetic stochasticity as an additional dimension of functional genomics in a variety of biological processes.

Similar to genetic diversity serving to increase the chances of a population to survive and reproduce, variability of epigenetic marks may provide an additional source of such changes. Until recently, most if not all studies that put epigenetic noise in focus concentrated on DNA methylation since whole-genome bisulfite sequencing naturally produced per allele information with single-nucleotide resolution. For instance, cancer cells increase DNA methylation variations in so-called Variably Methylated Regions (VMRs) (Pujadas & Feinberg, 2012). In surrounding normal somatic cells, such variations were not detected. Surprisingly, these VMRs are found at critical loci for development, such as axial pattern formation, neurogenesis, immune system development, and gut development (Pujadas & Feinberg, 2012). Later it became clear that borders between regions with high and low methylation entropy mainly coincide with borders between topologically associating domains (TADs) (Jenkinson et al., 2017).

Biologists are now discussing how environmental changes are translated into epigenetic variations of both somatic and germinal cells, what are molecular mechanisms that conform epigenetic information through sexual reproduction, and how genetic selection influences epiallele selection and diversity. For evolutionary biology, the phenotypic consequences of non-genetic inheritance (NGI) and their potential contribution to adaptation and diversification are pressing issues. The fact that parental exposure to any kind of unusual surrounding conditions may influence the progeny has been shown for morphology (Herman & Sultan, 2016), physiology (Shama & Wegner, 2014), behavior (Bohacek & Mansuy, 2015), longevity (Greer et al., 2011), and disease (Ben Maamar et al., 2019). Upon adaptation scenario, it increases offspring fitness (Shama & Wegner, 2014; Schunter et al., 2017; Ryu et al., 2018; Herman et al., 2012). Epigenetics has been shown to contribute to the adaptation of whole populations to the altered environment, thus maintaining new phenotypes in generations (Yin et al., 2019; English et al., 2015; Shea et al., 2011). The overall significance of epigenetics for evolutionary processes depends on the relative importance of NGI and genetic variation in creating phenotypic diversity (Baugh & Day, 2020). Indeed, environmental variation can mediate the evolution of NGI regulation in roundworms (Silva et al., 2021). However, data on the variability of NGI and its genetic basis from natural populations and from vertebrates is scarce.

Several groups have studied the genetic and epigenetic adaptation of marine threespine sticklebacks to freshwater (Jones et al., 2012; Terekhanova et al., 2014, Artemov et al., 2017)). Such a switch happened around 700 years ago when fish from the White sea were isolated in Mashinnoe lake due to the steady glacio-isostatic rise of the coast at the rate of 3.8 mm per year. Since then, freshwater morph has adapted to low-salt water and changed some phenotypic traits. Briefly, seawater and freshwater genomes are different at a few dozens of divergency islands, showing significantly different allele frequencies between two morphs (Terekhanova et al., 2019; Terekhanova et al., 2014). Apart from genetic features, there are multiple differentially methylated regions that distinguish them (Artemov et al., 2017). Moreover, we found that individual gill cell genomes of freshwater morph are characterized by higher interindividual dispersion of CpG methylation within CG-rich regions when compared to their seawater relatives.

Here, we used single-cell genomics and transcriptomics to characterize intraindividual epigenetic dispersion (terms dispersion, entropy, or variation will be further used in intraindividual context) in seawater and freshwater sticklebacks: highly abundant transcripts, number and coverage of open chromatin regions, stochasticity of DNA methylation. These characteristics were studied both for the whole genomes and specifically for DIs. Altogether, we aimed to find an epigenetic component in the biology of stickleback adaptation to the altered environment with different salinity.

## RESULTS

### scRNA-seq analysis allows transcriptional characterization of different cell types in the gills

To explore gene expression changes associated with adaptations to water salinity conditions, we studied the gills of two marine sticklebacks and two freshwater sticklebacks (Fig. 1A). Gills contact the surrounding water directly, thus being highly affected by the water salinity. Sequencing of RNA from gills’ individual cells (scRNA-seq) resulted in a total of 19,964 cells with at least 100 unique detected molecules: 1,905 and 8,002 cells in two marine sticklebacks, 4,753 and 5,304 cells in two freshwater sticklebacks (Suppl. Fig. 1,2). Thus, 50% of cells were derived from marine sticklebacks and 50% from freshwater sticklebacks. The extent of marine-freshwater expression divergence agreed well between the averaged scRNA-seq and the bulk RNA-seq (Rastorguev et al., 2018) data (Pearson’s R=0.2, p-value<10^-10^, Fig. 1B).

**Fig. 1.**
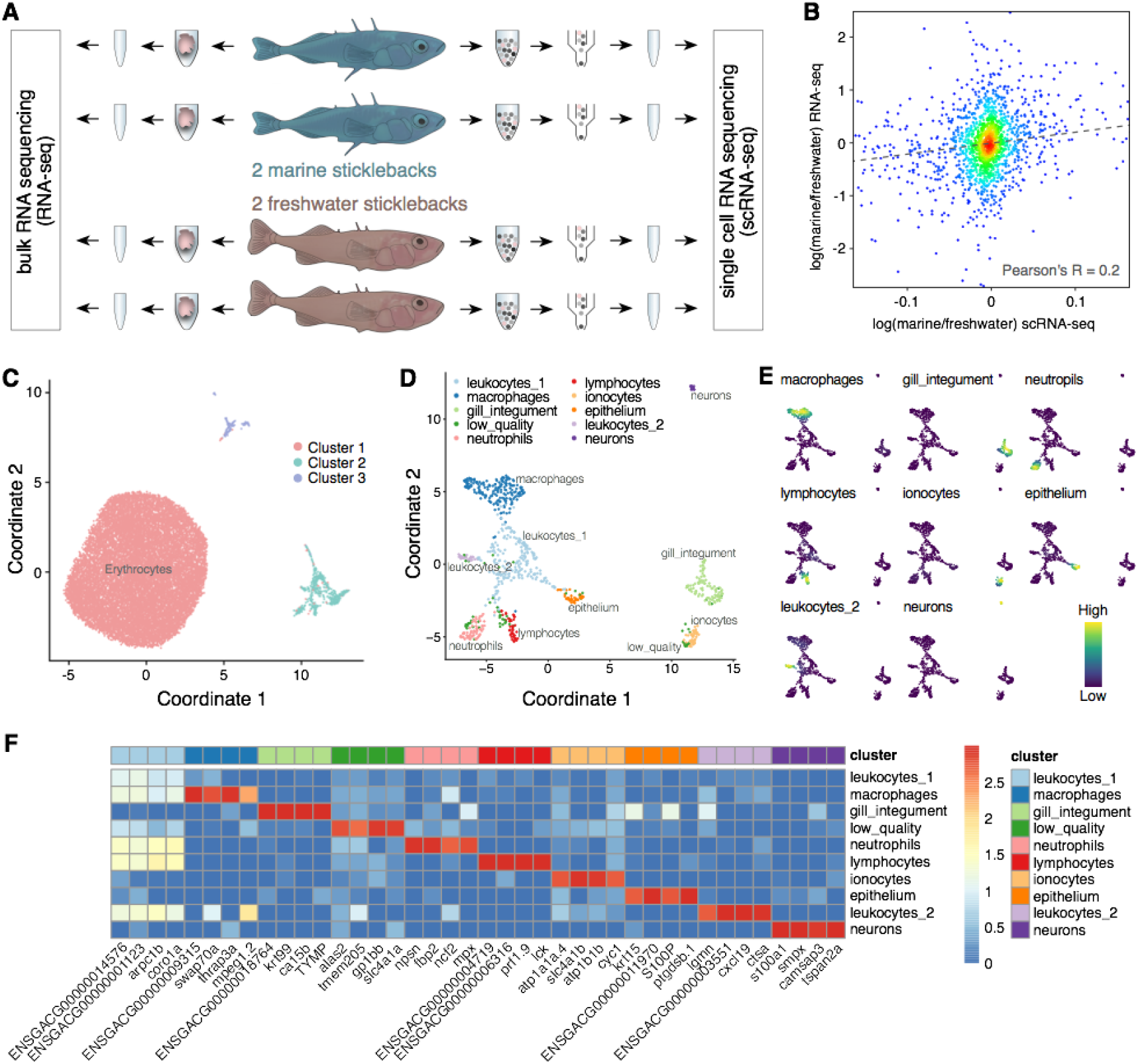
Transcriptomics in marine and freshwater sticklebacks. (A) Study design. Gene expression was profiled in gills of marine and freshwater sticklebacks using conventional RNA sequencing (bulk RNA-seq) and single-cell RNA sequencing (scRNA-seq). (B) Correlation of marine to freshwater expression fold change between bulk RNA-seq and averaged snRNA-seq datasets in log scale. Pearson’s R = 0.2, p-value < 10^-10^. (C) UMAP plot of 19,964 cells colored by cluster identity. (D) UMAP plot of 804 cells colored by cluster identity. Erythrocytes were removed. (E) Projection of expression levels averaged across cell type marker genes onto the UMAP plot shown in panel 1D. Labels above the UMAP plots mark cell types. (F) Average expression levels of cell-type marker genes in clusters. The same marker genes were used in panel 1E.

**Fig. 2.**
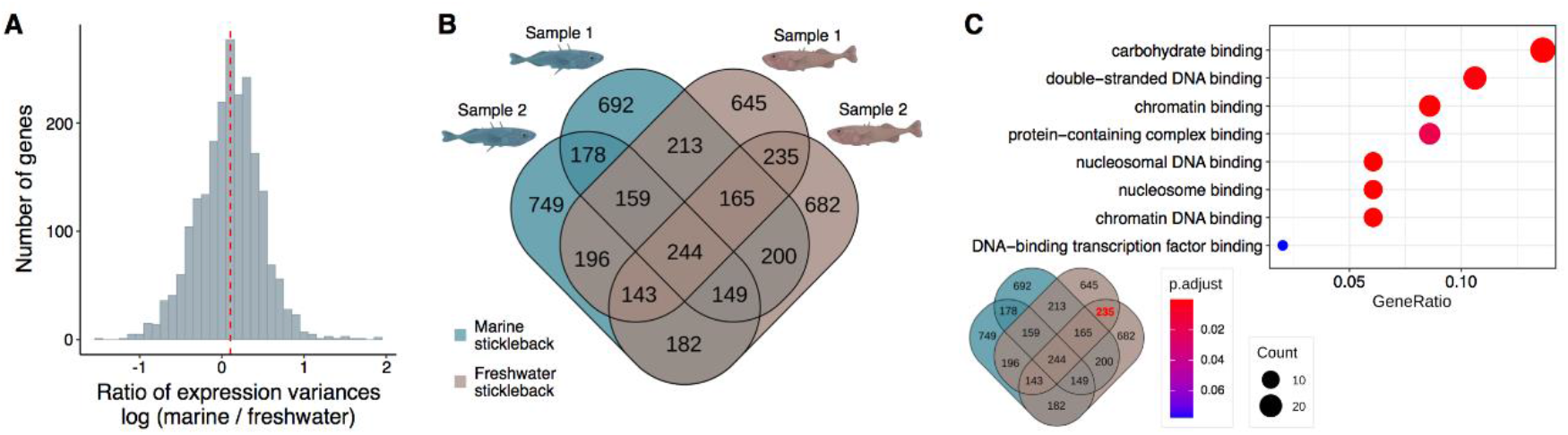
Transcriptional diversity in marine and freshwater sticklebacks. (A) The ratio of expression variance among marine stickleback cells to the expression variance among freshwater stickleback cells (in log scale). The dashed line represents the median of ratios among all genes. (B) Venn diagram of overlapping HVG defined in each sample of marine (green) and freshwater (brown) sticklebacks separately. Numbers indicate HVG. (C) Functional enrichment analysis of 235 HVG specific to freshwater sticklebacks, performed according to the Gene Ontology (GO) database (Carbon et al., 2020). Only erythrocyte cells are analyzed in panels A-C.

Visualization of the total expression profiles across cells revealed that the majority of them represented erythrocytes (Fig. 1C, Suppl. Table 2 - cluster markers). The remaining 804 cells could be separated into several clusters (Fig. 1D). Several resolutions were tested for clustering (0.05, 0.1, 0.3, 1.5, Suppl. Fig. 3). The best resolution for separating groups of cells with distinct patterns of expression was 0.3. Higher resolution divided cells into several clusters with similar content and expression of marker genes, while lower resolutions did not provide enough separation. We identified marker genes for each of these clusters (Suppl. Table 3 - cluster markers) and plotted four best markers for each cluster on a heatmap (Fig. 1F, Suppl. Fig. 4). To assign cell types to the resulting cell clusters, we manually searched marker genes that had zebrafish orthologs in the ZFIN database (Ruzicka et al., 2018). For each cluster, we searched for a recurring pattern of tissue specificity in categories for its marker genes in the ZFIN database. At the first step, *coro1a*, *arpc1b*, *zgc:64051* were identified as clear markers of immune cells, clarifying further annotation. The macrophage cluster showed high expression of *mpeg1.2*, which is a macrophage-specific marker (Rougeot et al., 2019), and additional markers *cd79a* and *swap70a*. Neutrophils were easily identifiable by several highly specific markers: *npsn*, *mpx,* and *mmp9* (Di et al., 2017). The lymphocyte cluster was identified as the only one expressing T cell-specific tyrosine kinase *lck* gene; additionally, *prf1.9* expression was also exclusive for this cluster. The cluster of ionocytes, or mitochondria-rich cells, MRCs (Wilson & Laurent, 2002), had elevated expression of genes responsible for catabolism as well as several specific markers: *atp1a1a.4*, *slc4a1b*, *atp1b1*, *slc4a1b*. *Atp1b1b* and *slc4a1b,* in particular, are gill ionocyte-specific markers (Farnsworth et al., 2020). Gill integument cluster containing Pavement cells, most likely (Wilson & Laurent, 2002), had high expression of keratin genes as well as *rhcga* gene which has homology to *rhcgl1* that was identified as gill integument gene (Farnsworth et al., 2020). Gill integument cluster was the most abundant out of the clusters that expressed keratin genes enriched in the various epithelium, and it is expected to be the biggest of epithelial cell clusters in gills. There was another smaller epithelial cluster with markers *S100P*, *ptgdsb*.1, *ENSGACG00000004694* specific for it, but there was not enough information to distinguish the type of epithelium. A small cluster of neuron-like cells, possibly neuroepithelium (Wilson & Laurent, 2002), had to be separated manually based on the UMAP visualization since clustering at the resolution of 0.3 did not isolate it. However, this cluster had highly specific markers *s100a1*, *scilna*, *sptbn2*, *CKB,* all of which are supposed to be elevated in the neuronal tissue. The first cluster of leukocytes (leukocytes_1) did not show any specific markers apart from the high expression of genes typical for blood immune cells, present in all blood immune cell clusters of this dataset. The second cluster of leukocytes (leukocytes_2) had several specific markers: *cxcl19*, *ctsa*, *tcn2*. However, none of the markers provided enough information to identify which type of immune cells it corresponded to.

Further, we analyzed gene expression dispersion in marine and freshwater samples. The largest homogeneous cell cluster representing erythrocytes was selected for this analysis to explore the dispersion separately from possible effects of cell-type composition changes between marine and freshwater sticklebacks. We performed the F-test for dispersion difference between marine and freshwater samples per each expressed gene (see Methods). This analysis revealed a slight increase of intraindividual transcription dispersion in marine sticklebacks compared to the freshwater population (Fig. 2A). The same analysis repeated on all cells, including erythrocytes and other cell types, produced highly similar results, indicating that the observed slight increase of divergence within the marine stickleback population cannot be explained by cell-type composition changes (Suppl. Fig. 5).

We further searched for highly variable genes (HVG) indicating high expression diversity within either the freshwater stickleback population or the marine population. For each gene, we calculated expression variability among cells of one sample and obtained lists of HVG ranked by their variability in marine or freshwater stickleback samples (Suppl. Table 4). This analysis was performed before the sample integration step in Seurat (Hao et al., 2021) to avoid possible technical confounders associated with the procedure of sample integration.

A total of 4,832 genes were identified as HVG in at least one of four stickleback samples. A comparison of HVG lists between samples revealed that 178 genes were defined as HVG in both marine samples and neither freshwater samples (Fig. 2B). Similarly, 235 genes were defined as HVG in both freshwater samples and neither marine samples (Fig. 2B). Functional enrichment analysis of 178 marine-specific HVG did not reveal significant functional categories, according to the Gene Ontology (GO) database (Carbon et al., 2020) (Suppl. Fig. 6). By contrast, a similar analysis of 235 freshwater-specific HVG resulted in eight significant functional categories (Fig. 2C), mostly related to chromatin binding and its nested GO terms. Analysis of all cells, including erythrocytes and other cell types, produced highly similar results (Suppl. Fig. 12). This functional analysis suggests that gene expression regulation at the level of chromatin accessibility might be an important component of stickleback adaptation to altered salt concentration in water.

### scATAC-seq analysis reveals higher variability in number and coverage of open chromatin sites in seawater sticklebacks

To explore changes associated with adaptations to water salinity conditions at the level of gene expression regulation, we studied chromatin accessibility in the gills of the same specimens of marine sticklebacks and freshwater sticklebacks at the single-cell resolution. We performed the scATAC-seq experiment (Fig. 3A) and obtained clusters of cells (Fig. 3B), where a distinct cluster represented erythrocyte cells (Fig. 3C, Suppl. Fig. 21). Similarly to gene expression analysis, we performed F-test for dispersion difference (see Methods) between marine and freshwater samples in scATAC-seq peaks and observed significantly higher variability of chromatin accessibility in marine sticklebacks compared to freshwater sticklebacks (Fig. 3D). For consistency with gene expression divergence analysis, we additionally calculated F-ratio for erythrocytes only, to factor out chromatin accessibility divergence explained by cell-type composition changes, and obtained similar results (Suppl. Fig. 27).

**Fig. 3.**
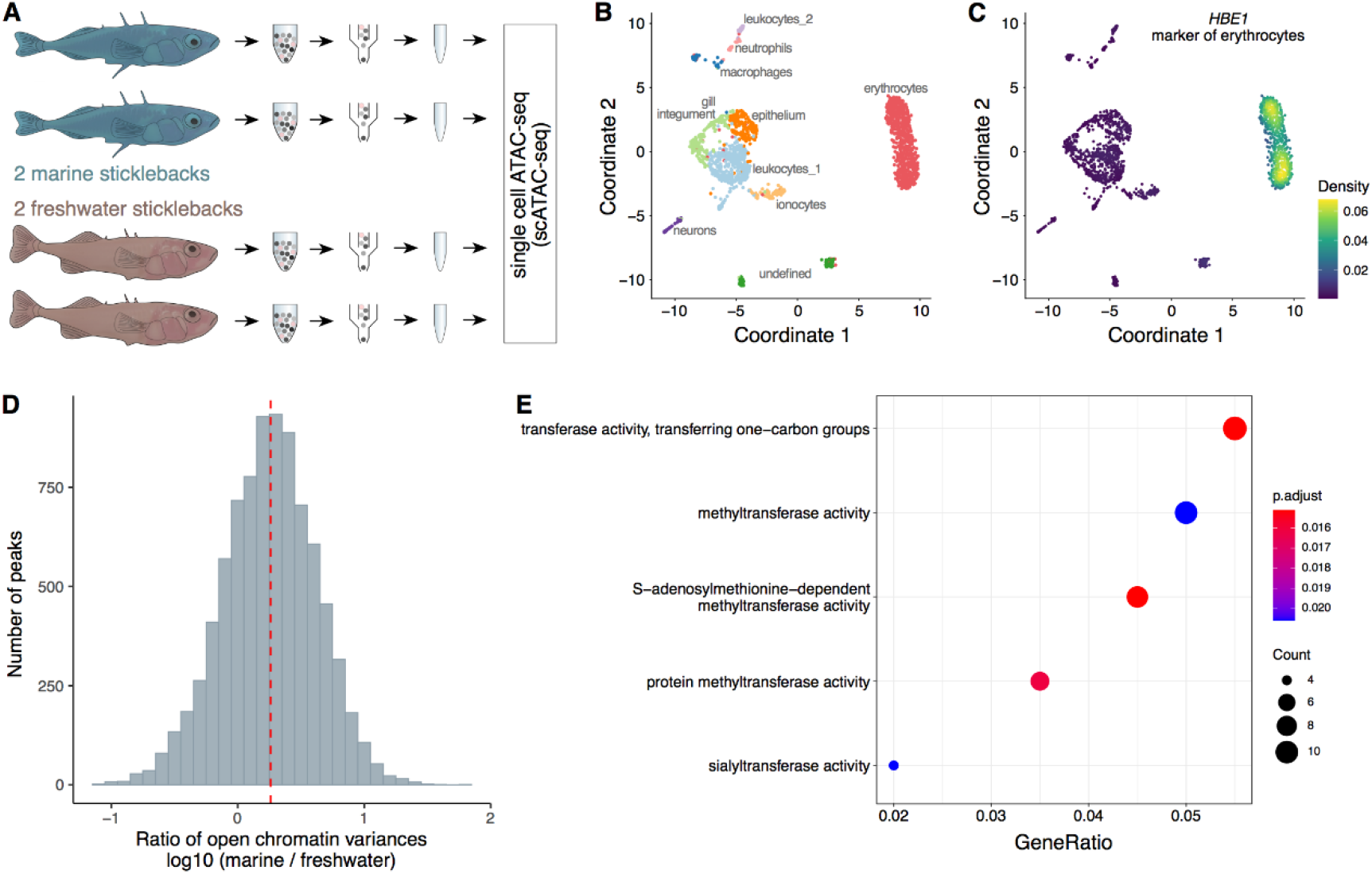
Diversity of chromatin accessibility in marine and freshwater sticklebacks. (A) Chromatin accessibility was profiled in gills of marine and freshwater sticklebacks using single-cell ATAC-seq method (sсATAC-seq). (B) UMAP plot of 1,757 cells colored by cluster identity. (C) Expression level of *HBE1* gene marker of erythrocytes in cells. (D) The ratio of chromatin accessibility variance among marine stickleback cells to the chromatin accessibility variance among freshwater stickleback cells, calculated for each peak for the unified set (in log scale). The dashed line represents the median of ratios. (E) Functional enrichment analysis of genes located next to marine-specific highly variable peaks (calculated using two metrics, see Methods), performed according to the Gene Ontology (GO) database (Carbon et al., 2020).

Functional enrichment analysis of genes located next to marine-specific highly variable scATAC-seq peaks (calculated using two metrics: mean number of non-zero peaks in a gene and mean peak height in a gene, see Methods) resulted in five significant functional categories (Fig. 3E), according to the Gene Ontology (GO) database (Carbon et al., 2020). All these categories were related to methyltransferase activity, suggesting an important role of DNA methylation in the adaptation of stickleback populations to water salinity conditions.

### DNA methylation entropy

To explore the diversity of DNA methylation in stickleback populations, we analyzed 24 freshwater and 22 marine sticklebacks, among which there were 19 males and 27 females, and calculated the methylation entropy in windows of five consecutive CpGs across the genome (see Methods). For simplicity we explain entropy as value that correlates with a number of different methylation patterns in a given window: within each window, we counted the number of possible epiallele states, normalized it for the read coverage, and defined the methylation entropy as a function of the frequency of each epiallele per window. Principle component analysis of entropy variation revealed separation of samples by the environment (PC1) and sex (PC2) (Fig. 4A-B). At the genome-wide level, mean entropy among both males and females was higher in marine fish compared to freshwaters (Fig. 4C).

**Fig. 4.**
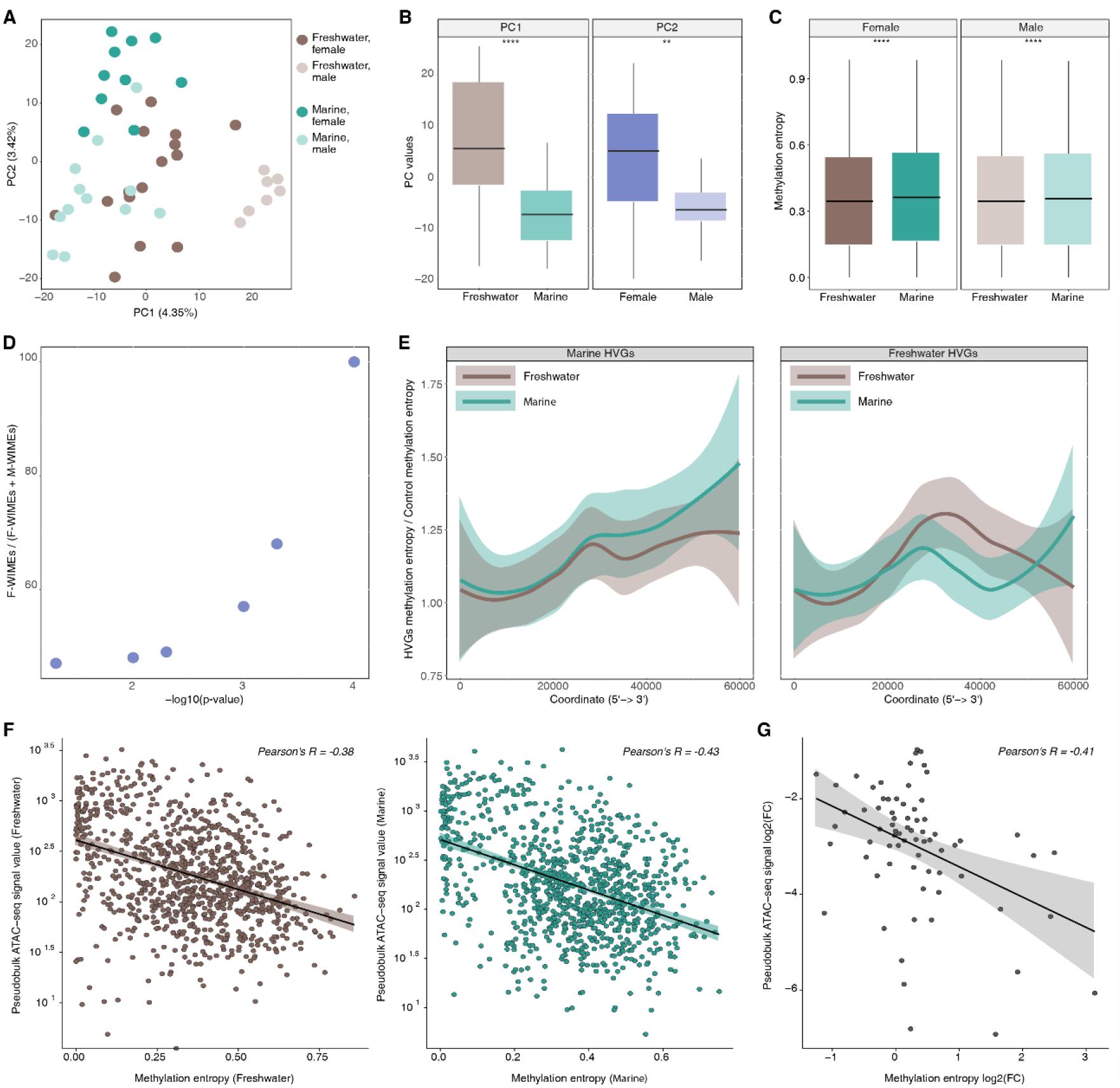
DNA methylation entropy. (A) PCA plot based on entropy variation among all analyzed samples. Each circle represents a sample. Circle colors represent environments and sexes. (B) PC1 separates samples by the environment (U test, p-value < 0.0001). PC2 separates samples by sex (U test, p-value < 0.01). (C) F-test shows differences in entropy between marine and freshwater populations at the genome-wide level (p-value < 0.0001). (D) The ratio between the windows of increased methylation entropy in freshwater (F-WIMEs) to the sum of freshwater (F-WIMEs) and marine (M-WIMEs) fish as a function of the U test p-value. (E) Entropy in the bodies of HVG was divided by the entropy of control genes. The resulting values were used to construct the distributions for two classes of HVG bodies. (F) Correlations between chromatin accessibility and averaged methylation entropy values Circles represent ATAC-seq peaks. (G) Correlation between the chromatin accessibility fold change (marine/freshwater) and the fold change of averaged methylation entropy values. Only 50 ATAC-seq peaks selected by sPLS-DA procedure (see Methods and Suppl. Fig. 35-38) are shown.

In our scRNA-seq and scATAC-seq experiments, four (two marine and two freshwater) fish were males. Thus, we used only males in our DNA methylation analysis to enable comparisons with no sex bias. By performing the U-test among males, we obtained 16,361 and 12,057 five-CpG windows (p-value < 0.001) that increased and decreased the methylation entropy in the freshwater population compared to the marine. The ratio of differential windows demonstrated a strong dependence: the higher the significance, the greater the shift towards freshwater (Fig 4D).

Next, we studied entropy distribution among the HVG and calculated the level of methylation entropy for all genes with altered transcriptional variance. Two types of X-axis scaling were used - in nucleotides and in relative units showing the percentage of gene length (0% near TSS and 100% at the gene end). Consistent with previous findings, the overall shape of the distribution displayed a significant drop in methylation entropy around TSS (Suppl. Fig. 32). Additionally, we observed a slight increase in the entropy for marine fish at a distance of around 5 Kb from the TSS and further explored it at the scaling of 60kb (Suppl. Fig. 33,34). Moreover, for both marine HVG and freshwater HVG, we observed an increase within gene bodies in marine fish. Higher overall entropy in marines (Fig. 4C) raises the possibility that the observed effect for gene bodies is not specific for HVG. Normalization by the remaining expressed genes showed a statistically significant increase of methylation entropy in gene bodies of HVG: in marine HVG, the entropy was higher in marines, while in freshwater HVG, the entropy was higher in freshwaters (Fig. 4E). Therefore, higher entropy of DNA methylation in gene bodies produces variation in gene transcription.

Further, we studied the dependencies between the methylation entropy and chromatin accessibility. First, we calculated Pearson correlation coefficients between the absolute values of entropy and pseudobulk ATAC-seq signal values for freshwater and marine fish separately, and observed strong and significant negative correlations in both cases (Pearson’s R = -0.38, p < 10^-10^ and Pearson’s R = -0.43, p < 10^-10^ for freshwater and marine fish, respectively; Fig. 4F). Next, we considered fold changes between marine and freshwater populations instead of absolute values of methylation entropy and ATAC-seq signal and observed a strong and significant negative correlation for a subset of ATAC-seq peaks, which allows the best classification of our samples into two classes, freshwater and marine, according to the sPLS-DA procedure (see Methods; Fig. 4G; Suppl. Fig. 35-38; Pearson’s R = -0.37, p = 0.048).

### Freshwater fish show elevated DNA methylation entropy in DIs and DI-like regions

We next studied how sequence divergence is correlated with methylation entropy. Many studies have reported that the marine stickleback genome has hundreds of low frequency standing adaptive haplotypes that all sweep to fixation upon freshwater colonization (Terekhanova et al., 2019; Roberts Kingman et al. 2021). These haplotypes are old, and in general are marked by substantial sequence divergence between marine and freshwater haplotypes.

Genetics traces the difference between freshwater and marine populations. According to previous studies, the whole-genome fixation index (Fst) for the populations is significantly lower than Fst in divergency islands (Terekhanova et al., 2019). These results suggest that the main genetic variation explained by the environment is concentrated within marker SNPs of the DIs. To check whether DNA methylation entropy is also different in DIs, we analyzed DIs and +/- 50 kb flanking genomic regions. We also performed the methylation entropy normalization to exclude the potential impact of genetic events such as C↔T transitions. Thus, we calculated entropy across genome of marine and freshwater fish using information from non-bisulfite-treated DNA. C to T transitions within CpG would artificially produce stably non-methylated cytosines. While T to C transitions within TpG would create artificially methylated C. Real epigenetic entropy is a result of normalization of observed entropy from bisulphite converted genomes by the entropy that comes from genetic events. All 19 DIs produced significantly higher DNA methylation entropy within the DIs (Fig. 5A).

**Fig. 5.**
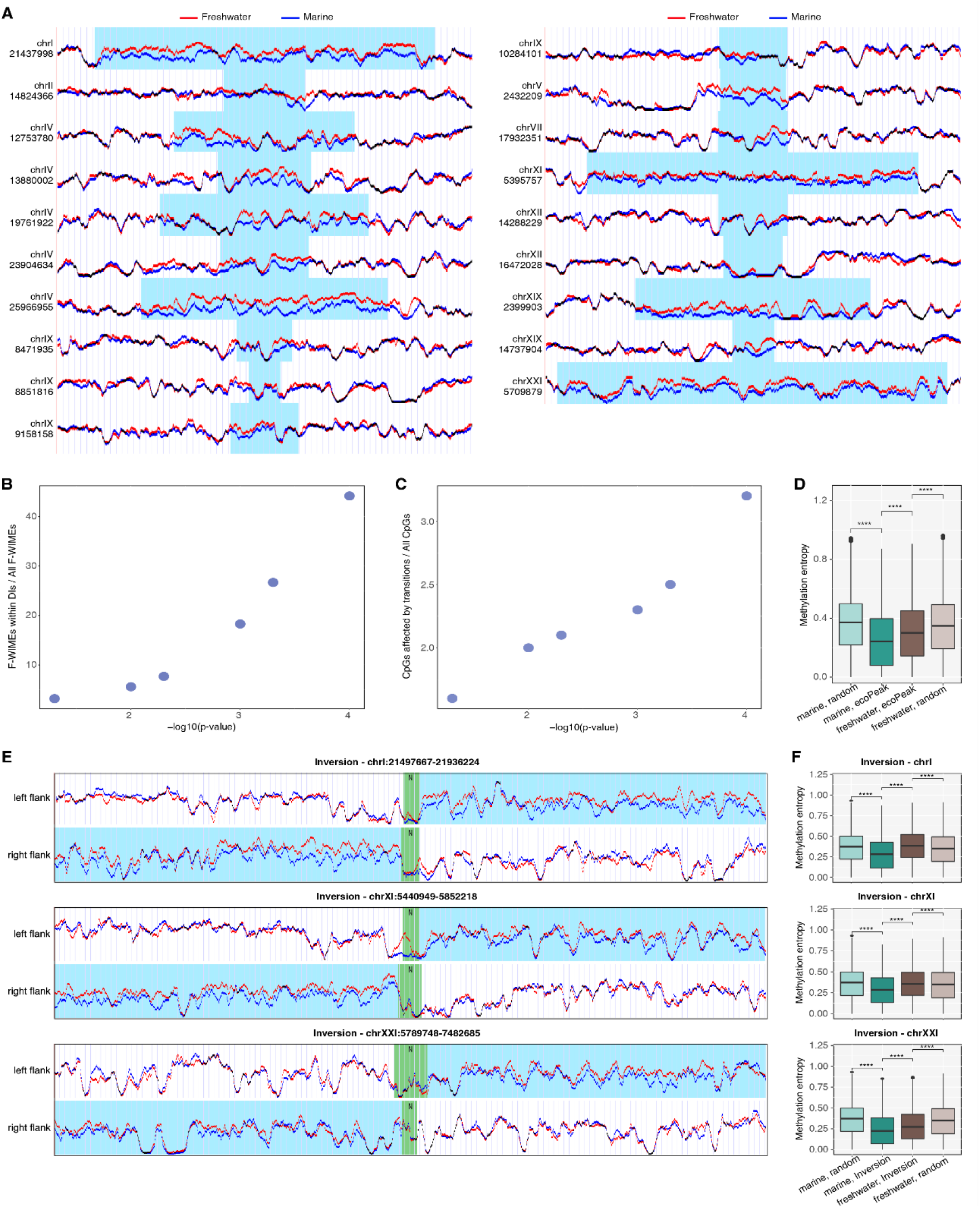
Freshwater fish show elevated DNA methylation entropy in DIs and DI-like regions. (A) DNA methylation entropy of freshwater (red) and marine fish (blue) within 19 DIs and 50-kb flanking regions. The scaling of each DI is different and adjusted to the flanking regions to contain 50 kb. DIs are highlighted by blue boxes. Coordinates correspond to the Stickleback Feb. 2006 (Broad/gasAcu1) genome assembly. (B) The relative amount of F-WIMEs located in DIs. (C) The number of transitions C->T and G->A occurring within CpG dinucleotides in DIs given as a function of the U test p-value. (D) DNA methylation entropy in the global specific EcoPeaks shows the opposite trend compared to random regions. (E) DNA methylation entropy of freshwater (red) and marine (blue) fish within breakpoints of the inversions, flanking inverted repeats (highlighted in green), and 100-kb flanking regions. Flanks included in the chromosomal inversions are highlighted in blue. (F) Boxplot representation of the methylation entropy distribution for the same inversions as in panel E.

We next estimated the abundance of five-CpG windows with increased entropy in freshwaters within the DIs as a function of U test significance and calculated a proportion of the windows in DIs among all windows (Fig. 5B). 45% of the windows with increased entropy in freshwaters concentrated in DIs (U test p-value = 0.0001).

Thus, some DIs that are divergent in sequence are characterized with differential entropy level but are not differentially methylated. Vice versa, 50% of all difference in entropy is concentrated within DIs. These results suggest some uncoupling between the genome and epigenome.

To additionally dissect the presumable genetic effect on the F-WIMEs (<u>W</u>indows with <u>I</u>ncreased <u>M</u>ethylation <u>E</u>ntropy in <u>F</u>reshwaters), all marker SNPs in Dis were tested for coincidence with CpGs. While transversions were filtered out by the pipeline (see Methods), transitions could create a potential source for the entropy bias. Indeed, for a CpG in freshwaters that matched a TpG in marines, the entropy within the window containing the CpG was obligatory increased in freshwaters. 100% of marine reads were classified as unmethylated (less chaos), while freshwaters had a chance to produce both methylated and unmethylated reads (more chaos). Accordingly, we detected 3.2% of CpG being affected by C->T and G->A transitions (in absolute numbers, С(f)->T(s) = 862 and G(f)->A(s) = 117; Fig. 5C). Moreover, some transitions, although with two-fold lower rates, occurred in a way that could potentially increase the entropy in marine sticklebacks (T(f)->C(s) = 480 and A(f)->G(s)=121).

In addition to 19 presented DIs, we examined how the methylation entropy is distributed in other known stickleback loci that drive the adaptation process. For example, the (Roberts Kingman et al. 2021) paper presents a newly defined set of EcoPeaks (haplotypes seen to convergently evolve in freshwater conditions) and TempoPeaks (haplotypes that evolve in contemporarily evolving ponds). While we observed a small decrease in entropy in freshwater environments genome-wide, specific Eco and TempoPeaks demonstrated a similar behavior of entropy to DIs (Suppl. Fig. 39, 40). This effect was most noticeable in global specific EcoPeaks (Fig. 5D). In sensitive Eco and TempoPeaks, the direction of entropy change coincided with the genome-wide picture (Suppl. Fig. 41-43).

In addition, we studied the inversions on chromosomes 1, 11, and 21, which were involved in the adaptation process (Jones et al. 2012). Since there was a rather large intersection between these inversions and DIs (Jaccard index = 0.77), we expected that the entropy in the inverted regions would be significantly higher in freshwater fish (Fig. 5E-F).

Overall, our data suggest that DIs represent a significant fraction of the fish genome with characteristically increased methylation entropy in freshwater fish. This difference cannot be explained solely by genetic variability and, thus, it represents a unique epigenetic feature of the sticklebacks adaptation.

### Enrichment in motifs within DIs

Genetic variability cannot fully explain WIME’s in DIs. However, it may create multiple motifs for the binding of key transcription or chromatin factors that, in turn, through interactions with de novo methyltransferases, cause an increase in the methylation entropy. Thus, we analyzed ATAC-seq peaks within DIs to focus on potential regulatory elements concentrated in open chromatin regions. Short 11-bp sequences (marker SNPs inside ATAC-seq peaks +/- 5 flanking nucleotides) were considered as potential binding sites for methylation modifying factors. A total of 3,089 such sequences were subjected to the enrichment analysis of known motifs and *de novo* motifs using HOMER software (Heinz et al., 2010). Interestingly, we found the CTCF binding site (CTCF-bs) at the top of the list containing the most presented motifs. Notably, CTCF-bs enrichment within DIs was three-fold higher compared to the background sequences (Suppl. Table 9). Although HOMER operates with motifs defined mainly for humans and mice by default, we can be confident in the detected enrichment of CTCF-bs in the stickleback genome due to the conservative nature of the CTCF-bs sequence (Kadota et al., 2017), as well as very dramatic and significant three-fold enrichment.

## DISCUSSION

Three-spined sticklebacks have been in the focus of evolutionary biologists since the late 1960s (Hagen, 1967). Small, easily accessible polymorphic fish were available for scientists in America, Europe, and Asia. Stickleback became popular as a model to study different aspects of adaptation: the role of a diet in explanation of a difference between benthic and limnetic morphs (Day et al., 1994); body shape changes between stream and lake fish (Hendry et al., 2002); marine and freshwater populations (Terekhanova et al., 2014; Jones et al., 2012). In all these cases, genetics was an obvious field of science that formed a basis for measurable characteristics to describe morphs. However, diet, water flow, and water salinity have an obvious environmental constituent that affects not only the natural choice of best genotypes but also forms favorable epigenetic landscapes that facilitate the survival, growth, and reproduction of fish in particular surrounding circumstances. Because of the incomparable time that is needed to establish fitted features, epigenetics is preferable to genetics to adopt fast changes. However, epigenetic landscapes such as DNA methylation cooperate with genotypes: upon activation of adaptive gene reprogramming, some genotypes may be more or less favorable to gene functioning, and vice versa, epigenetic changes may compensate for unfavorable genotypes. Moreover, epigenetics produces an additional level of within-population diversity, resulting in more adaptive power of a species. Accordingly, we focused in this work on the dispersion of the magnitude of gene transcription, chromatin accessibility, and DNA methylation. Apart from genetic diversity, dispersion is an additional characteristic of the population which may point to its ability to adopt changes on the level of gene functioning or reflect the variability of the environment. In the round goby fish, maternal RNA expression levels correlated with the water temperature experienced by the mother before oviposition and identified temperature-responsive gene groups such as core nucleosome components or the microtubule cytoskeleton (Adrian-Kalchhauser et al., 2018). The more variation in temperature, like in wild nature, the more dispersion in RNA level. Presumably, one might detect very low RNA dispersion in cultured fish compared to those in wild nature.

Previously, we have shown that CpG-rich regions of freshwater sticklebacks maintain higher levels of DNA methylation dispersion among individual fish compared to marines. Here, we tested its possible consequences on transcription and chromatin. We did not find dramatic changes in the dispersion of gene transcription between two morphs in the scRNA-seq experiment. Single-cell transcriptomics quantitatively detects RNAs from highly expressed genes; thus, variations in expressions of lowly represented genes might be obscured by low coverage sequencing of thousands of individual cells. However, HVG were still detected, and GO analysis has pointed to chromatin factors as most dispersed in their transcriptional level. Indeed, we have detected higher variability in the number and coverage of open chromatin sites in seawater sticklebacks. However, these sites were located in genes with low transcription that were undetectable by scRNA-seq. Thus, we can not conclude that higher variations in open chromatin sites cause a higher transcriptional variance. However, GO analysis of ATAC-seq HVG led us to the hypothesis that marine sticklebacks may have higher variance in the level of enzymes that maintain DNA methylation homeostasis. Bisulfite sequencing has confirmed an increase in DNA methylation entropy at the whole-genome level in marine fish. In males, the entropy difference was more profound than in females. Since all fish were almost the same age, a possible explanation for the observed sex-dependent entropy difference may be the action of female-specific hormones, which somehow modulate actors of the methylation pathway.

The genetic difference between <u>m</u>arine and freshwater sticklebacks is mainly concentrated in DIs (mDIs and fDIs, correspondingly) with much lower Fst across the rest of the genome (Terekhanova et al., 2019). Since the sticklebacks are originally seawater animals, each SNP within DIs has a relatively sharp Linkage Disequilibrium (LD) peak in marine sticklebacks : the fish lived in elevated salt conditions for millions of years. The genome of freshwater fish is a result of the selection of particular haplotypes that eventually form DIs with relatively linked SNPs, which leads to a broad peak of LD within DIs in freshwater fish. Elevated entropy of DNA methylation in fDIs may create a new interface for transcription factors - some of them are stripped from methylated DNA, but the others may bind to their consensus sites that contain m5C and resemble T. Thus, increased variability in DNA methylation may compensate for the decrease in the genetic variability of fDIs in the sense of the diversity of bound transcription factors. Indeed, results of scATAC-seq experiments show no significant changes in the number and coverage of open chromatin regions within DIs, suggesting that chromatin structure is upon adaptation. The borders between ordered and disordered DNA methylation coincide with borders between TADs (Jenkinson et al., 2017) are bound by CTCF (Dixon et al., 2012; Nora et al., 2012). Here, we observe that СTCF motifs are enriched around marker SNPs of the DIs. Adaptation results in a shift of frequencies of the marker SNPs and may change the frequency of appearance of the CTCF binding domain, which in turn may affect TAD boundary organization. Further studies are needed to reveal the role of CTCF and TAD structure in the adaptation of sticklebacks. In general, more and more data shows that modulation of chromatin structure is an important player in evolution. Our observation may be the first example where chromatin acts in the generation of morphs at a relatively short time interval (around 800 years). From a more locus-specific perspective, multiple lines of evidence, from biochemistry to comparative genomics, indicate that chromatin influences the local mutation rate (Chen et al., 2012; Prendergast & Semple, 2011; Tolstorukov et al., 2011; Warnecke et al., 2008).

Linking phenotypes to the DIs is problematic in sticklebacks. Clearly, freshwater adaptation occurs via a genome-wide program of haplotype replacement. But we do not understand what phenotypes affect nearly all of these haplotypes. The present study adds a much-needed epigenetic dimension to stickleback adaptation. However, one should be much careful about conclusions that confound correlation with causation. Do these regions "enable adaptation" or instead correlate with adaptation? Did these mutations increase in frequency "to allow more flexibility in functional states"? It remains unclear because separating the flexibility in functional states from changes in gene expression seems impossible. The finding that genetic divergence within DIs happens in part due to nucleotide substitutions within CTCF binding sites provides us with a hope that future studies may link genetic and epigenetic flexibility with functional gene biology of adaptation.

## METHODS

### Samples

Fish were collected in the Mashinnoe Lake (freshwater morph, 66°17.749N, 33°21.829E, estimated age 700 years) and from marine shore at White Sea Biological Station (marine morph, 66°57.040N, 33°10.400E). Gills were cut with sterile scissors. Gills were thoroughly washed with chilled 1 × PBS and transferred to a Petri dish. The tissue was cut into small pieces with sterile scissors and washed twice with chilled 1 × PBS. The pellet was trypsinized with 200 μl TrypLE ™ Express Enzyme (Gibco, USA) for 2 minutes. One ml of fetal bovine serum (FBS; Hyclone, USA) was added to the cell suspension to inhibit trypsin activity. The cells were harvested by filtering the cell suspension through a filter (80 microns). The filtrate was centrifuged at 2000 rpm for 5 minutes.

### Isolation of nuclei

Nuclei were isolated according to the 10x Genomics protocol for “Nuclei Isolation for Single Cell Multiome ATAC + Gene Expression Sequencing’’ available at https://www.10xgenomics.com/. The cells were washed 2 times with 1 × PBS + 0.04% BSA, and the number of cells was determined. Nuclei were isolated from 100,000-1,000,000 cells. Added 100,000-1,000,000 cells to a 2 ml microcentrifuge tube. It was centrifuged at 300 rpm for 5 min at 4 ° C. All supernatant was removed without destroying the cell sediment. Then we added 100 μl of chilled lysis buffer (10 mM Tris-HCl (pH 7.4), 10 mM NaCl, 3 mM MgCl2, 0.1% Tween-20, 0.1% Nonidet P40 Substitute (if using Sigma (74385) 100% solution, prepare a 10% stock), 0.01% Digitonin (incubate at 65°C to dissolve precipitate before use), 1% BSA, 1 mM DTT, 1 U/µl RNase inhibitor, Nuclease-free water), and incubated for 3-5 minutes on ice. We next evaluated the efficiency of lysis using an automatic cell counter and added 1 ml of chilled wash buffer (10 mM Tris-HCl (pH 7.4), 10 mM NaCl, 3 mM MgCl2, 1% BSA, 0.1% Tween-20, 1 mM DTT, 1 U/µl RNase inhibitor, Nuclease-free water) to the lysed cells. It was centrifuged at 500 rpm for 5 min at 4 ° C. The supernatant was removed without disturbing the pellet of the nuclei. Based on cell concentration and assuming ∼ 50% of nuclei lost during cell lysis, we resuspended in a chilled diluted nuclei buffer (1XNuclei Buffer (20X), 1mM DTT, 1 U/µl RNase inhibitor, Nuclease-free water). See Suppl. Table 8. All work procedures were performed on ice. We determined the concentration of nuclei using an automatic cell counter, and then immediately switched to Chromium Single Cell ATAC Reagent.

### Single-cell RNA sequencing (scRNA-seq)

Single-cell experiments were performed using a 10x Chromium single cell 3’ v2 reagent kit by precisely following the manufacturer’s detailed protocol to construct 10x Genomics single-cell 3’ libraries. Single-cell libraries were run using paired-end sequencing on the HiSeq1500 platform (Illumina) according to the manufacturer’s instructions.

### Single-cell ATAC sequencing (scATAC-seq)

Single-nuclei experiments were performed using a 10x Chromium Single Cell ATAC Library & Gel Bead Kit by precisely following the manufacturer’s detailed protocol to construct a 10x Single Cell ATAC Library. Single-nuclei libraries were run using paired-end sequencing on the HiSeq1500 platform (Illumina) according to the Chromium Single Cell ATAC Reagent Kits User Guide.

### scRNA-seq data processing

A total of 1,133,906,325 paired-end sequencing reads of scRNA-seq were processed using publicly available 10x Genomics software – Cell Ranger v3.1.0 [Zheng et al., 2017] (Suppl. Table 1). The sparse expression matrix generated by the Cell Ranger analysis pipeline with the list of 21,474 cells was used as input to the Seurat software v3.1 [Stuart et al., 2019].

Seurat pipeline standard quality control steps were performed, and cells were filtered for nFeature_RNA > 100 and percent of mitochondrial genes < 2 (Suppl. Fig. 1). Doublet detection was performed with Scrublet [Wolock et al., 2019]. The detected doublet rate was below 0.7% for all samples.

To account for technical variation, we performed cross-species integration. At the first step, for marine and freshwater samples separately, we performed normalization using “LogNormalize” with the scale factor of 10,000 and identified 2,000 variable features. Next, we performed cross-species integration by finding corresponding anchors in marine and freshwater samples using 30 dimensions. We then computed 50 principal components and tested their significance by JackStraw. We selected the first 20 principal components for subsequent UMAP and Seurat clustering analyses.

To compare scRNA-seq with the bulk RNA-seq datasets, we calculated differential expression between marine and freshwater sticklebacks for scRNA-seq and bulk datasets (Fig. 1B). Differentially expressed genes between freshwater and marine samples in scRNA-seq were identified with the Seurat *FindMarkers* function.

All cells in the scRNA-seq dataset were clustered with the Seurat *FindClusters* function with resolution=0.1 and plotted using UMAP dimensionality reduction (Fig. 1C). In addition, a UMAP plot with the same clustering parameters and coloring of cells by the sample was produced (Suppl. Fig. 2). 804 non-erythrocyte cluster cells were re-clustered separately with resolution=0.3 (Fig. 1D).

Marker genes were identified among clusters of 804 non-erythrocyte cluster cells with the Seurat *FindAllMarkers* function. *FindAllMarkers* was run 3 separate times with different parameters: (1. only.pos = TRUE, 2. only.pos = TRUE, assay = "RNA", 3. only.pos = TRUE, test.use = "MAST", assay = "RNA") top 20 marker genes for each cluster from each run of the function were taken, aggregated, duplicates removed and plotted on a heatmap (Suppl. Fig. 4). Percentages of cells with any expression of a marker gene in a cluster were used for generating the heatmap. Colors in the heatmap were generated with the row-normalized percentages matrix. Four best markers were chosen from each cluster and plotted on a separate heatmap (Fig. 1F). The average expression of characteristic markers for clusters was plotted on the UMAP dimensionality reduction plot using the Nebulosa package (Alquicira-Hernandez & Powell, 2021) *plot_density* function (Fig. 1E).

Highly variable genes (HVG) for Venn diagrams and subsequent analyses (Fig. 2B-D) were identified as the top 2,000 HVG produced by *FindVariableFeatures* function in the Seurat pipeline for each sample separately.

Functional enrichment analysis of 235 HVG specific to freshwater sticklebacks was performed with the clusterProfiler R package (Yu et al., 2012) using the Gene Ontology (GO) database (Ashburner et al., 2000, Carbon et al., 2020, Mi et al., 2018) (Fig. 2C). As a background, a union of all HVG identified for each sample was used. The adjusted p-value cutoff was set to 0.1. Additionally, the same analysis was performed for marine-specific HVG (Suppl. Fig. 6) as well as functional enrichment analysis using the KEGG database (Kanehisa et al., 2020), both for freshwater- and marine-specific HVG (Suppl. Fig. 7, 8).

F-ratio was calculated per gene as a ratio of expression variance among marine stickleback cells to the expression variance among freshwater stickleback cells. Prior to the F-ratio calculation, we performed normalization of the number of cells per sample. First, a sample with the minimal number of cells was identified. Next, for other samples, a downsampling procedure was performed, which randomly selected the number of cells in each sample equal to the sample with the minimal number of cells. The downsampling was done by a random sampling without replacement. Genes for F-ratio calculation were chosen as a union between 2,000 HVG identified per each sample (N=4,832). The resulting F-ratios were plotted as a histogram (Fig. 2D). Additionally, the same procedure was redone for all genes from all cells in the dataset (Suppl. Fig. 5). Functional enrichment analysis using the GO and KEGG databases was performed for 100 top and bottom genes ordered by their F-ratio (Suppl. Fig. 9, 10). GO enrichment produced no terms that passed the significance threshold.

### scATAC-seq data processing

A total of 444,039,620 paired-end sequencing reads from 4 samples of scATAC-seq were processed using publicly available 10x Genomics software Cell Ranger ATAC v2.0 (Satpathy et al., 2019). (Suppl. Table 6) The sparse open chromatin peaks matrix generated by the Cell Ranger ATAC analysis pipeline with the list of 42,569 cells was used as input to the Signac software v1.2.1 (Stuart et al., 2020). Signac pipeline standard quality control and cell filtration steps were performed for each sample individually (Suppl. Fig. 14-17); parameters for filtration for each sample are presented in the table (Suppl. Table 7).

Next, peaks from all four samples were merged following the Signac default “merging objects” procedure. For this, a unified set of peaks for all samples was created using the *GenomicRanges* package *reduce()* approach, which merges the overlapping peaks to form a single one. The resulting count matrix over a unified set of peaks was used for further differential variance analysis. There was no integration of samples (as was the case for scRNA-seq data) since tools for integration and batch correction perform poorly for scATAC-seq, especially with low cell count in some samples (Baek & Lee, 2020). To calculate differences in variance in open chromatin peaks in cells from marine and freshwater samples, the F-test was used. F-test was applied to each peak - a row in the count matrix from the previous step. Variances in cells from marine compared to freshwater samples (“marine cells” compared to “freshwater cells”) were tested. The resulting ratios of variances (F-ratios) for each peak between marine or freshwater cells were plotted as a histogram (Fig. 3D). The same procedure was performed for erythrocyte cells only (Suppl. Fig. 27). Prior to the F-ratio calculation, we performed a downsampling procedure to equalize the number of cells, in exactly the same way as for scRNA-seq dataset (see above).

F-ratio for divergence islands was calculated in the following way: the number of peaks with non-zero counts overlapping the DI was calculated for each DI for each cell to create a DIs/cells matrix. Marine and freshwater cells were downsampled to a random subset of 6,000 cells each to equalize the number of cells. Next, the F-ratio per DI was calculated as the ratio of variances between marine and freshwater cells in this matrix. Downsampling and F-ratio calculation were repeated 1,000 times. Mean F-ratios per DI from the 1,000 bootstraps were presented as a box-plot (Suppl. Fig. 31). The same procedure was repeated for the erythrocyte cells alone (Suppl. Fig. 30).

Functional enrichment analysis based on the Gene Ontology database (Yu et al., 2012) was performed on the sets of top variable genes in cells from samples in each salinity condition. To estimate gene variance between cells from the peaks count matrix, the following procedure was used. For each gene, two metrics were calculated: (1) the number of peaks with non-zero counts overlapping the gene and (2) mean counts in peaks overlapping the gene. Using these two metrics, two genes/cells matrices were calculated from the original peaks/cells count matrix. The same F-test procedure was performed for each of the genes/cells matrices, as for the peaks/cells one. The result was two lists of genes with corresponding F-ratio values. From these lists, two sets of genes with high variance in marine cells (F-ratio > 2) and two sets of genes with high variance in freshwater cells (F-ratio < 0.7) were calculated. Next, for each salinity condition, the two sets of HVG were intersected. That produced two sets of HVG, 254 genes in marine and 254 in freshwater fish, which resulted from intersecting calculations based on the metrics (1) and (2) described above. These two sets of condition-specific HVG were converted to *Danio rerio* orthologs and provided as an input to the GO enrichment analysis using the clusterProfiler package with the adjusted p-value threshold of 0.1. The resulting top 30 GO terms for marine-specific (Fig. 3E) and freshwater-specific (Suppl. Fig. 26) HVG were selected. The same procedure was repeated for the erythrocyte cells alone (Suppl. Fig. 28, 29).

Blood clusters were identified for each sample as clusters with the high estimated activity of genes: *HBE1*, *cpox*, *snx3*, and open chromatin peaks in the HBE1 region. Gene activities were calculated from the pattern of open chromatin from scATAC-seq data using the *GeneActivity()* Signac function. Estimated gene activities were plotted as violin and average expression UMAP plots using Seurat *Vln_plot()* and *Dim_plot()* functions. HBE1 open chromatin peaks were plotted using Signac *CoveragePlot()* function (region = "groupXI:13662623-13663375") (Suppl. Fig. 21-24).

Cells from the first marine sample were clustered into 10 distinct clusters. Gene activity was calculated from the pattern of open chromatin from scATAC-seq data using the GeneActivity() Signac function. Cells were annotated using labels of non-erythrocyte cells from scRNA-seq analysis. Labels were assigned on a cell-by-cell basis using *FindTransferAnchors()* Signac function. The first marine sample without erythrocyte clusters was used for direct label transfer (Suppl. Fig. 25). Next, clusters based on ATAC-seq data were annotated with consistent labels and refined using cell markers, the same as for scRNA-seq. The resulting labels were plotted with UMAP dimensionality reduction (Fig. 3B).

### Bisulfite conversion and whole-genome bisulfite sequencing

1 mkg of Stickleback genomic DNA was mixed with 10 ng lambda phage DNA and sheared with ultrasound to the average size of 300 bp. End-repair, dA tailing, and methylated adaptor ligation were performed with NebNext DNA UltraII kit (NEB). After adaptor ligation, libraries were bisulfite converted with EZ DNA Methylation Kits (ZYMO RESEARCH) according to the manufacturer’s protocol. After conversion, final libraries were amplified with NEBNext Q5U® Master Mix (NEB) and sequenced with Illumina HiSeq1500.

### WGBS data processing

Paired-end reads were processed with Trim Galore ver. 0.5.0 (Krueger et al., 2021) to remove adapter sequences and trim bases with low quality scores (<20). Validated reads were aligned to Broad/gasAcu1 genome assembly with Bismark software (Krueger & Andrews, 2011). Most of the CpGs were covered by at least 10 reads in all samples. Bisulfite conversion efficiency (> 99%) was assessed using both lambda phage and methylation of non-CpG context. Aligned reads were randomly downsampled to achieve an uniform number across all samples (42mln).

The methylation entropy for five-CpG bins was calculated using an approach implemented in the *MethPipe* pipeline (Song et al., 2013). For each sliding window, the frequencies of methylation patterns were calculated, then the products were summed up according to the formula:

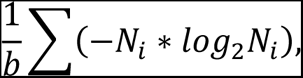

where *b* is the number of CpG sites and *Ni* is the frequency of methylation pattern *i*.

To select five-CpG bins that have different entropy, we performed the U-test followed by the Benjamini-Hochberg procedure. We used an average value across all samples belonging to the same group to plot entropy distributions (near TSS, in gene bodies, etc.).

To verify the robustness of our method for assessing epigenetic heterogeneity at different levels of coverage, we merged all reads by sample groups (Freshwater-Males, Marine-Males, Freshwater-Females, Marine-Females), performed downsampling, and calculated methylation entropy in the way described above.

Data for constructing correlations were obtained in two ways. In the first case, we took a random sample of 1000 peaks and averaged the entropy values for the selected peaks. In the second case, we did similar steps, but instead of absolute values, we averaged the log2FCs between freshwater and marine fish.

We used *splsda()* function of mixOmics package (Rohart et al., 2017) for feature selection purpose. We also determined the optimal peaks number using the function *tune.splsda()* with 5-fold cross-validation and 50 repeats.We used CrossMap software (Zhao et al., 2014) to liftover coordinates between gasAcu1-4 and gasAcu1 assemblies.

### WGS data processing

Terekhanova et al. sequenced and pulled non-bisulfite-converted reads of marine and freshwater fish (Terekhanova et al., 2019). We processed, aligned the reads and calculated pseudo-methylation entropy exactly the same way as for WGBS dataset. Then we used the obtained entropy values as a normalization factor.

## DATA AND CODE AVAILABILITY

The R code performing the main steps of data analysis described in this paper is freely available at GitHub: https://github.com/artgolden/stickleback_paper. The raw sequencing reads are deposited at SRA under the BioProject PRJNA765182.

## ACKNOWLEDGMENTS

This study was supported by the Russian Science Foundation grant 19-14-00347, 19-74-30026 and the fieldwork for NM was supported by RFBR grant 20-04-01139. We are grateful to Dr. Ilya Kurochkin for helpful discussions on scRNA-seq analysis. The calculations were done using computational resources of the federal collective usage center MCC NRC “Kurchatov Institute”, http://computing.nrcki.ru/.

## AUTHOR CONTRIBUTIONS

E.P. and N.M. conceived this study. A.M., N.M., and D.K. performed the experiments. A.G., A.S., A.A., E.K., and E.P. analyzed the data. E.K., A.G., A.S., and E.P. prepared the figures. All authors contributed to writing the manuscript.

