## Supplemental Materials for "Epigenetic variations are accompanying landmarks of freshwater adaptation in threespine sticklebacks"

### **SUPPLEMENTARY FIGURES**

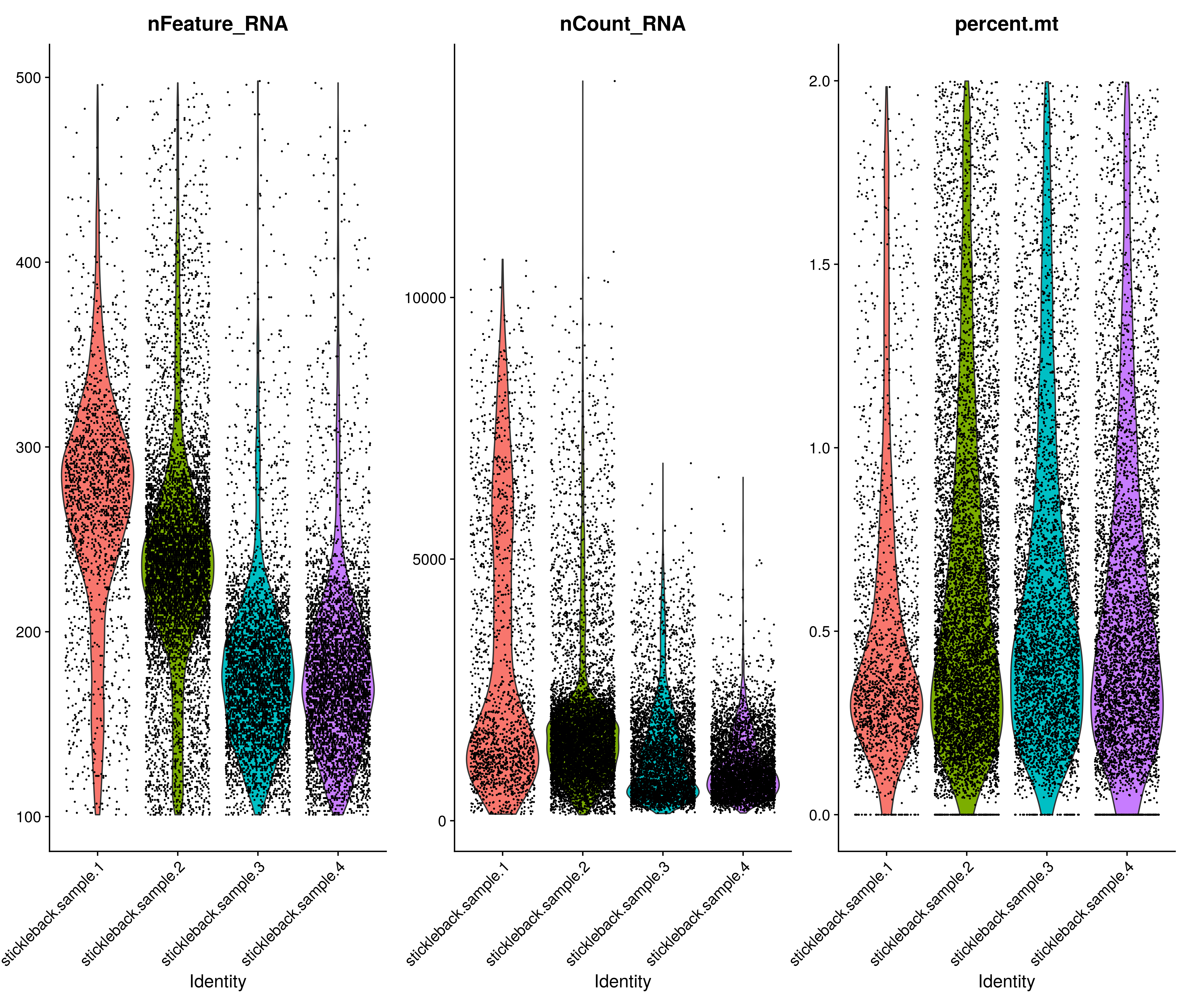

**Suppl. Fig. 1.** Quality control metrics for scRNA-seq data reported by the Seurat pipeline. Groups represent samples. Filtering thresholds: nFeature_RNA > 100, percent.mt < 2.

| **A** | 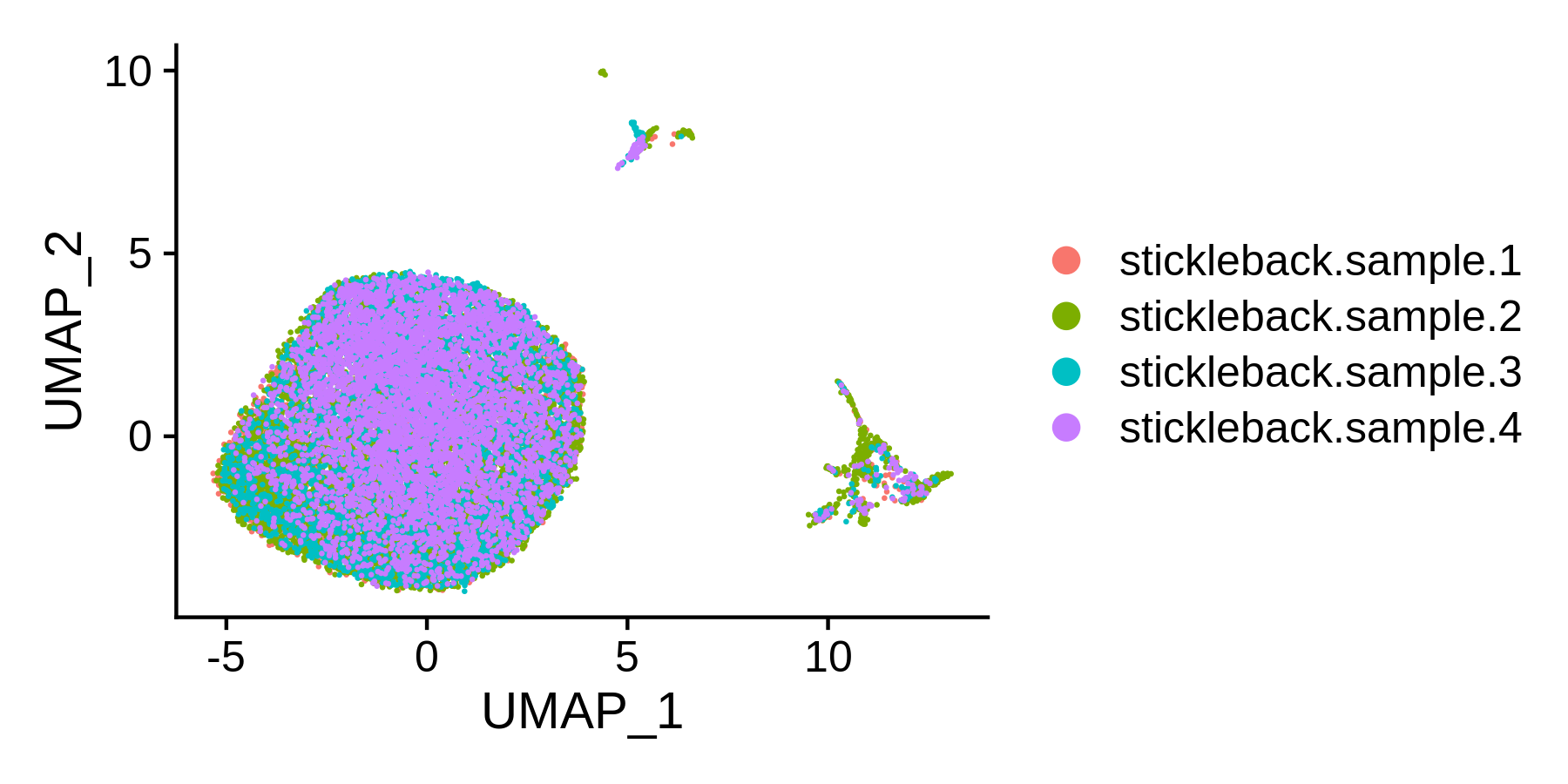 |
| --- | --- |
| **B** | 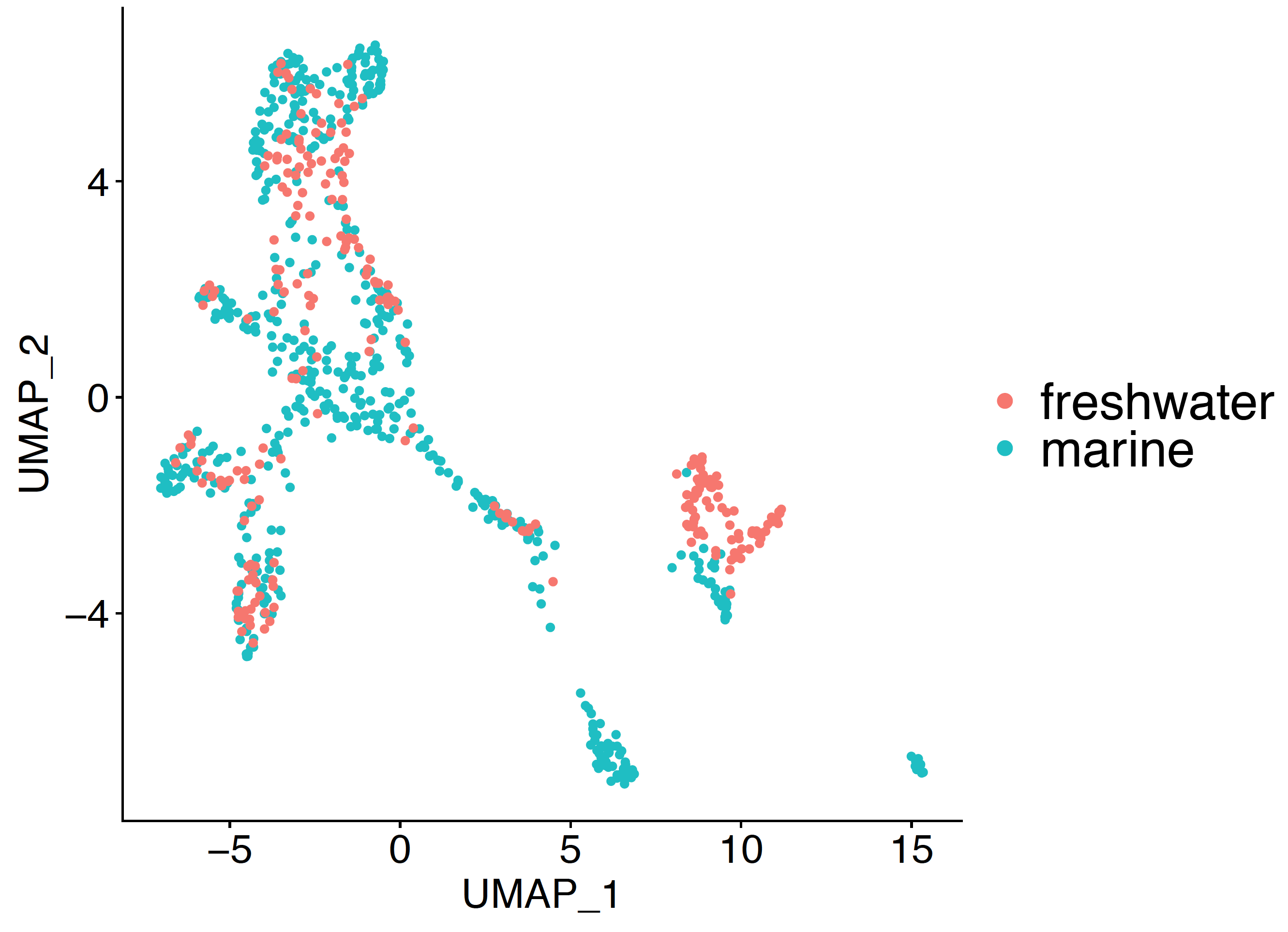 |

**Suppl. Fig. 2.** (A) UMAP plot of 19,964 cells from four integrated samples colored by sample identity. (B) UMAP plot of 804 non-erythrocyte cells colored by sample identity: freshwater (red) or marine (green). Fisher's exact two-sided test p-value = 0.0005 for 10 cell-type clusters presented in Fig. 1D.

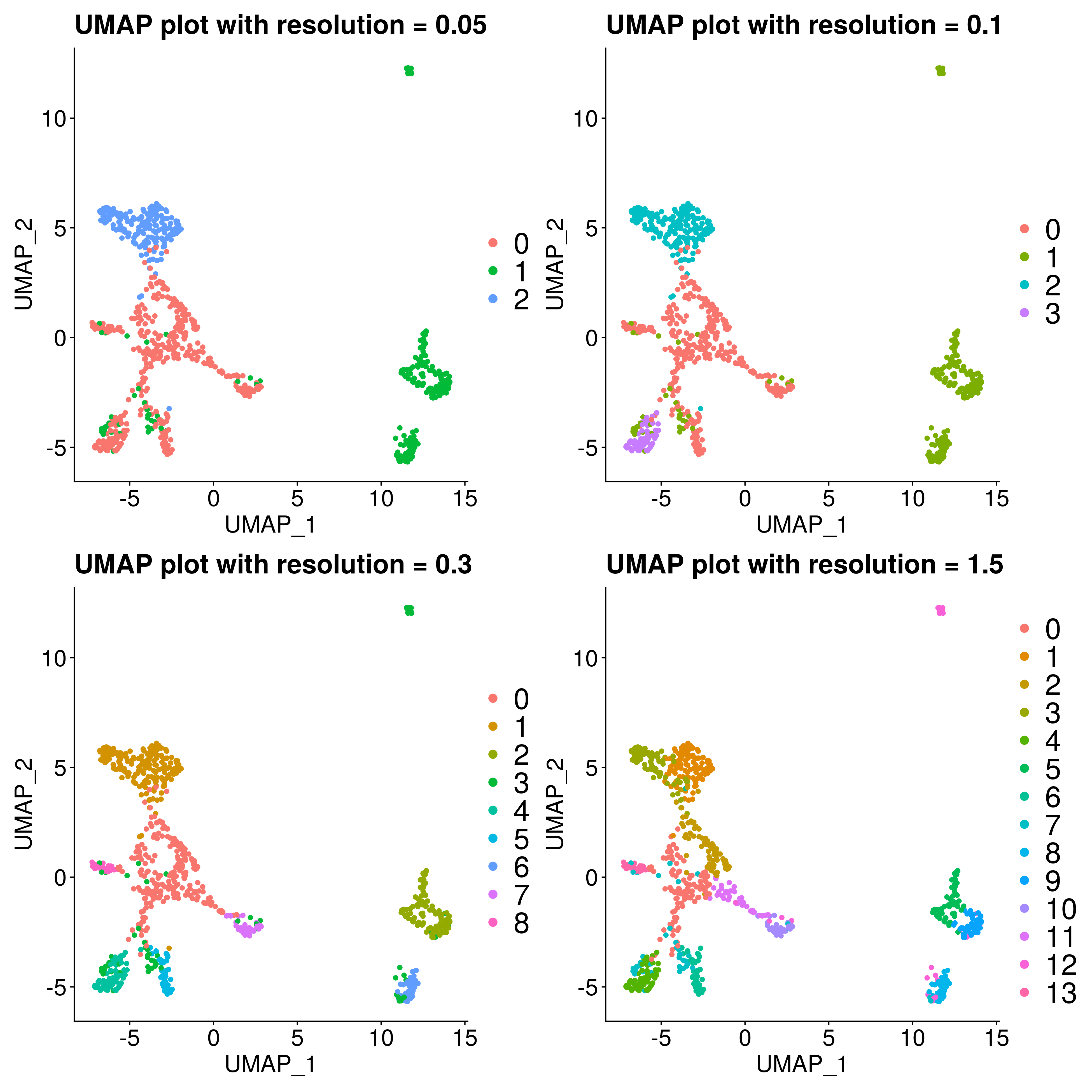

**Suppl. Fig. 3.** UMAP plots showing reclustering of 804 non-erythrocyte cells at different resolutions (0.05, 0.1, 0.3, and 1.5).

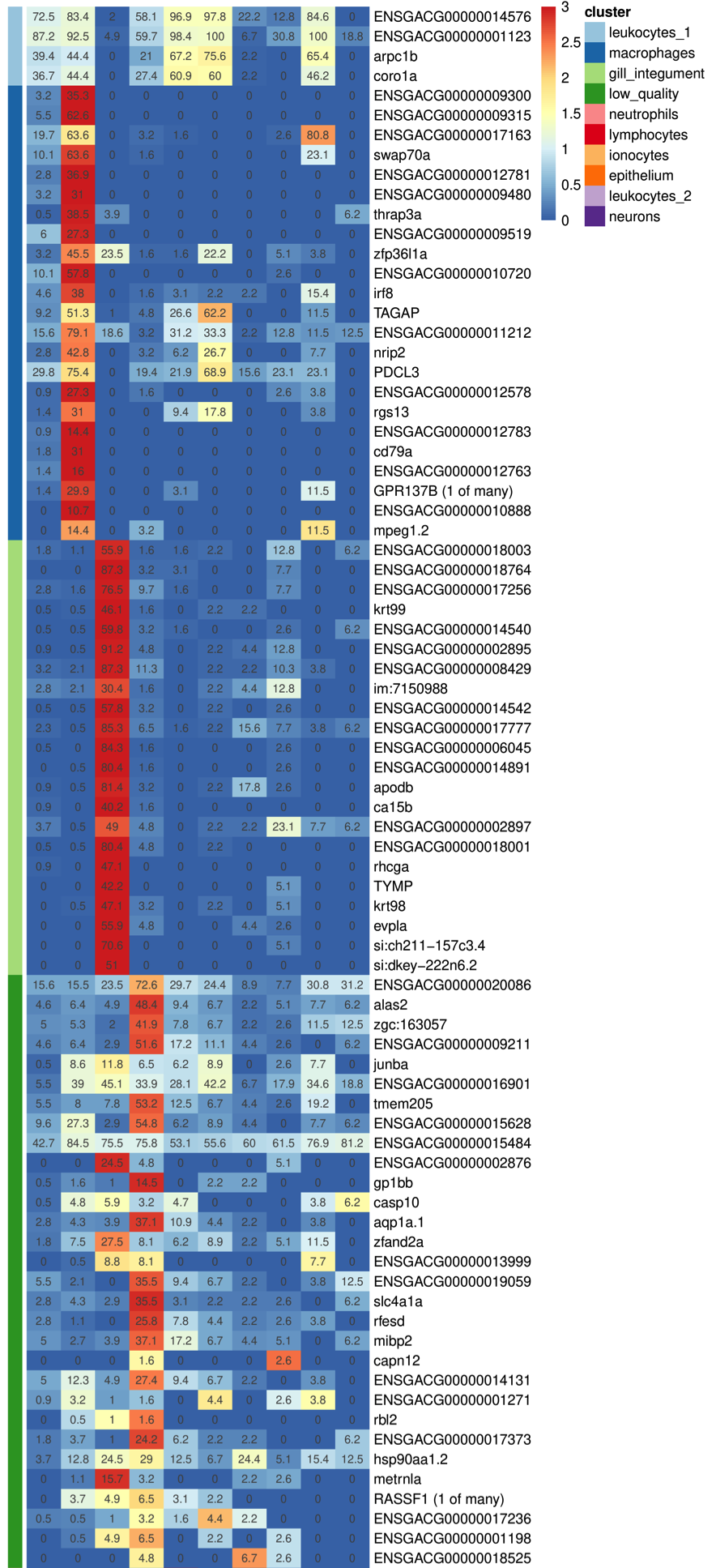

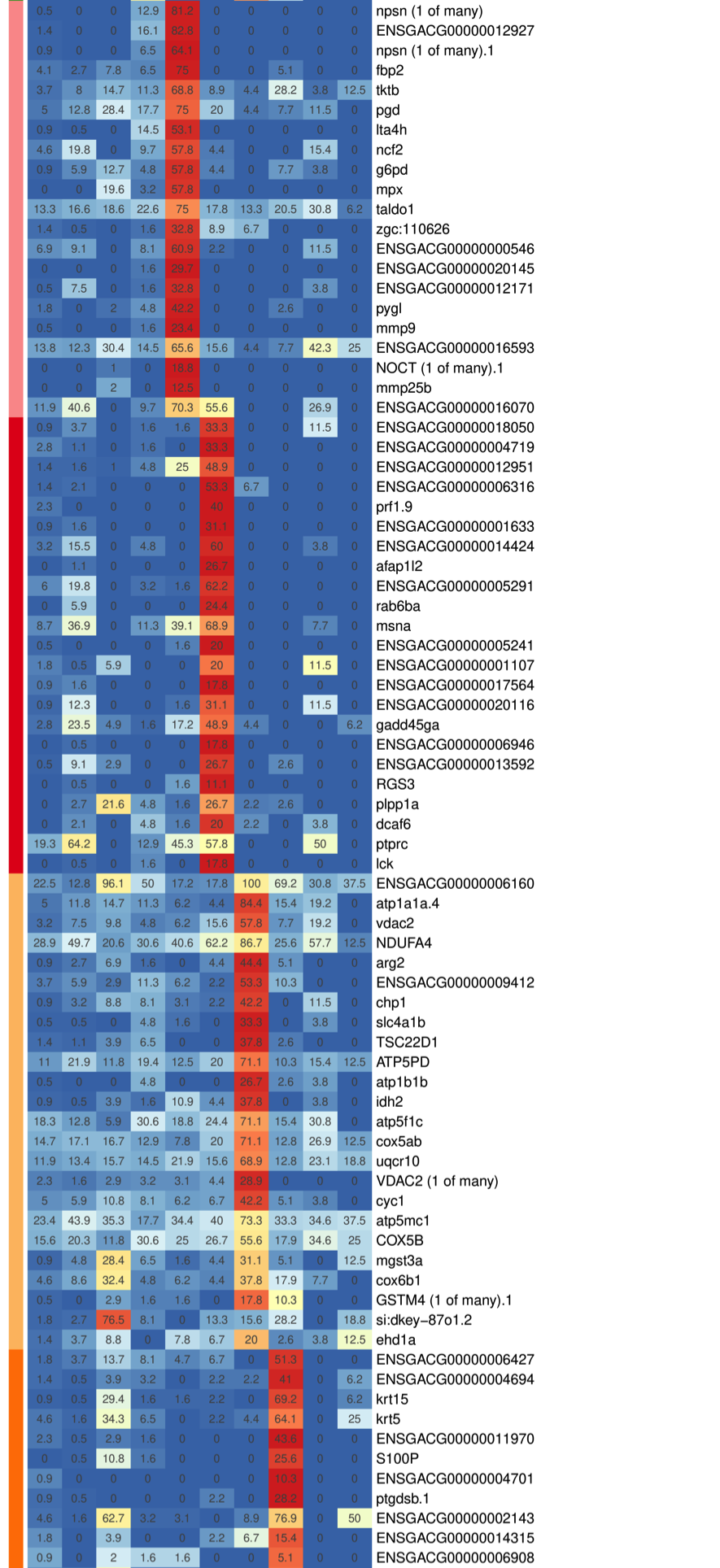

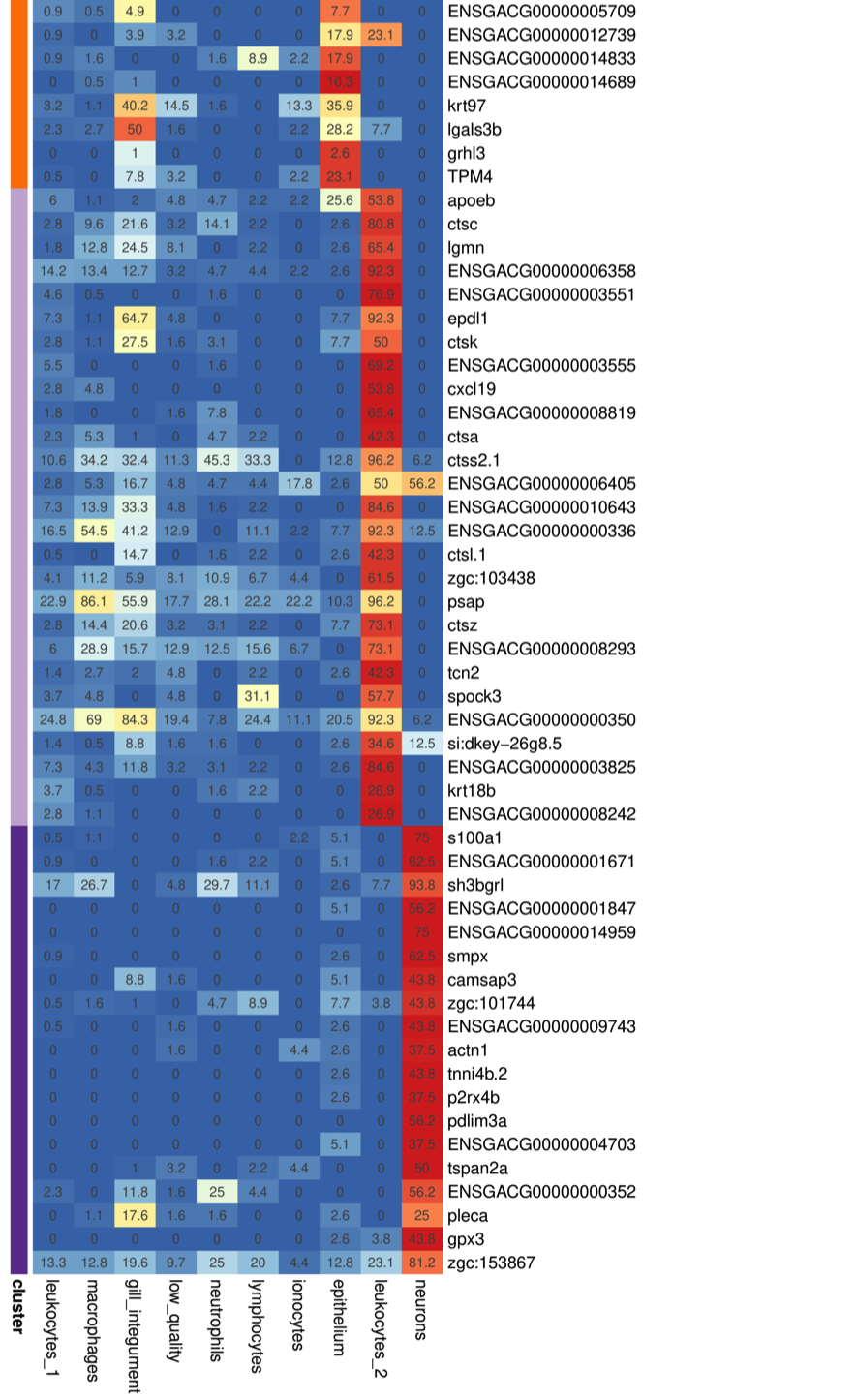

­

**Suppl. Fig. 4.** Heatmap of marker genes of non-erythrocyte clusters in the scRNA-seq dataset. Percentages of cells with any expression of a marker gene in a cluster were used for generating the heatmap. These percentages are also displayed inside the cells. Colors in the heatmap were generated with the row-normalized percentages matrix.

|  | 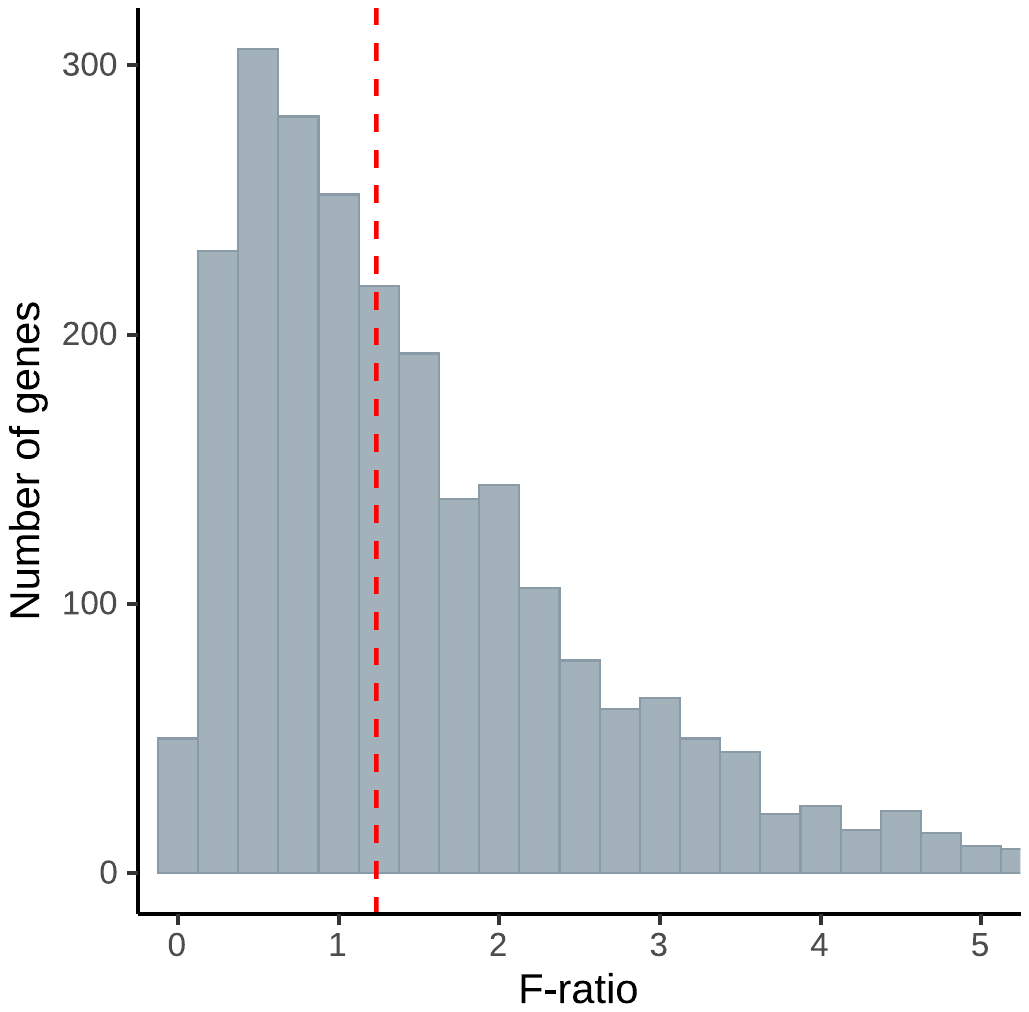 |
| --- | --- |

**Suppl. Fig. 5.** F-ratio (marine/freshwater) histogram calculated for all genes in the scRNA-seq dataset. All cells are analyzed.

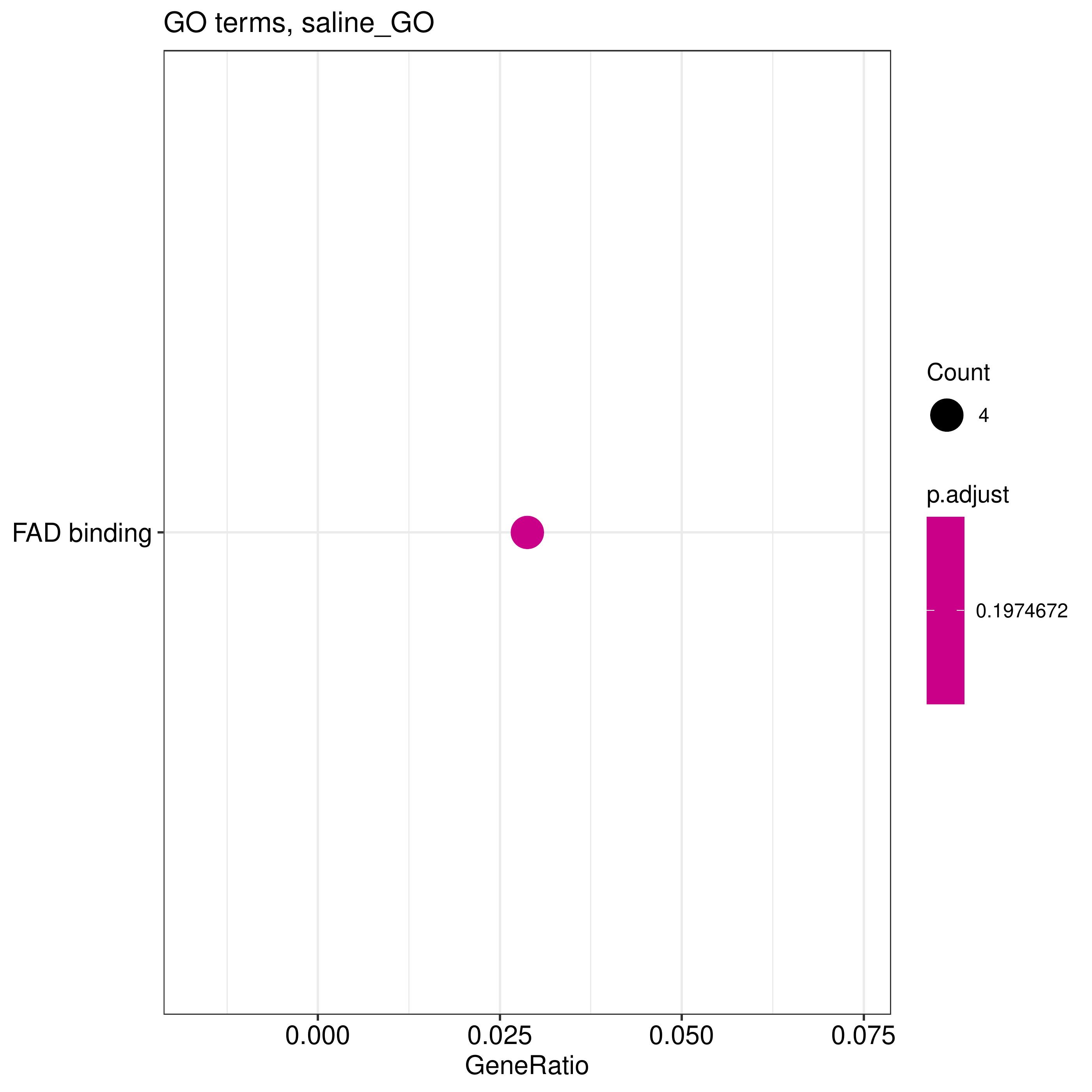

**Suppl. Fig. 6.** Functional enrichment analysis for HVG specific to marine sticklebacks. The analysis was performed using the clusterProfiler package based on the Gene Ontology (GO) database.

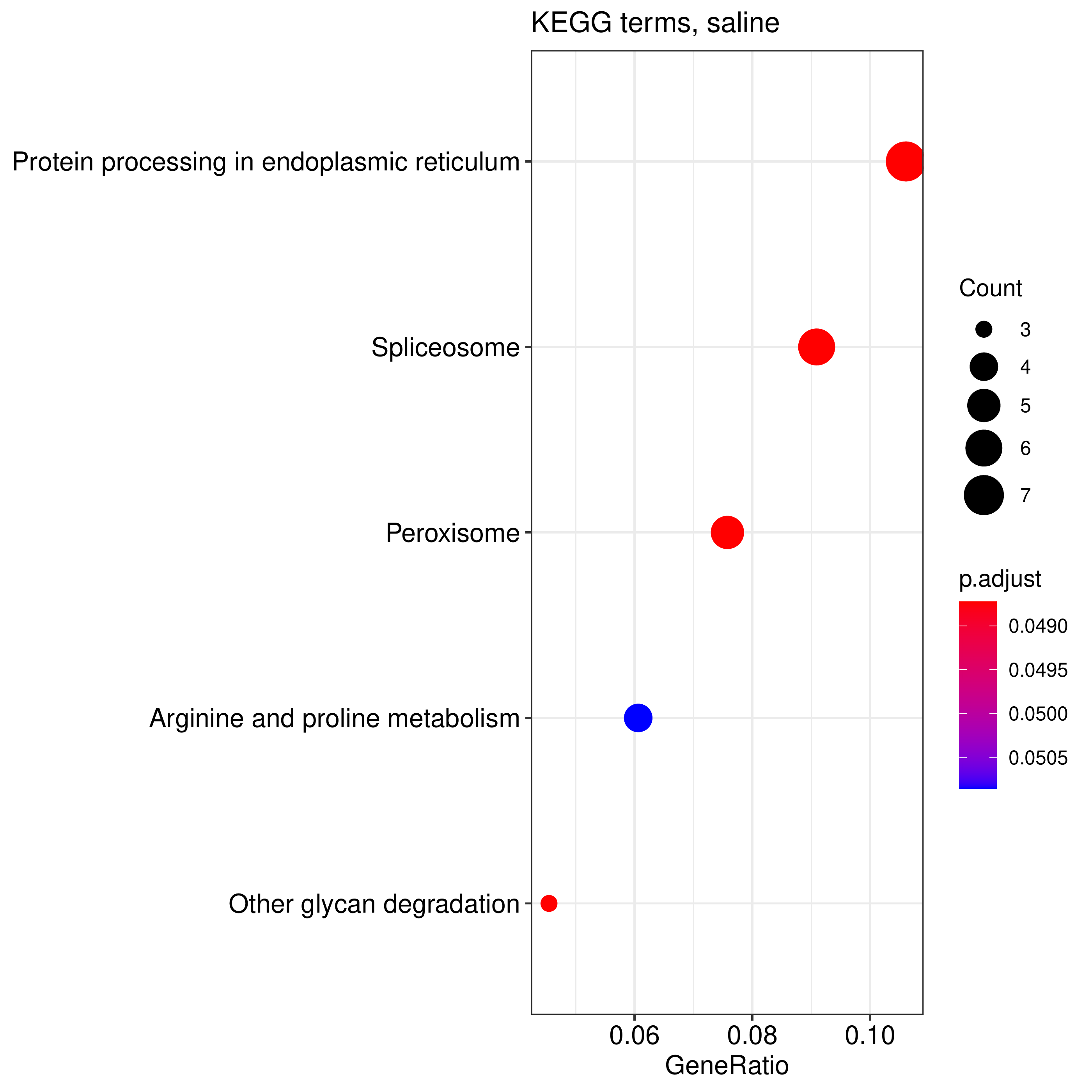

**Suppl. Fig. 7.** Functional enrichment analysis for HVG specific to marine sticklebacks. The analysis was performed using the clusterProfiler package based on the KEGG database.

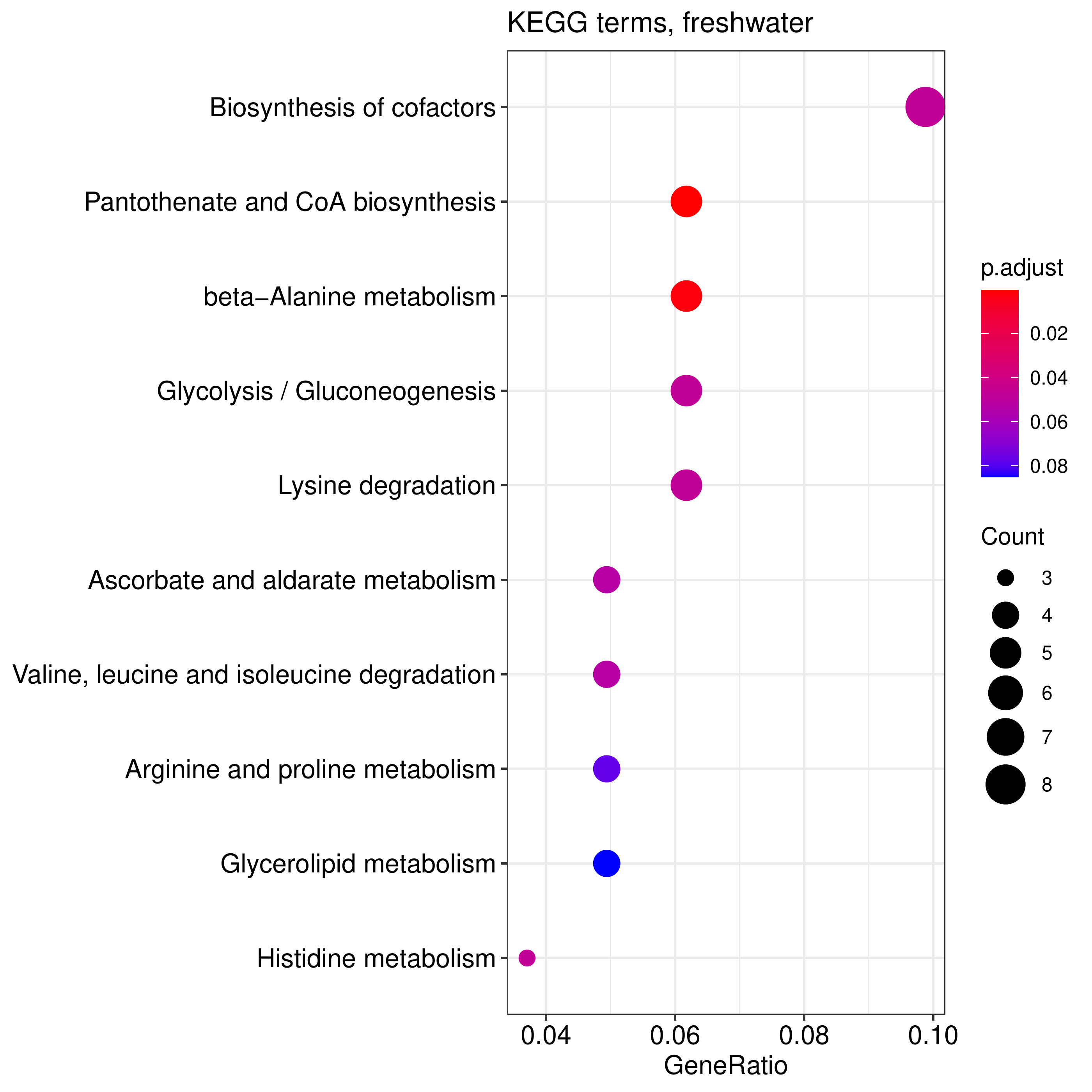

**Suppl. Fig. 8.** Functional enrichment analysis for HVG specific to freshwater sticklebacks. The analysis was performed using the clusterProfiler package based on the KEGG database.

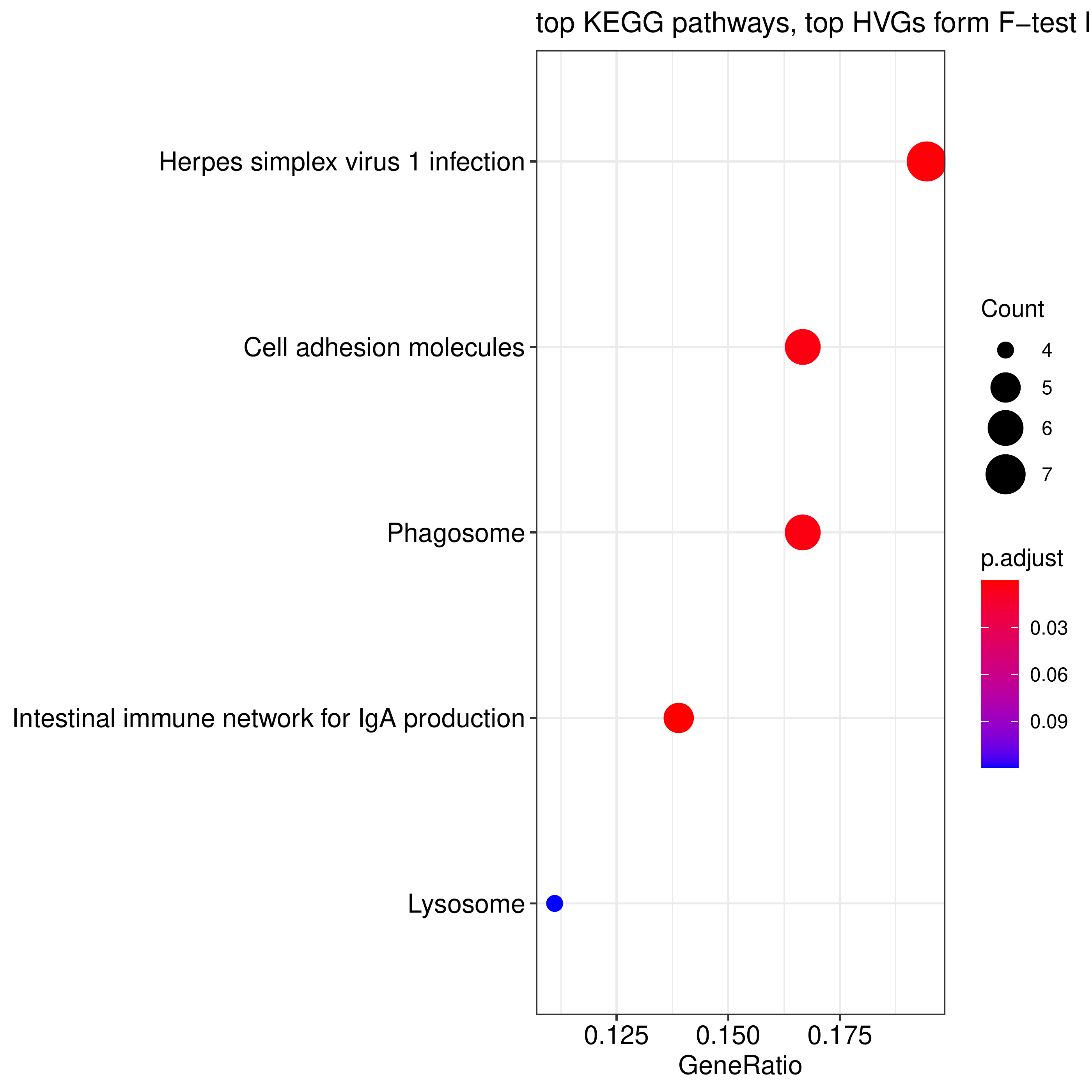

**Suppl. Fig. 9.** Functional enrichment analysis for 100 marine-specific HVG with the highest F-ratio values (marine/freshwater) for all cells from the erythrocyte cluster. The analysis was performed using the clusterProfiler package based on the KEGG database.

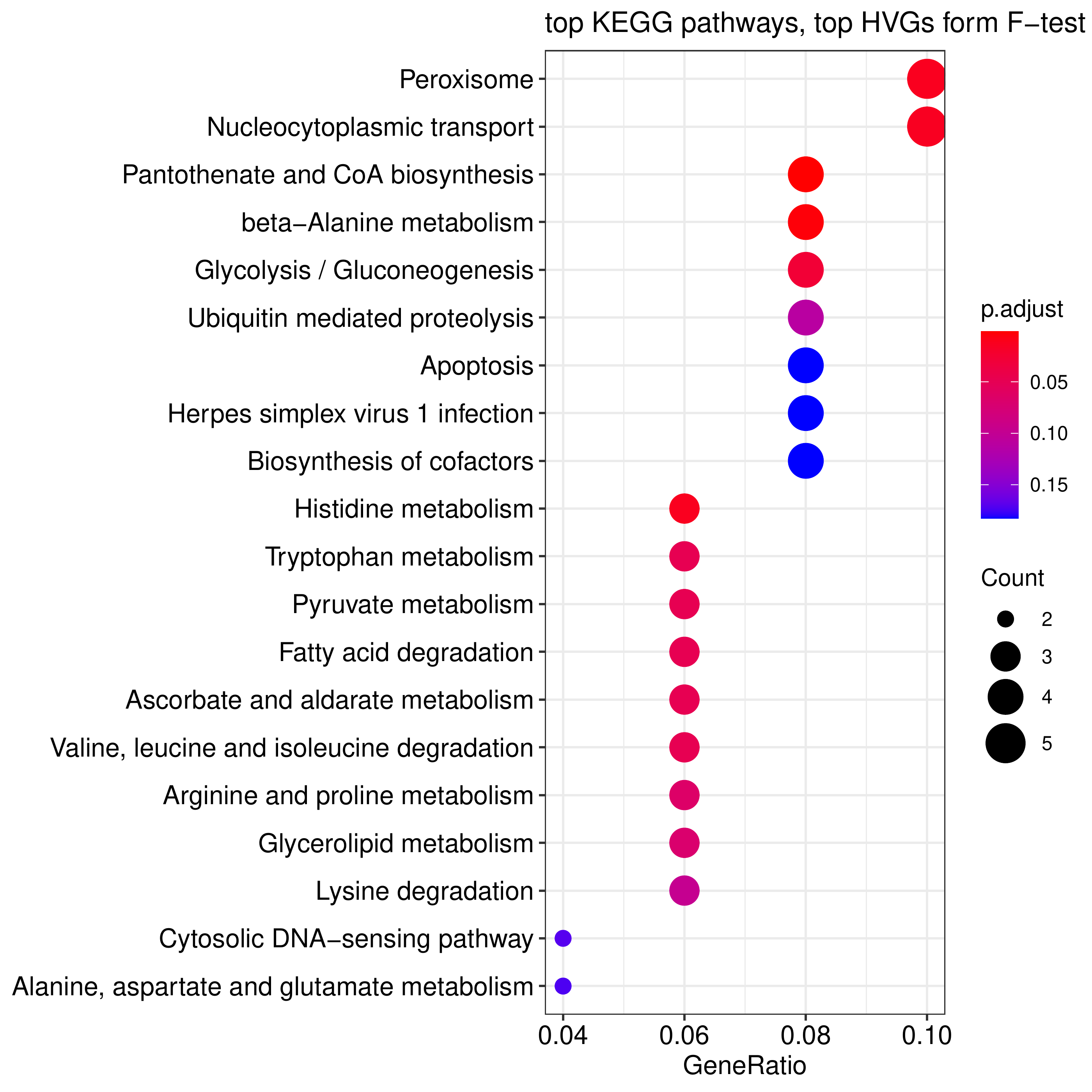

**Suppl. Fig. 10.** Functional enrichment analysis for lowest HVG defined by F-test ranking for all cells from the erythrocyte cluster. The analysis was performed using the clusterProfiler package based on the KEGG database.

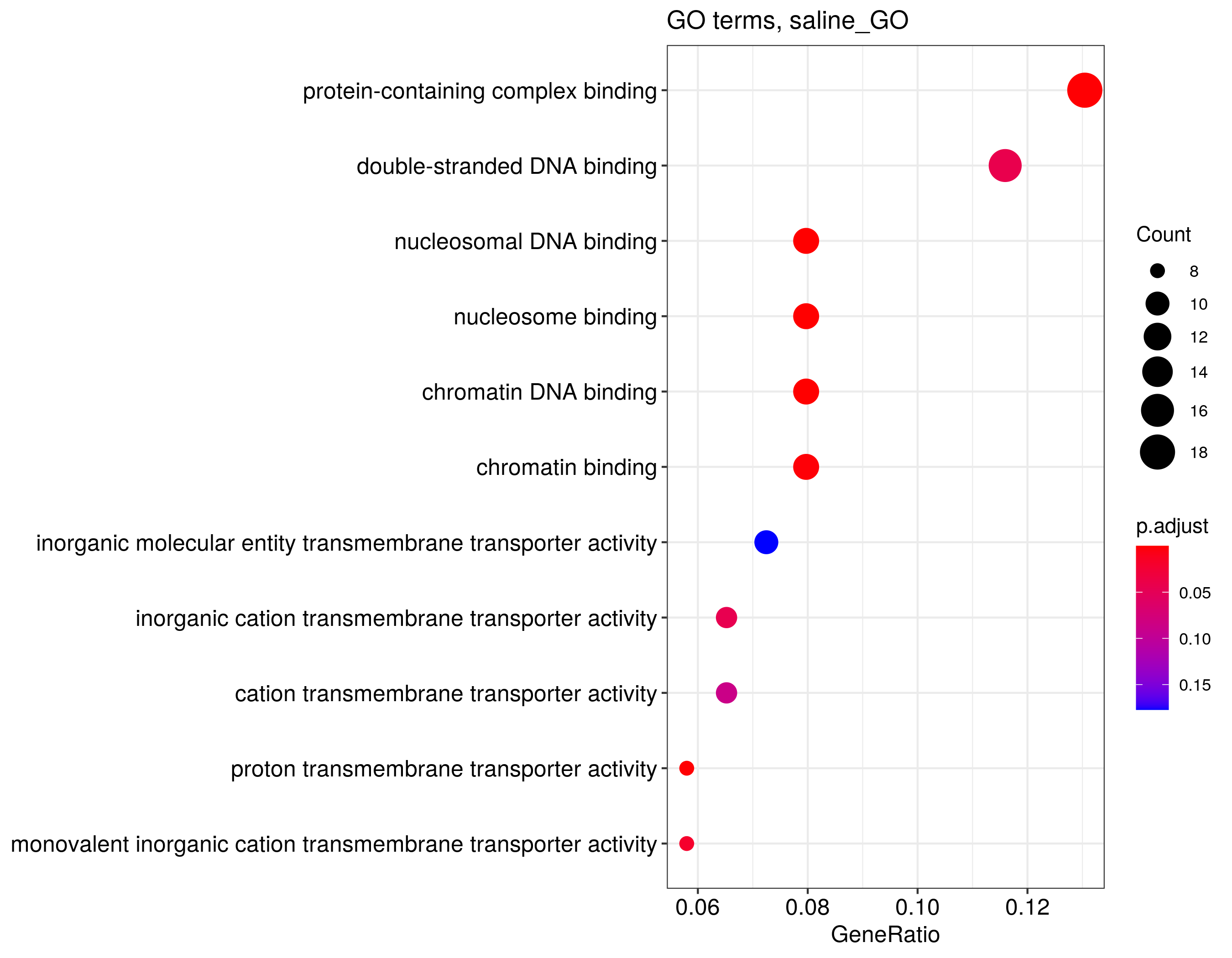

**Suppl. Fig. 11.** Functional enrichment analysis for HVG specific to marine sticklebacks for all cells in the scRNA-seq dataset. The analysis was performed using the clusterProfiler package based on the GO database.

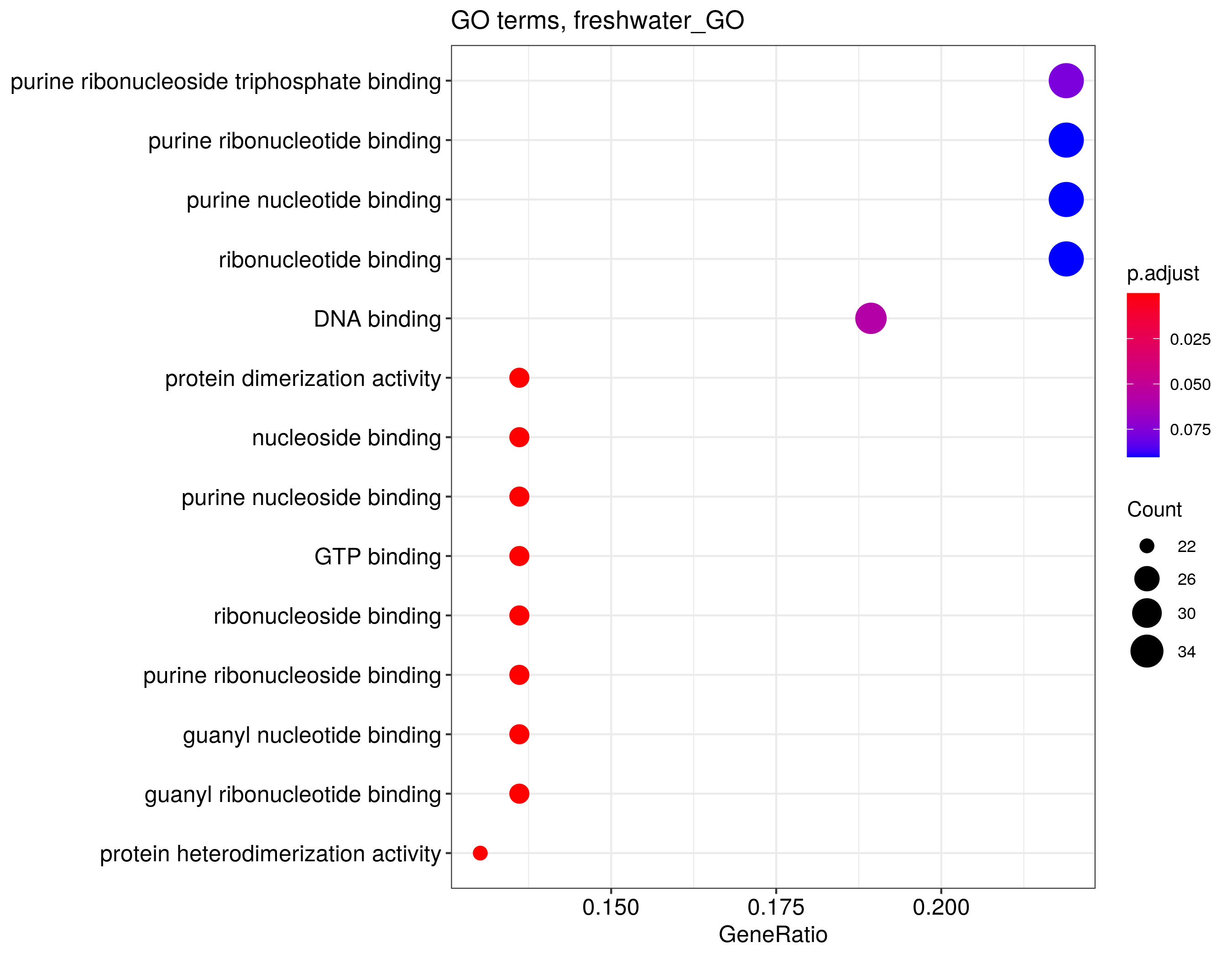

**Suppl. Fig. 12.** Functional enrichment analysis for HVG specific to freshwater sticklebacks for all cells in the scRNA-seq dataset. The analysis was performed using the clusterProfiler package based on the GO database.

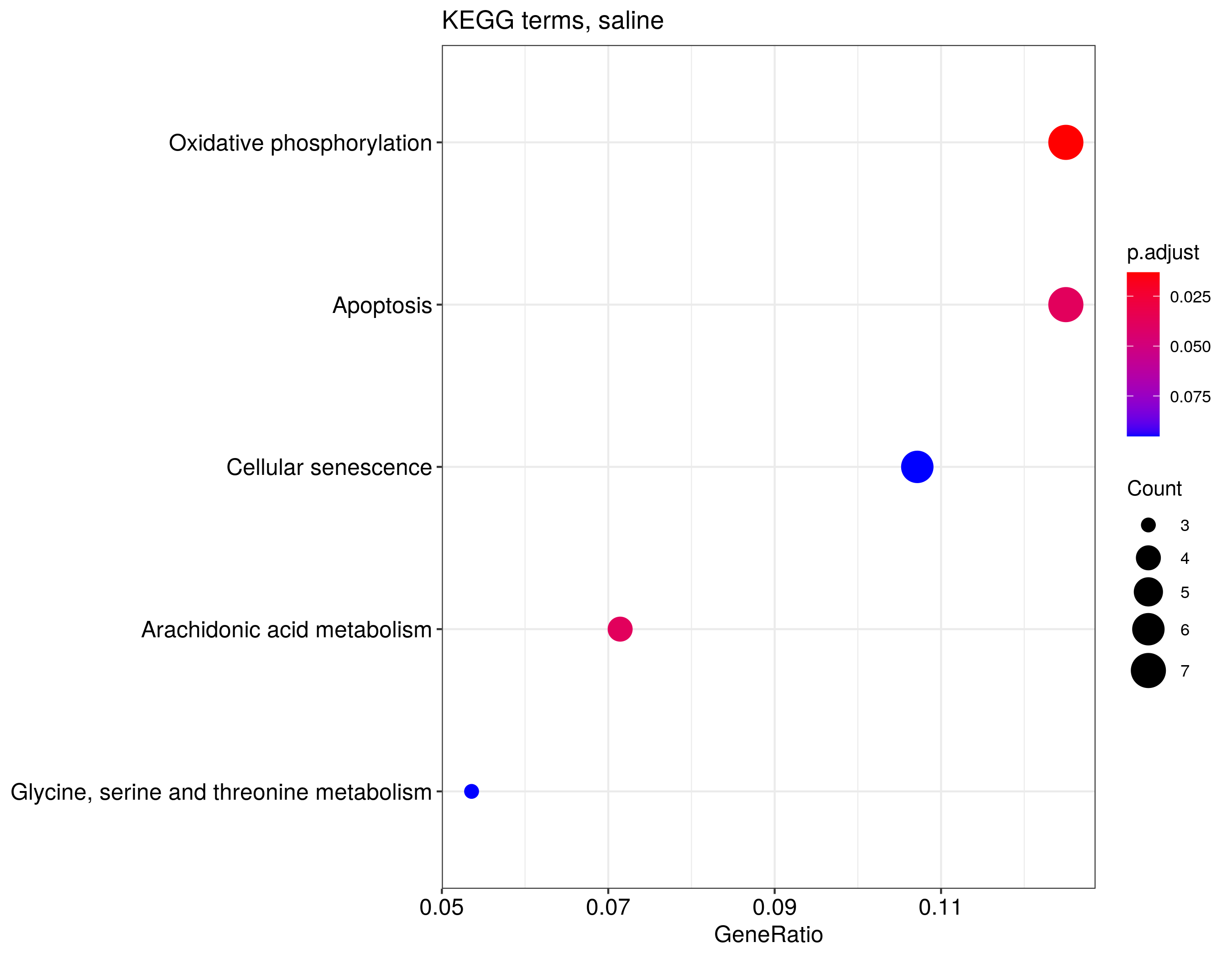

**Suppl. Fig. 13.** Functional enrichment analysis for HVG specific to marine sticklebacks for all cells in the scRNA-seq dataset. The analysis was performed using the clusterProfiler package based on the KEGG database.

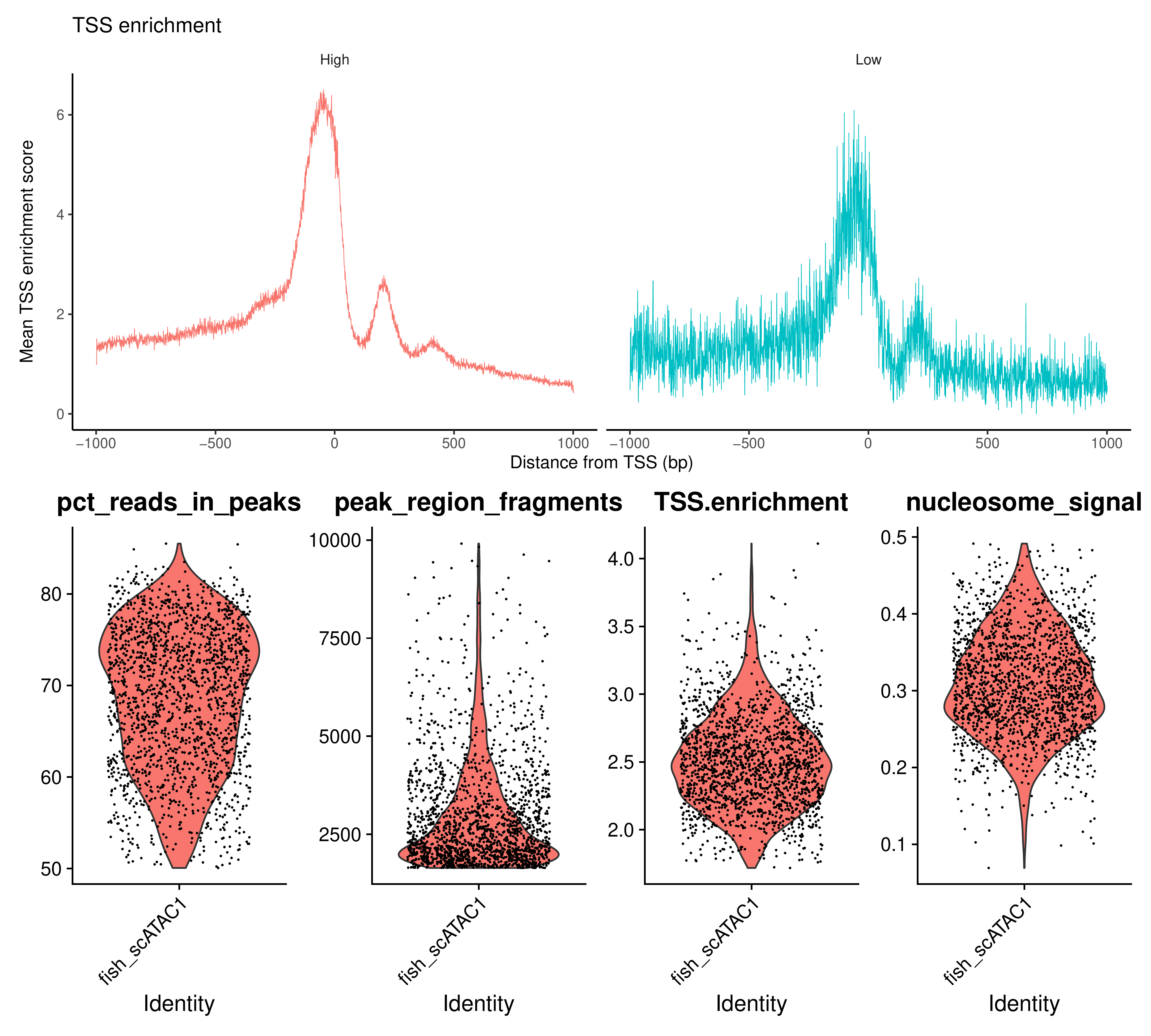

**Suppl. Fig. 14.** Quality control metrics produced by the Signac pipeline for the first marine stickleback sample. Threshold values are provided in Suppl. Table 7.

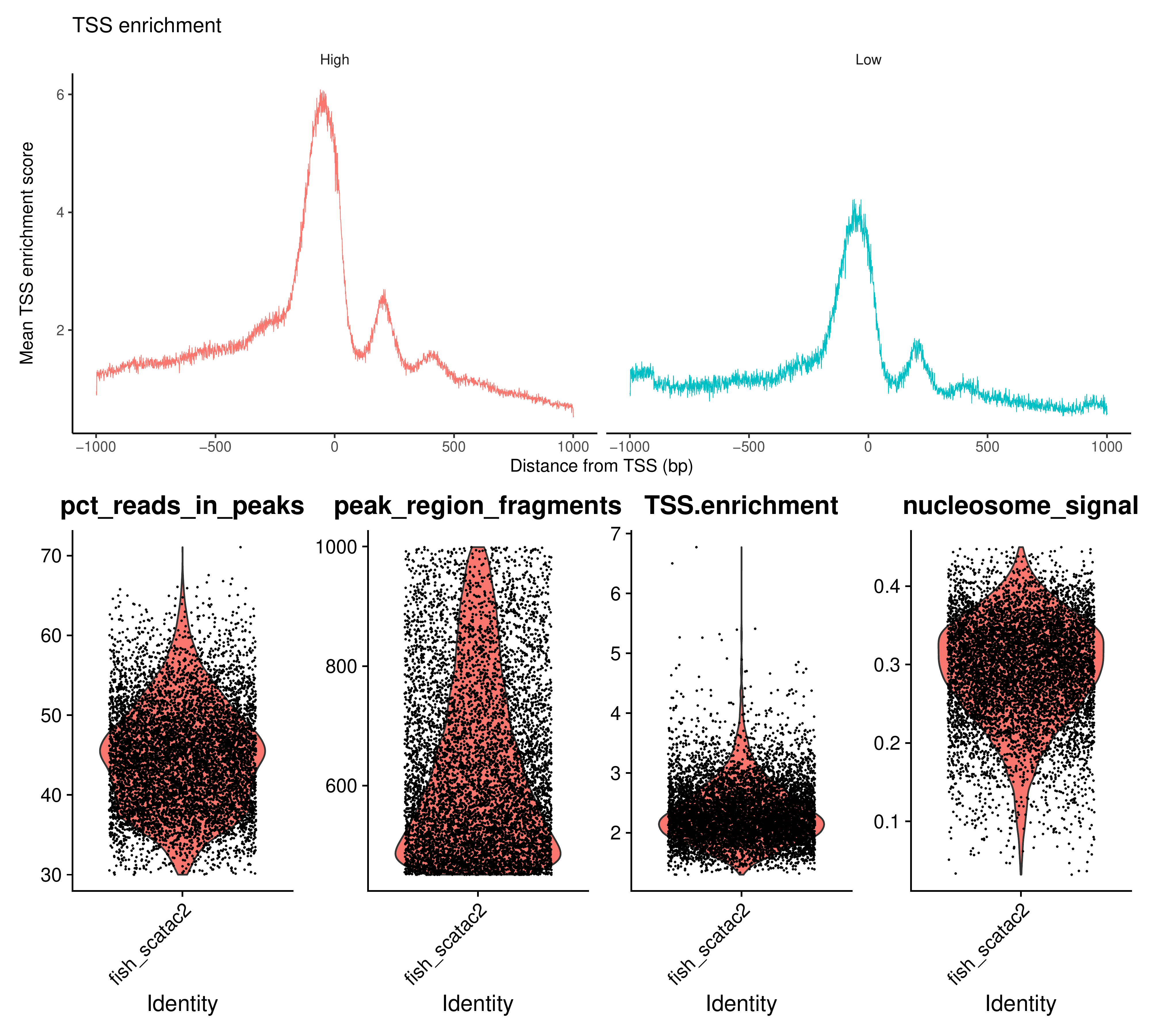

**Suppl. Fig. 15.** Quality control metrics produced by the Signac pipeline for the second marine stickleback sample. Threshold values are provided in Suppl. Table 7.

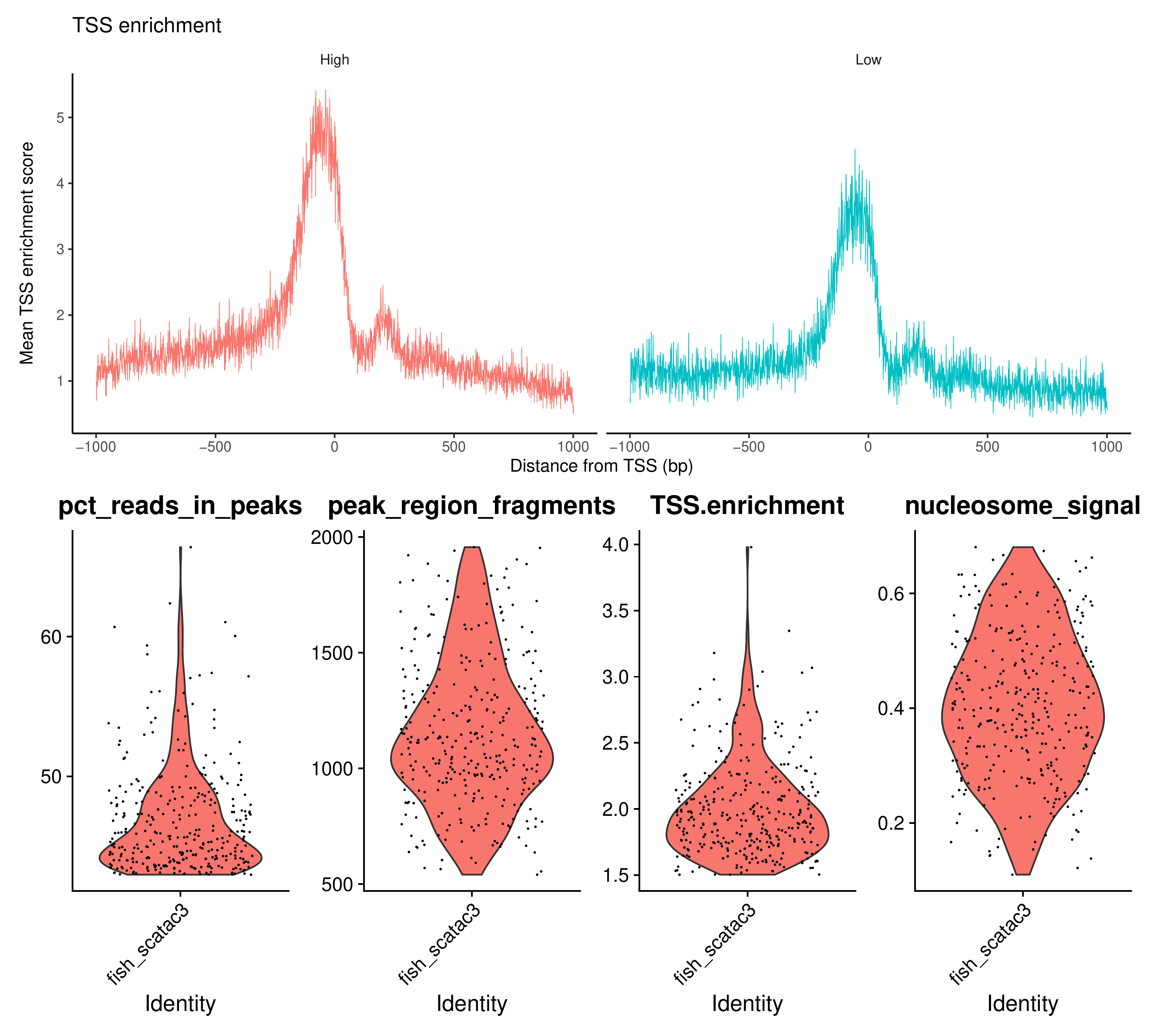

**Suppl. Fig. 16.** Quality control metrics produced by the Signac pipeline for the first freshwater stickleback sample. Threshold values are provided in Suppl. Table 7.

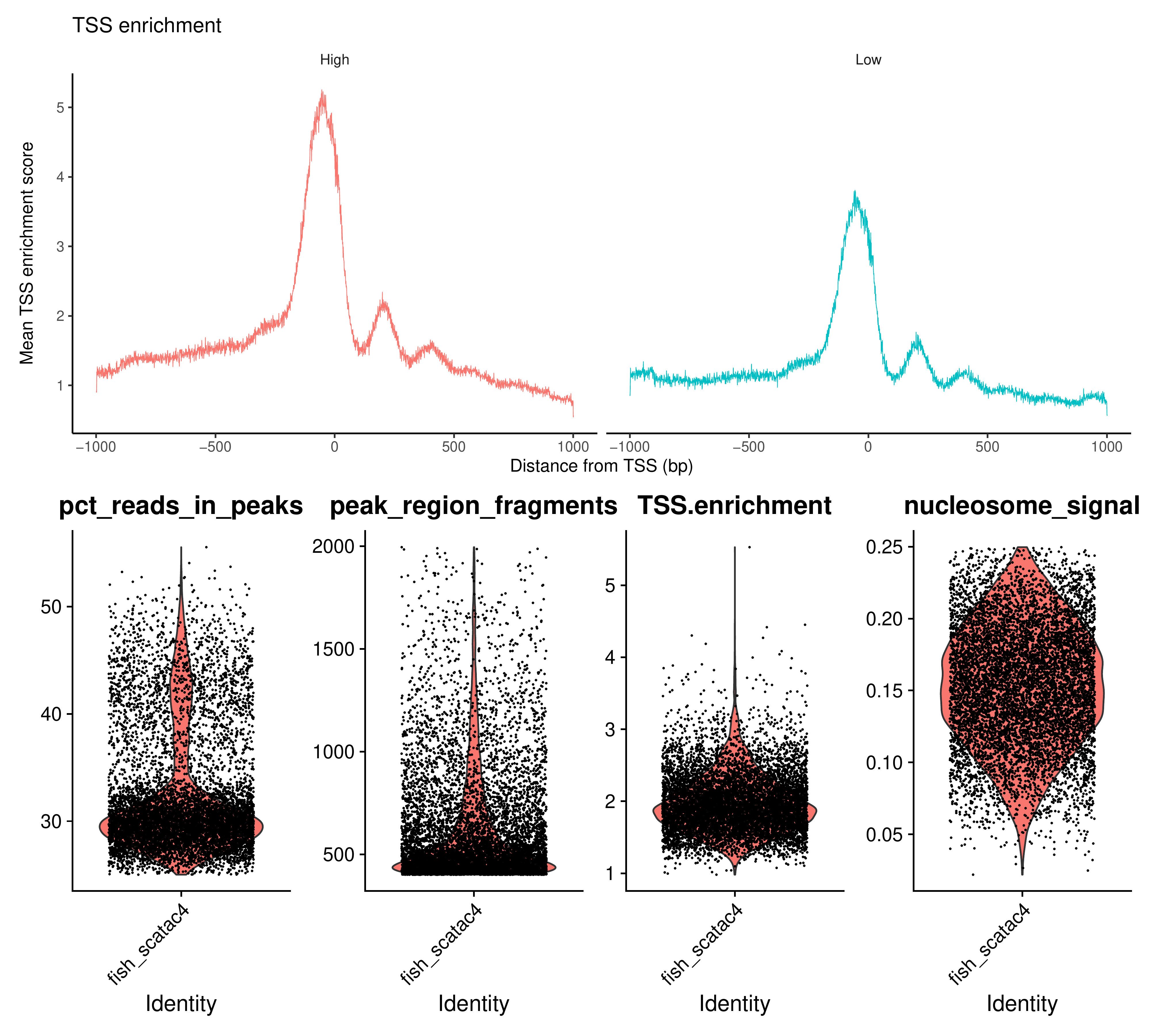

**Suppl. Fig. 17.** Quality control metrics produced by the Signac pipeline for the second freshwater stickleback sample. Threshold values are provided in Suppl. Table 7.

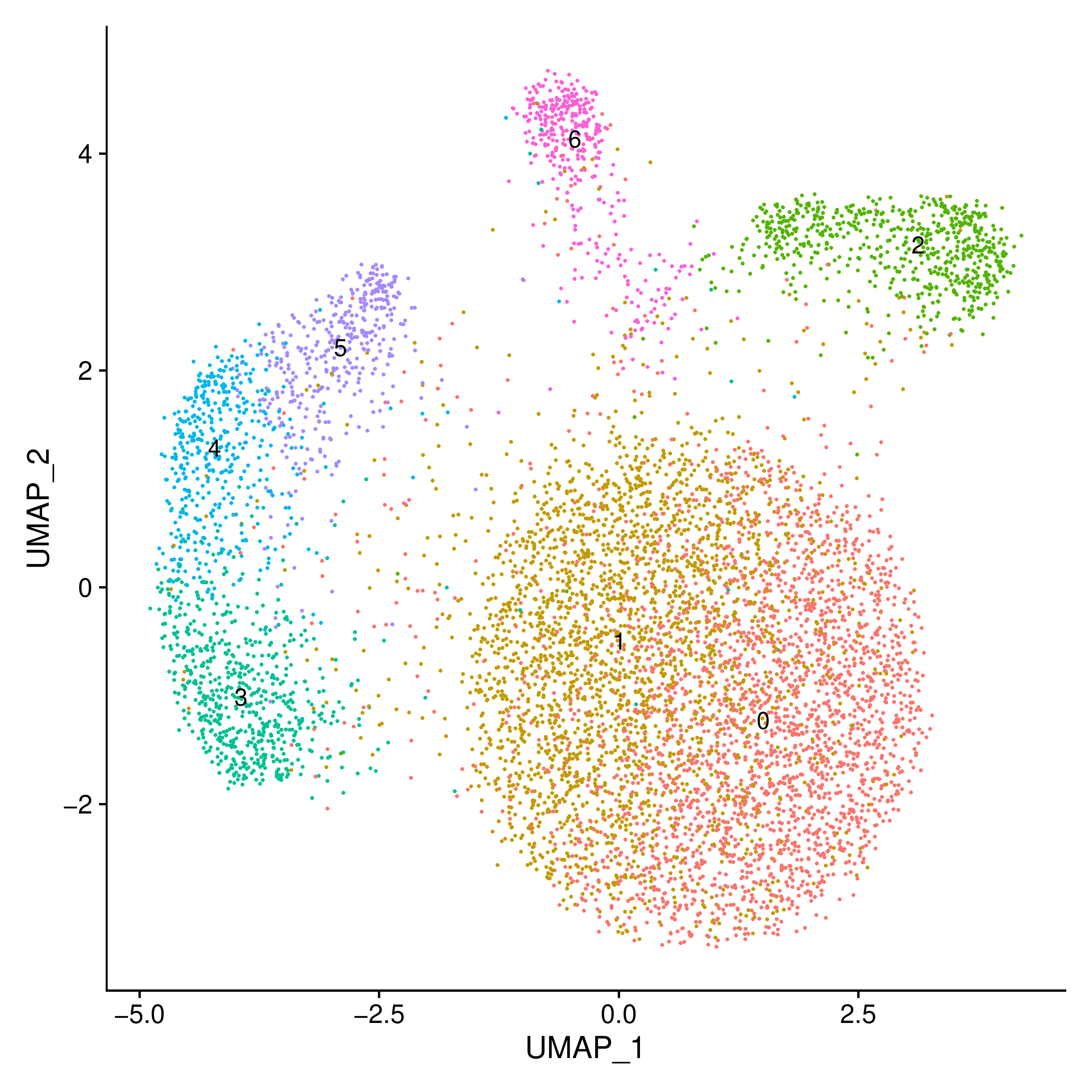

**Suppl. Fig. 18.** UMAP plot showing clustering of cells in the second marine sample, resolution = 1.2.

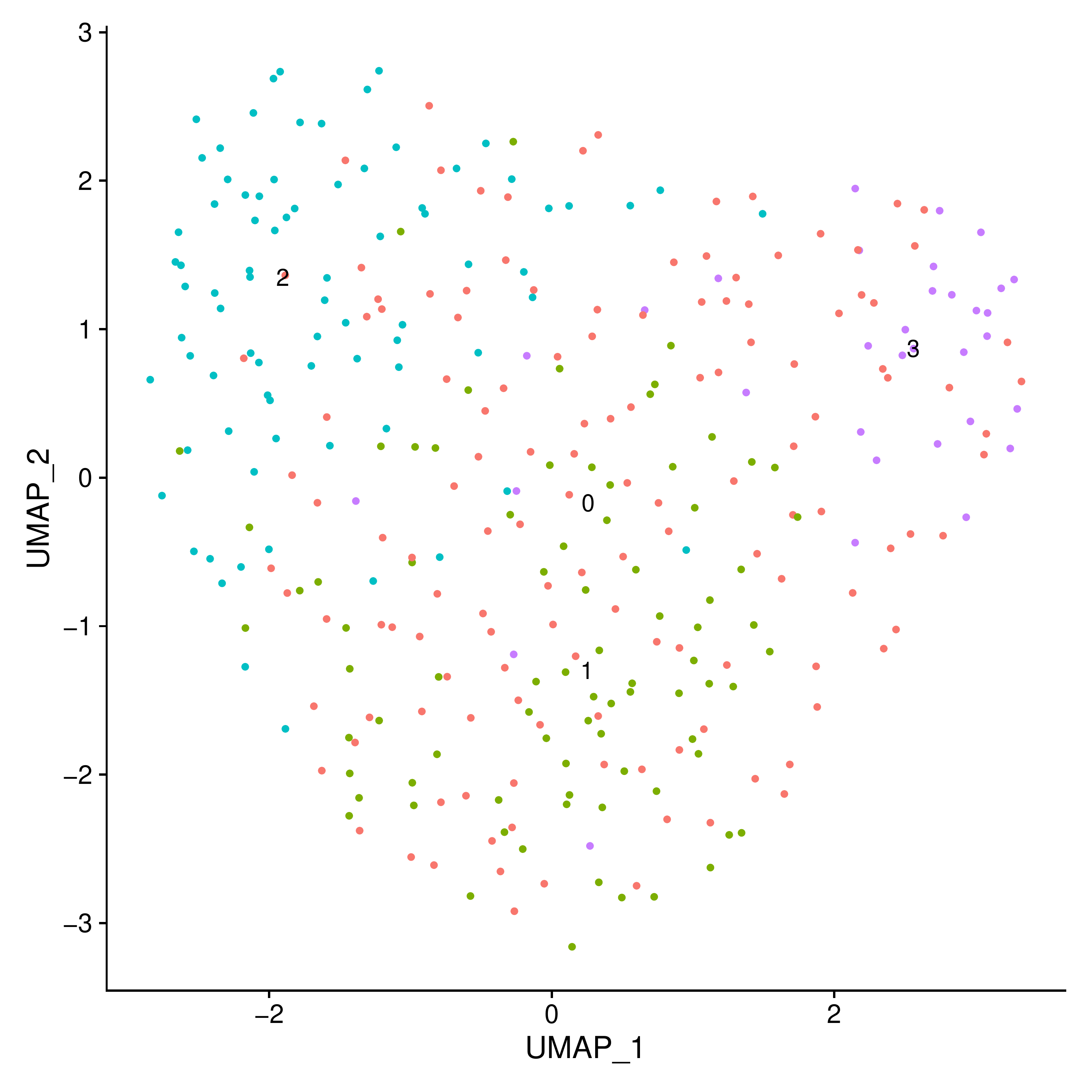

**Suppl. Fig. 19.** UMAP plot showing clustering of cells in the first freshwater sample, resolution = 0.9.

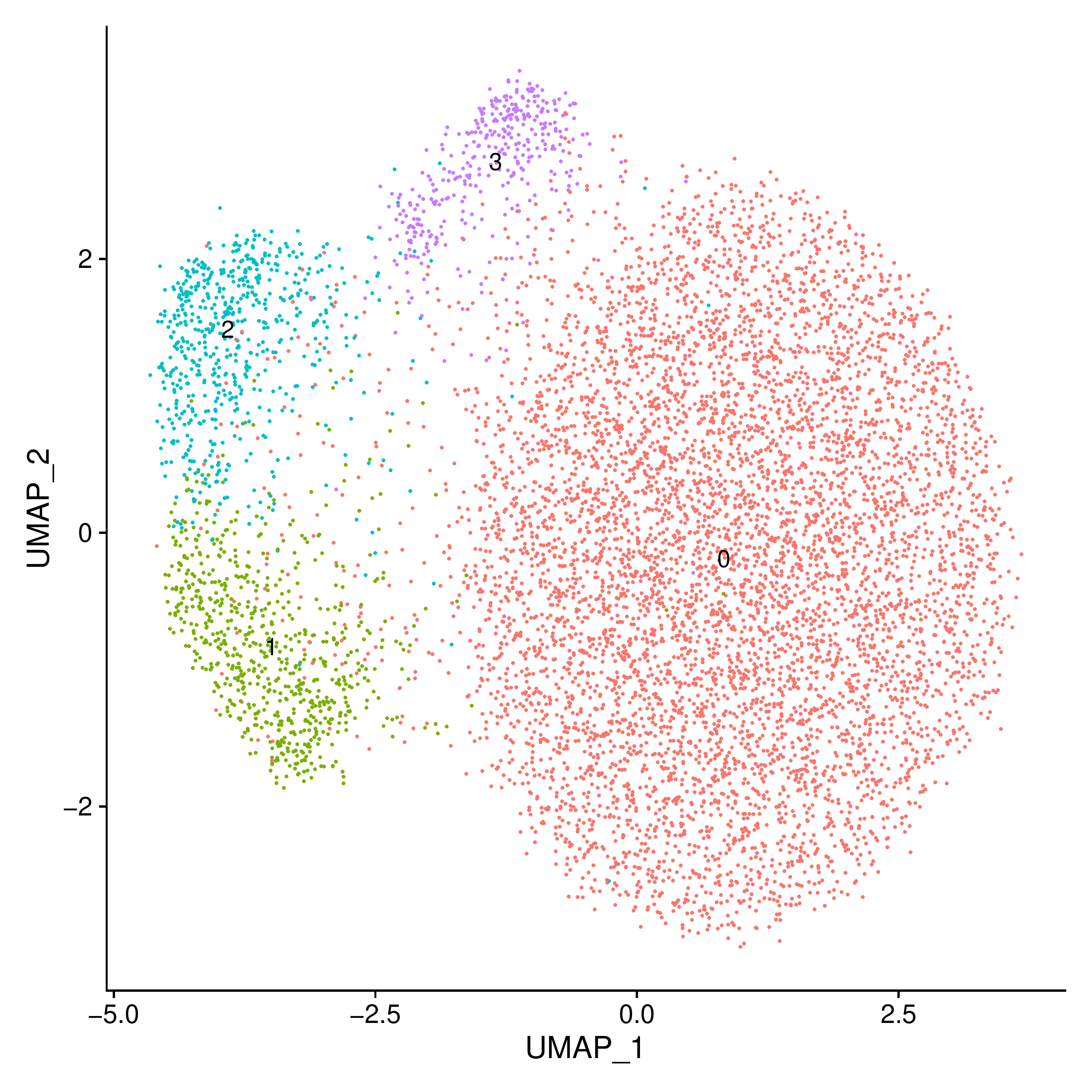

**Suppl. Fig. 20.** UMAP plot showing clustering of cells in the second freshwater sample, resolution = 0.6.

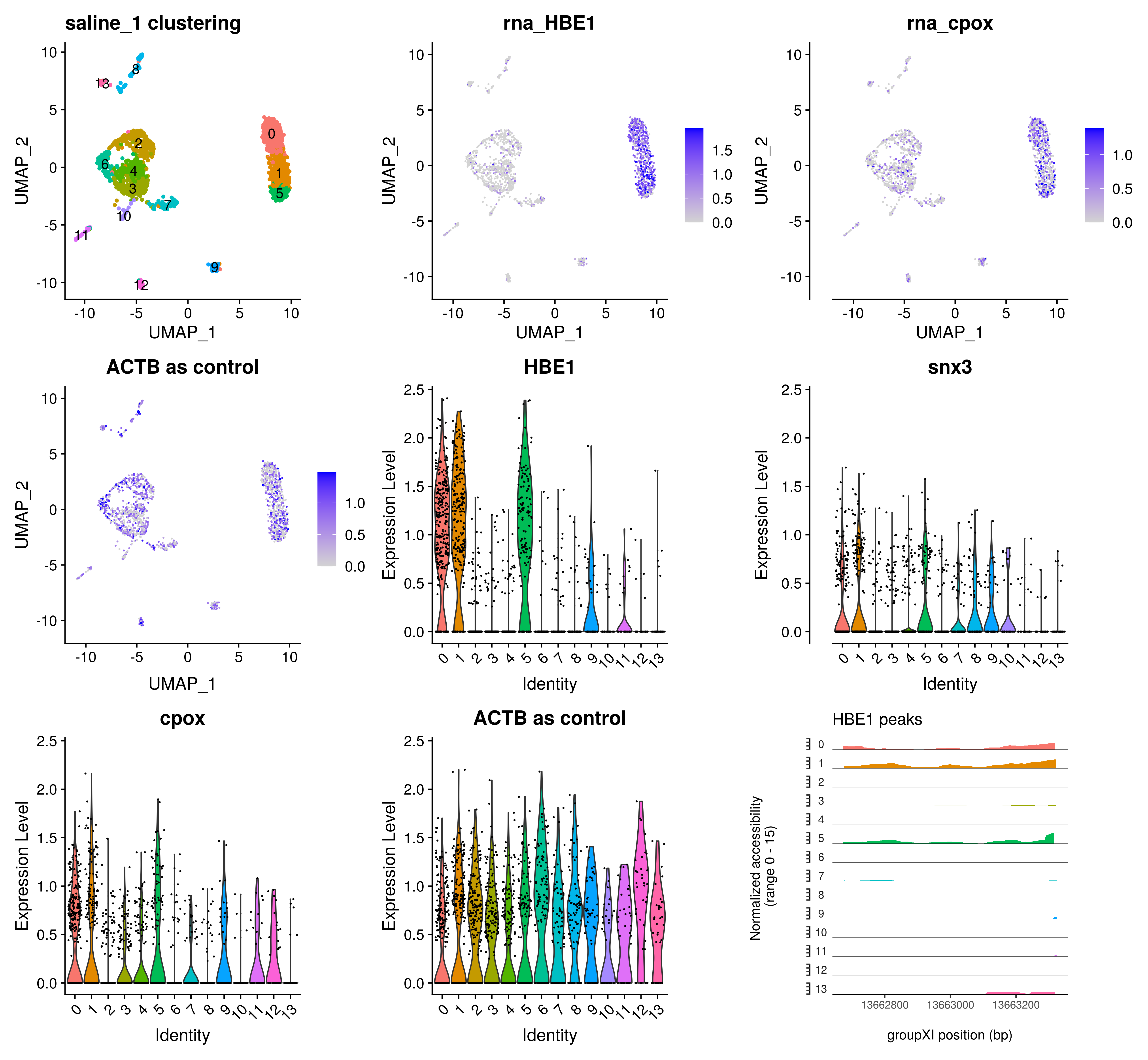

**Suppl. Fig. 21.** Identification of cell clusters corresponding to erythrocytes in the first marine sample using the activity of genes related to heme metabolism as well as open chromatin coverage of the *HBE1* gene. The activity of the *ACTB* gene was used as a control. Gene activities were calculated from chromatin accessibility data. Clusters 0, 1, and 5 were identified as erythrocytes.

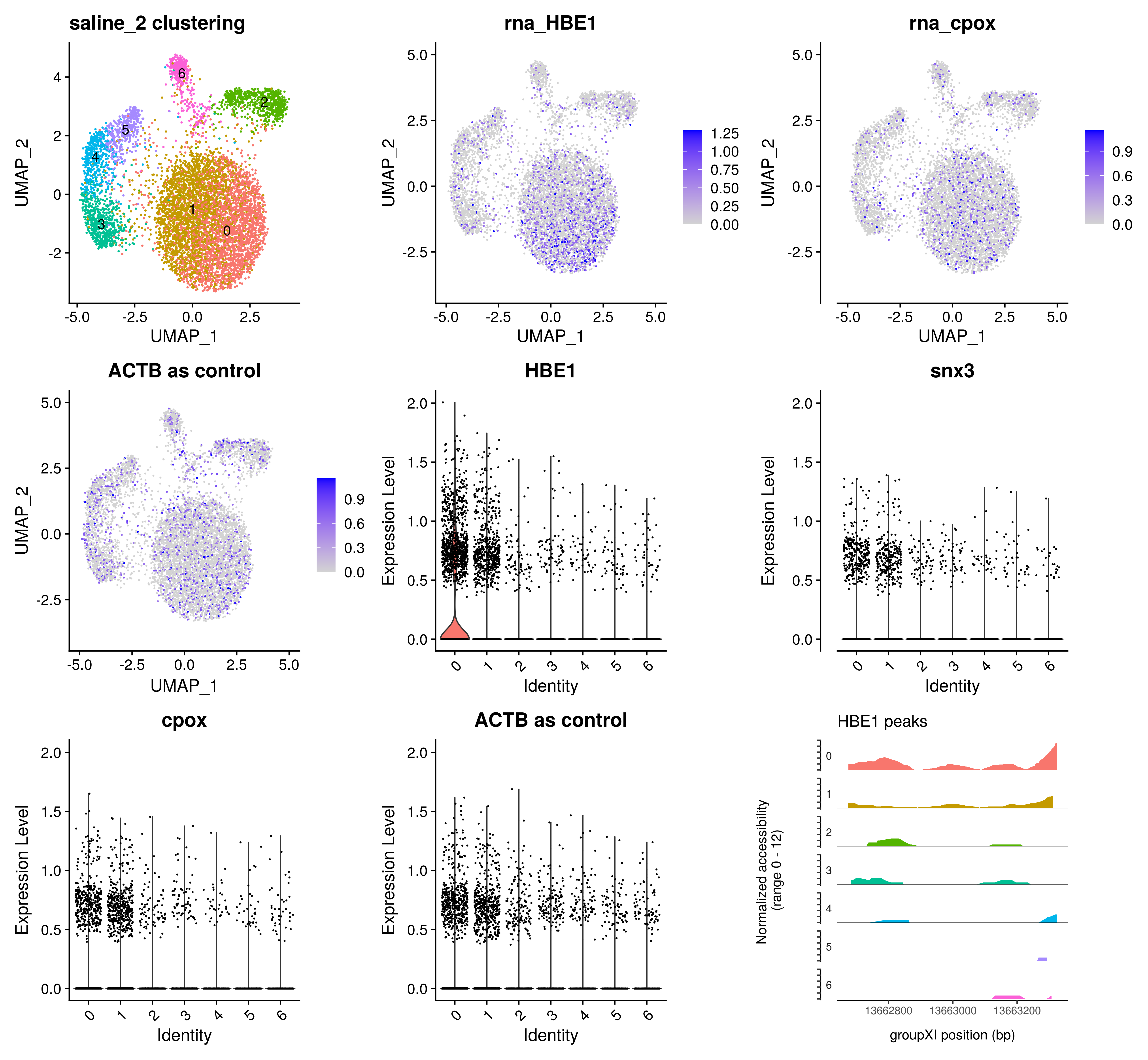

**Suppl. Fig. 22.** Identification of cell clusters corresponding to erythrocytes in the second marine sample using the activity of genes related to heme metabolism as well as open chromatin coverage of the *HBE1* gene. The activity of the *ACTB* gene was used as a control. Gene activities were calculated from chromatin accessibility data. Clusters 0 and 1 were identified as erythrocytes.

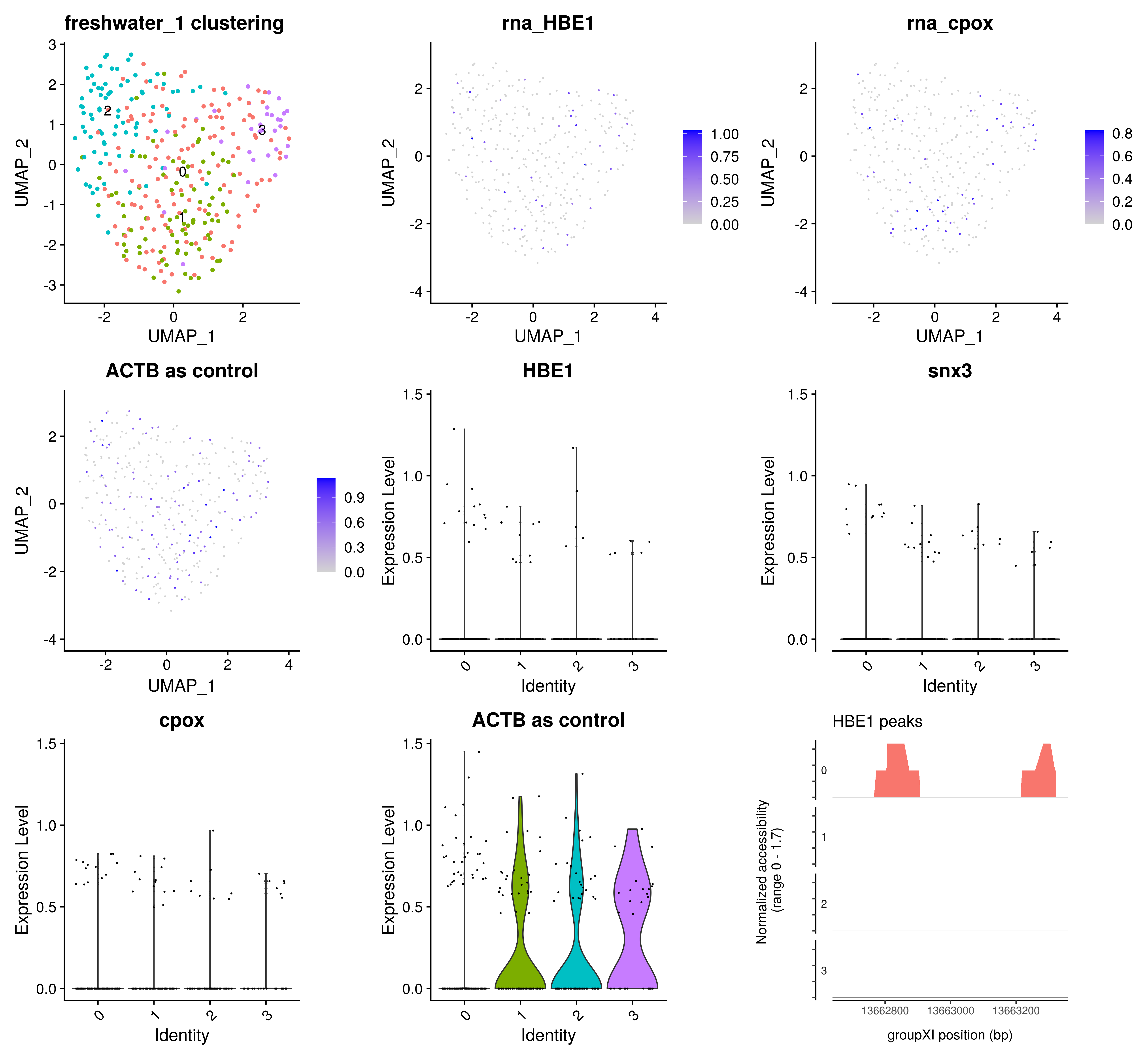

**Suppl. Fig. 23.** Identification of cell clusters corresponding to erythrocytes in the first freshwater sample using the activity of genes related to heme metabolism as well as open chromatin coverage of the *HBE1* gene. The activity of the *ACTB* gene was used as a control. Gene activities were calculated from chromatin accessibility data. Cluster 0 was identified as erythrocytes.

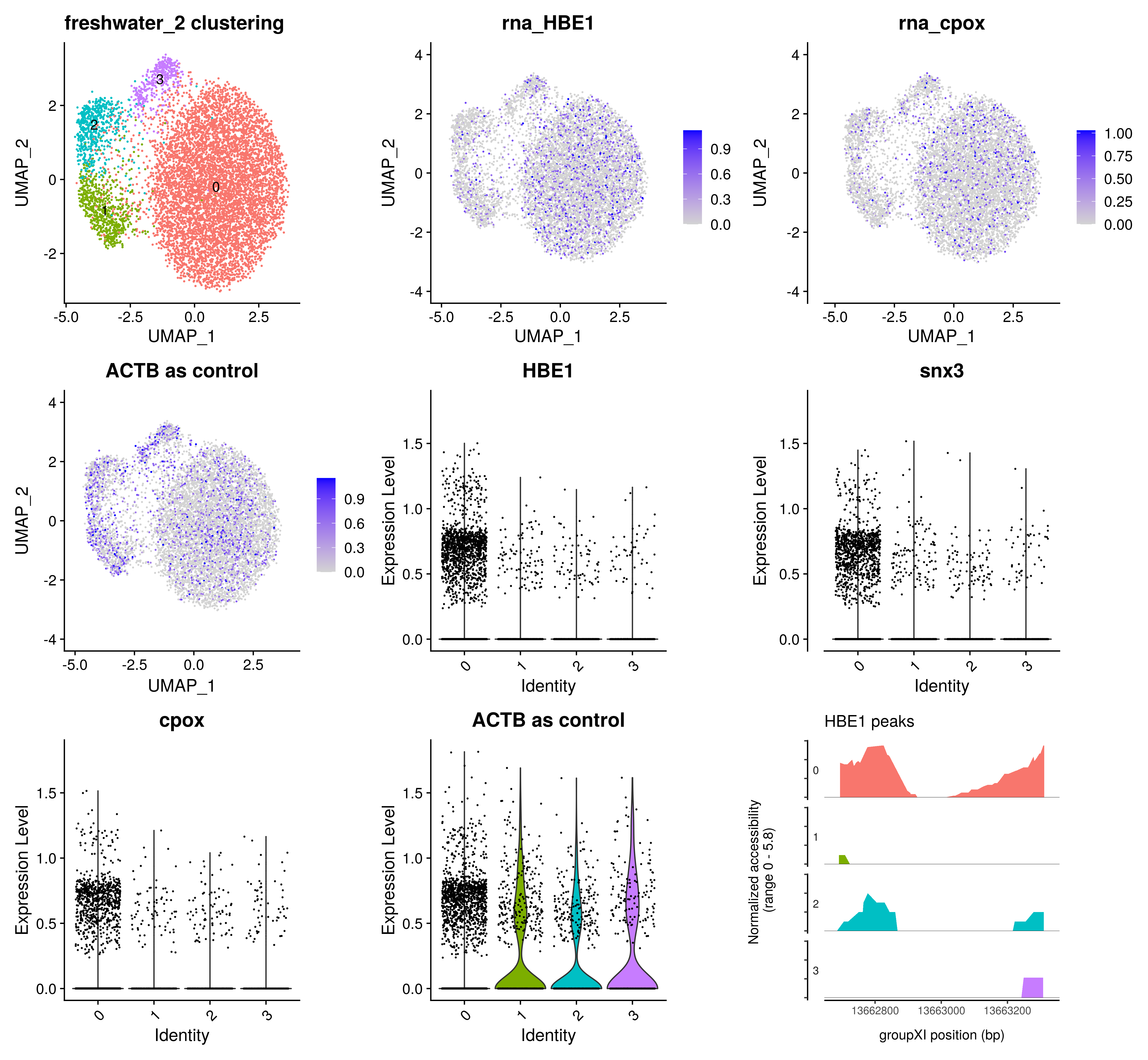

**Suppl. Fig. 24.** Identification of cell clusters corresponding to erythrocytes in the second freshwater sample using the activity of genes related to heme metabolism as well as open chromatin coverage of the *HBE1* gene. The activity of the *ACTB* gene was used as a control. Gene activities were calculated from chromatin accessibility data. Cluster 0 was identified as erythrocytes.

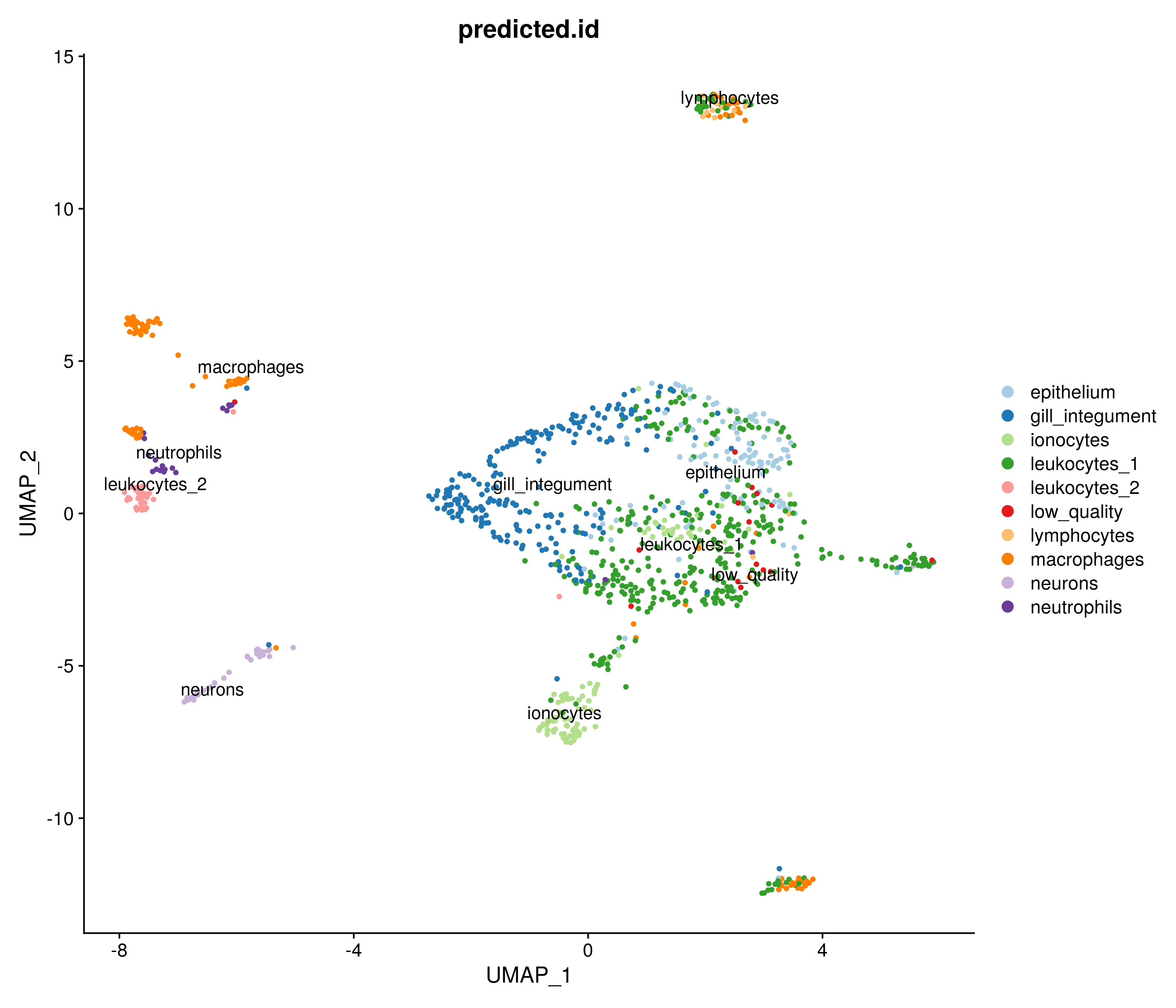

**Suppl. Fig. 25.** UMAP plot of direct label transfer from scRNA-seq data annotation of non-erythrocyte clusters to the first marine sample of scATAC-seq data on a cell-by-cell basis using the *FindTransferAnchors()* Signac function. Erythrocyte clusters from scATAC-seq data are not included.

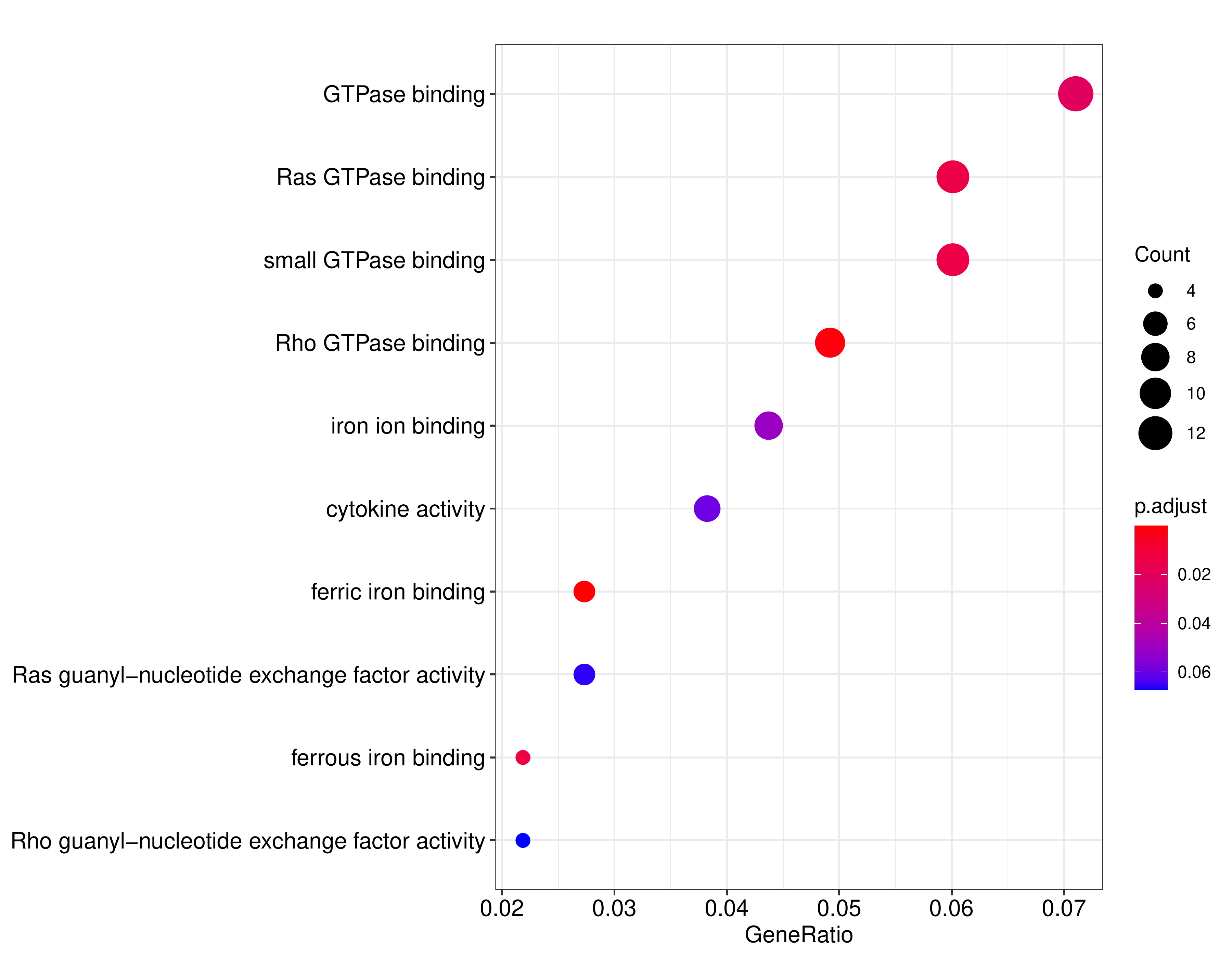

**Suppl. Fig. 26.** Top GO terms for freshwater-specific HVG. Two lists of differentially variable genes were generated with two methods: calculating peak heights per gene in each cell and calculating the number of non-zero peaks for a gene in each cell. Then the lists were filtered for F-ratio > 2 to obtain top genes with high variance in freshwater samples. Then the filtered top subsets of the lists were intersected. The resulting gene set was converted to Zebrafish orthologs and provided as an input to the GO enrichment analysis with the filtering for p-value < 0.1.

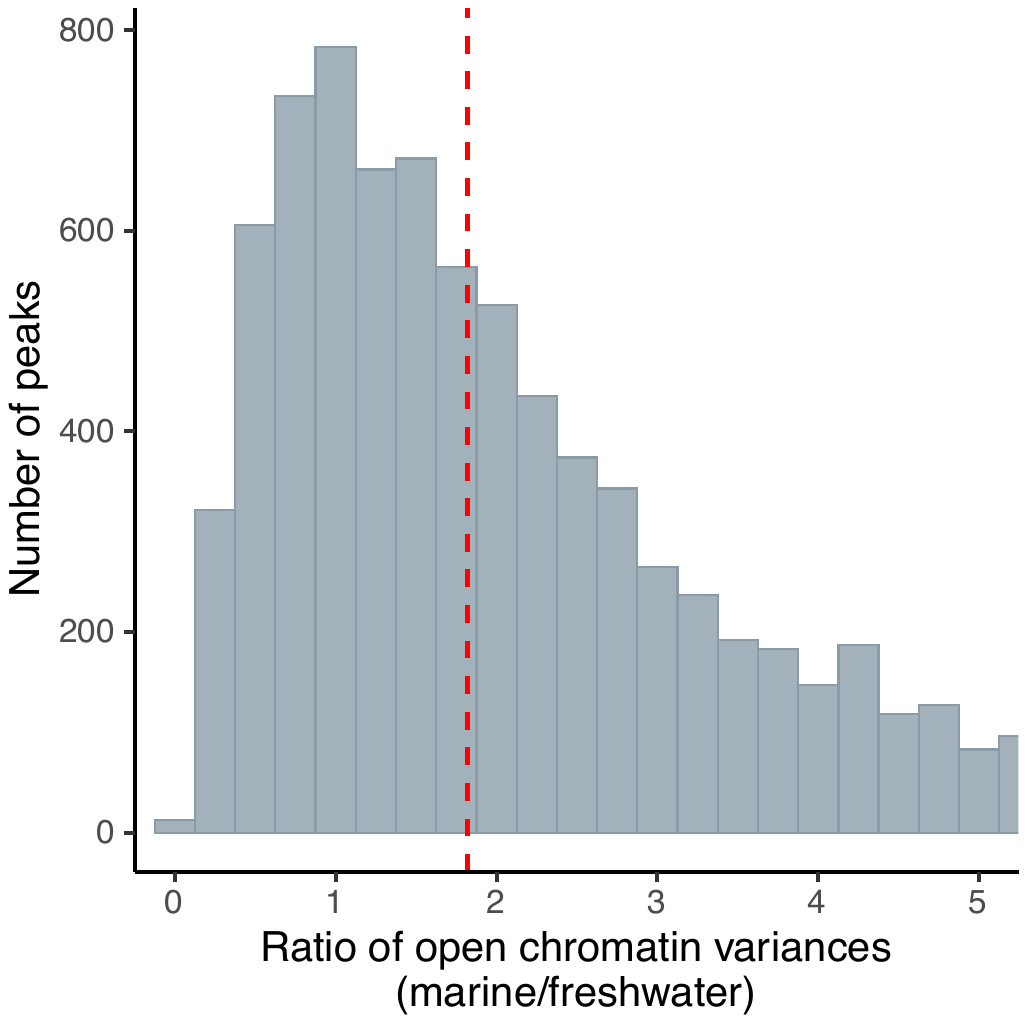

**Suppl. Fig. 27.** F-ratio (marine/freshwater) histogram calculated for all peaks and cells in erythrocyte clusters.

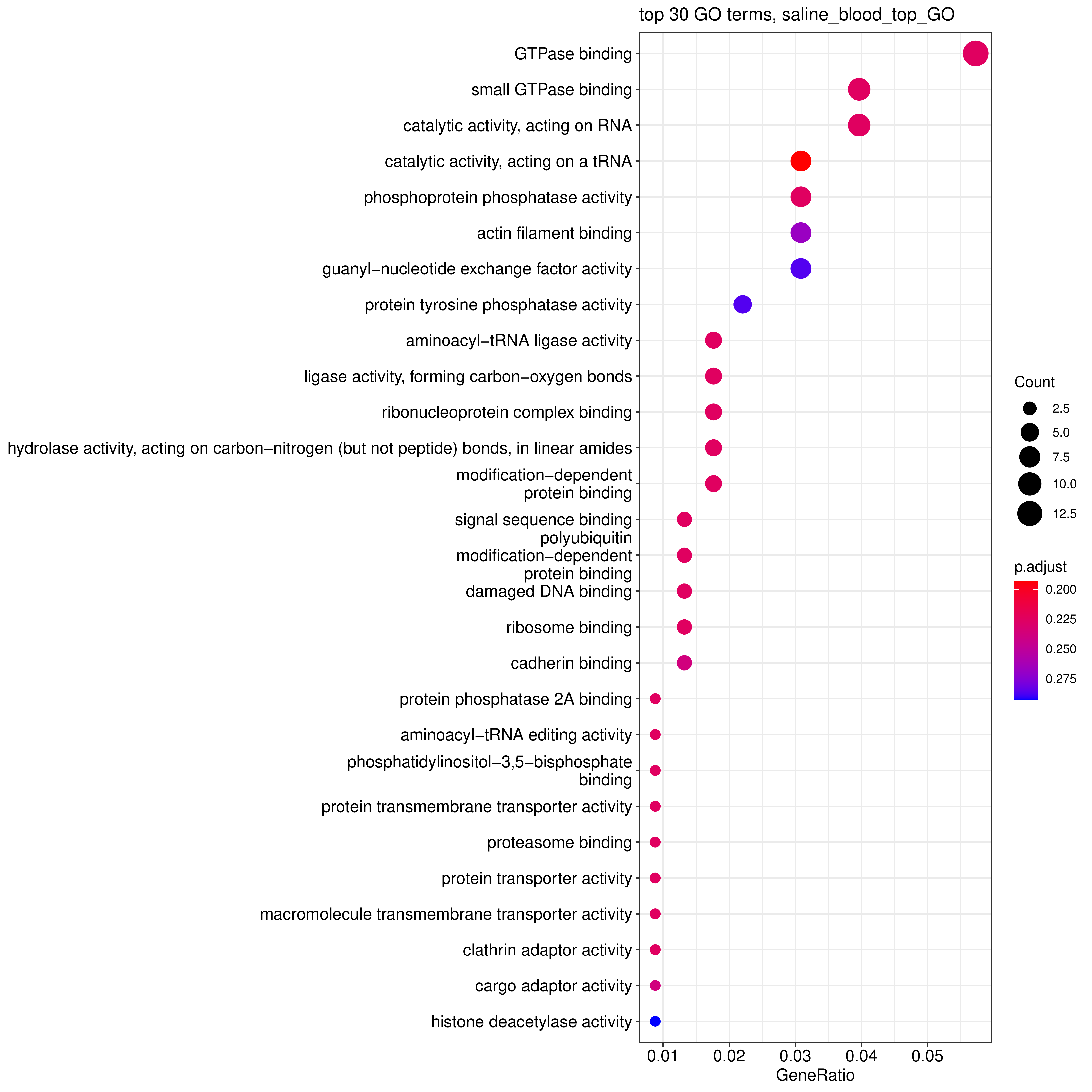

**Suppl. Fig. 28.** Top GO terms for marine-specific HVG. Two lists of differentially variable genes were generated with two methods: calculating peak heights per gene in each cell and calculating the number of non-zero peaks for a gene in each cell. Then the lists were filtered for F-ratio > 2 to obtain top genes with high variance in marine samples. Then the filtered top subsets of the lists were intersected. The resulting gene set was converted to Zebrafish orthologs and provided as an input to the GO enrichment analysis with the filtering for p-value < 0.1. Only cells from the erythrocyte cluster were used in the analysis.

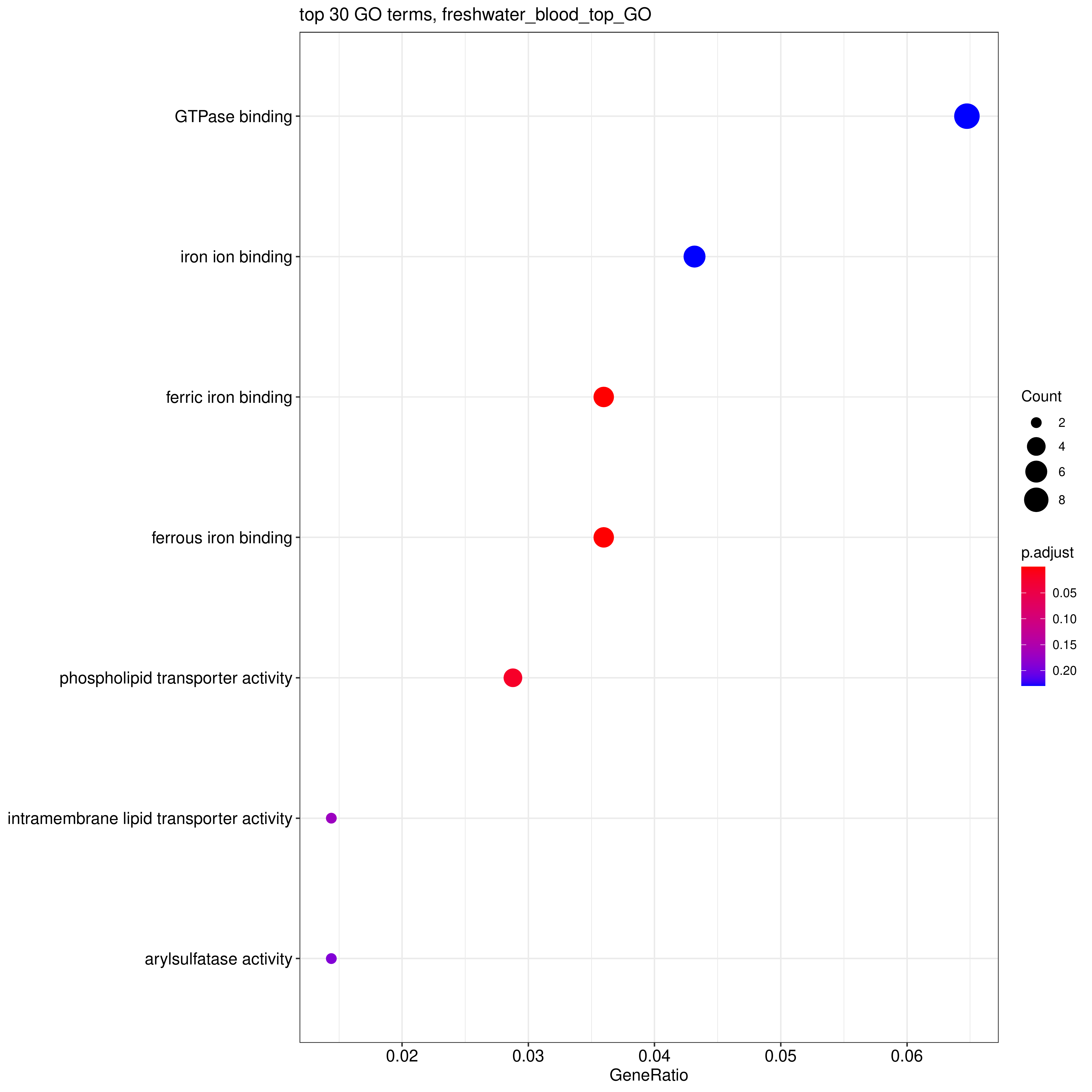

**Suppl. Fig. 29.** Top GO terms for freshwater-specific HVG. Two lists of differentially variable genes were generated with two methods: calculating peak heights per gene in each cell and calculating the number of non-zero peaks for a gene in each cell. Then the lists were filtered for F-ratio > 2 to obtain top genes with high variance in freshwater samples. Then the filtered top subsets of the lists were intersected. The resulting gene set was converted to Zebrafish orthologs and provided as an input to the GO enrichment analysis with the filtering for p-value < 0.1. Only cells from the erythrocyte cluster were used in the analysis.

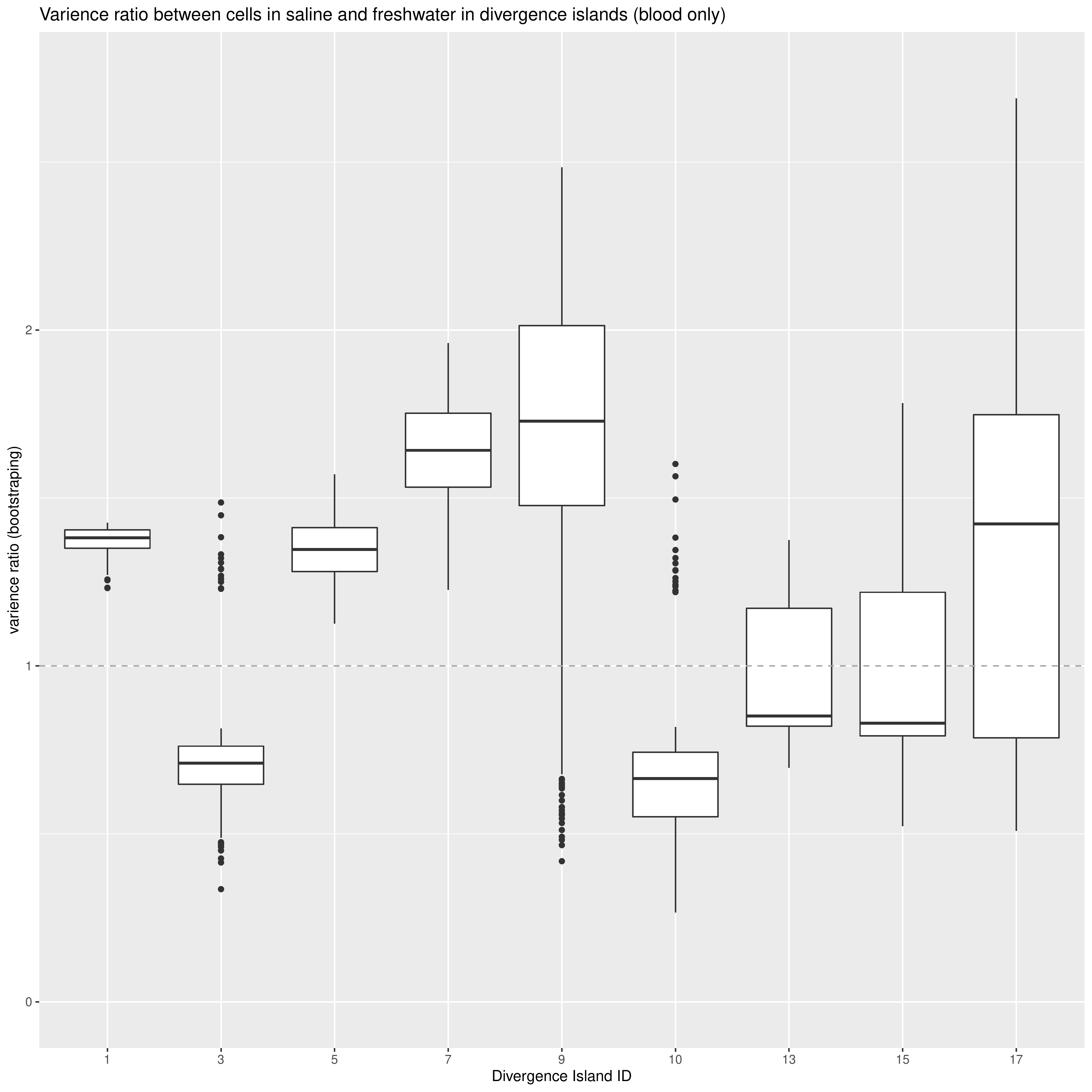

**Suppl. Fig. 30.** Box plots showing average F-ratios per DI in erythrocytes based on 1,000 bootstraps. F-ratio for divergence islands was calculated in the following way. First, the number of peaks with non-zero counts overlapping the DI was calculated per DI per cell to produce a DIs/cells matrix. Marine and freshwater cells were downsampled to a random set of 6,000 cells each to balance the number of cells. Next, F-ratio per DI was calculated as a ratio of variances between marine and freshwater cells in this matrix. Downsampling and F-ratio calculation were repeated 1,000 times. Only cells from erythrocyte clusters were used.

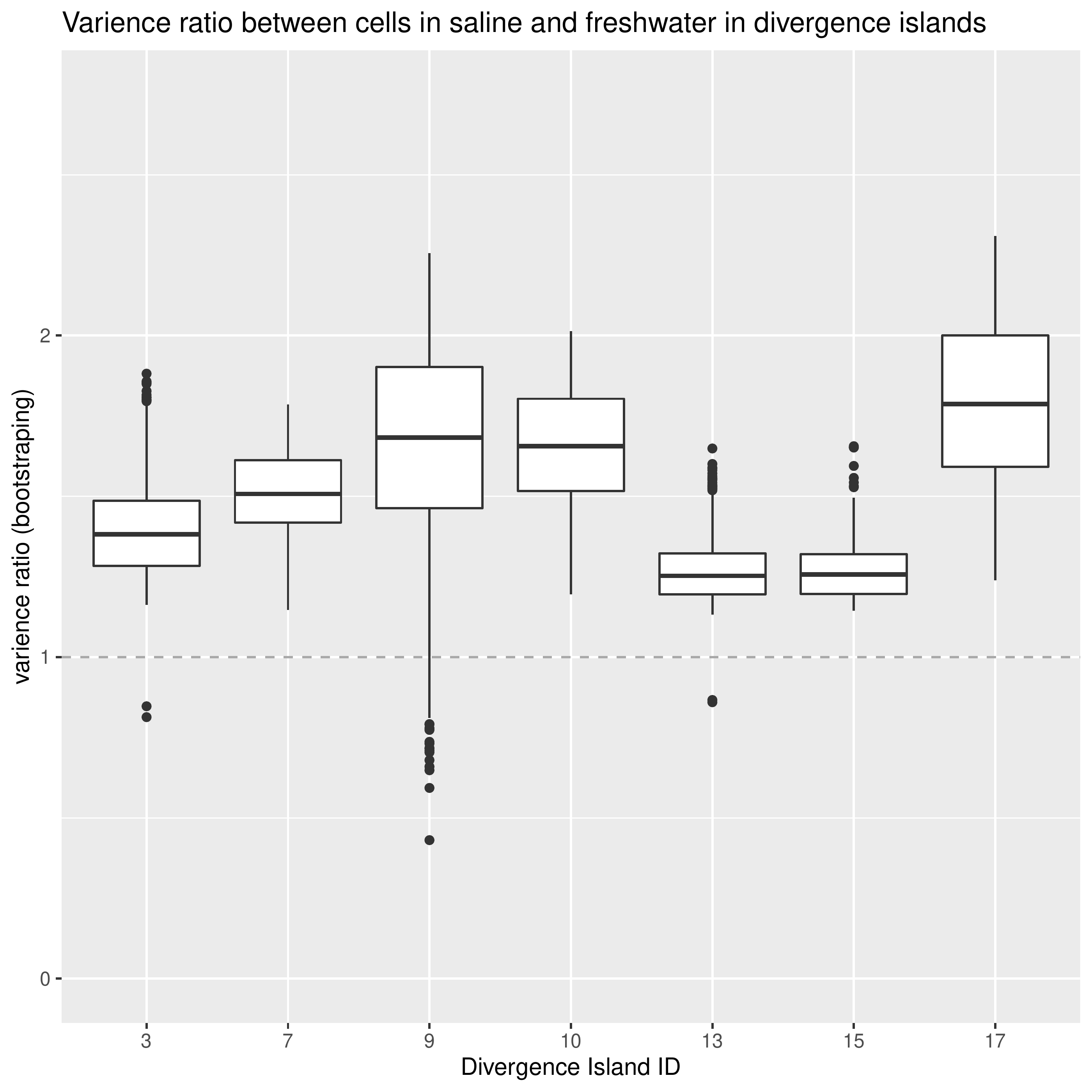

**Suppl. Fig. 31.** Box plots showing average F-ratios per DI in all cells based on 1,000 bootstraps. F-ratio for divergence islands was calculated in the following way. First, the number of peaks with non-zero counts overlapping the DI was calculated per DI per cell to produce a DIs/cells matrix. Marine and freshwater cells were downsampled to a random set of 6,000 cells each to balance the number of cells. Next, F-ratio per DI was calculated as a ratio of variances between marine and freshwater cells in this matrix. Downsampling and F-ratio calculation were repeated 1,000 times. All cells from the filtered scATAC-seq dataset were used.

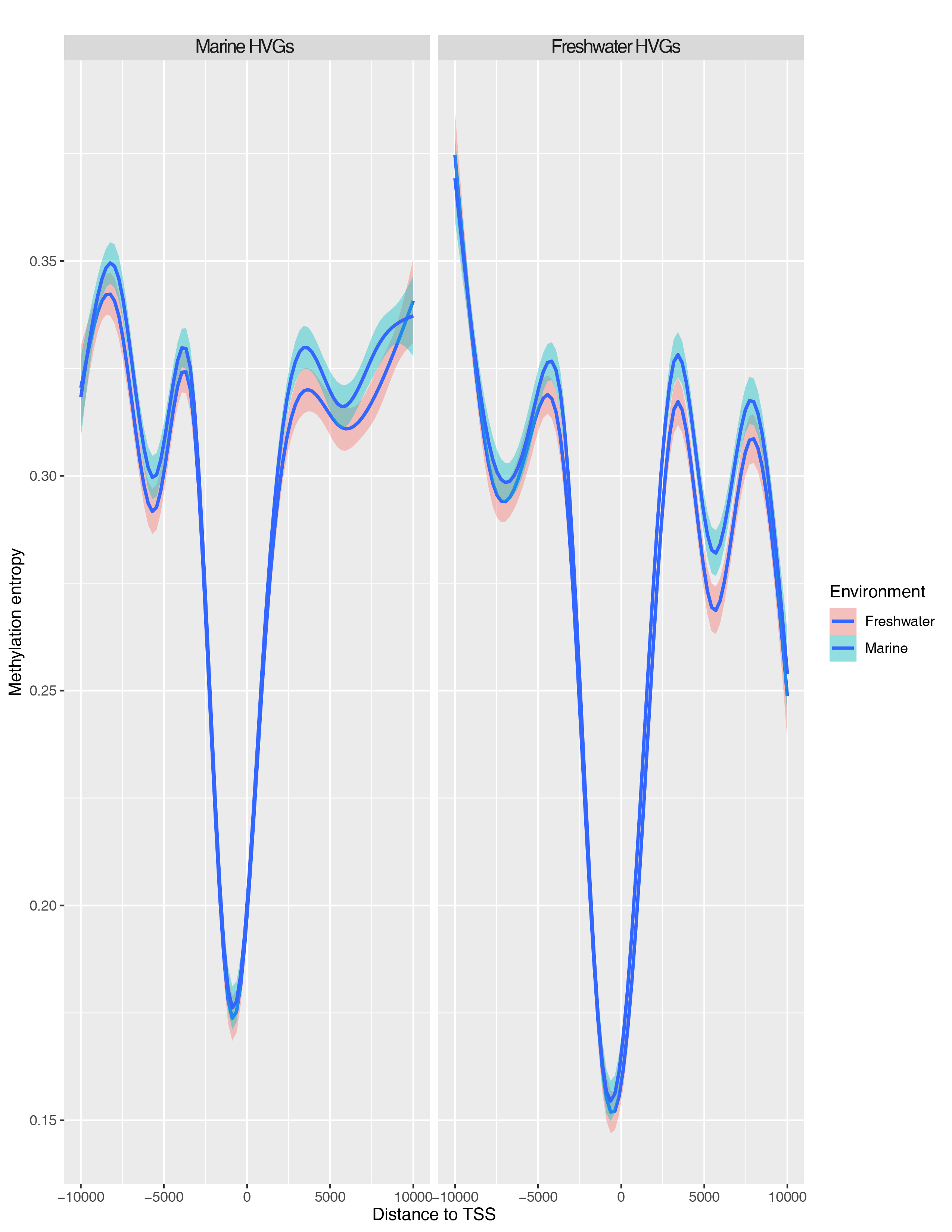

**Suppl. Fig. 32.** Methylation entropy distribution near TSS of differentially variable genes. Two types of genes were used: showing increased transcriptional variance in marine (marine HVGs, left) and freshwater (freshwater HVGs, right) environments.

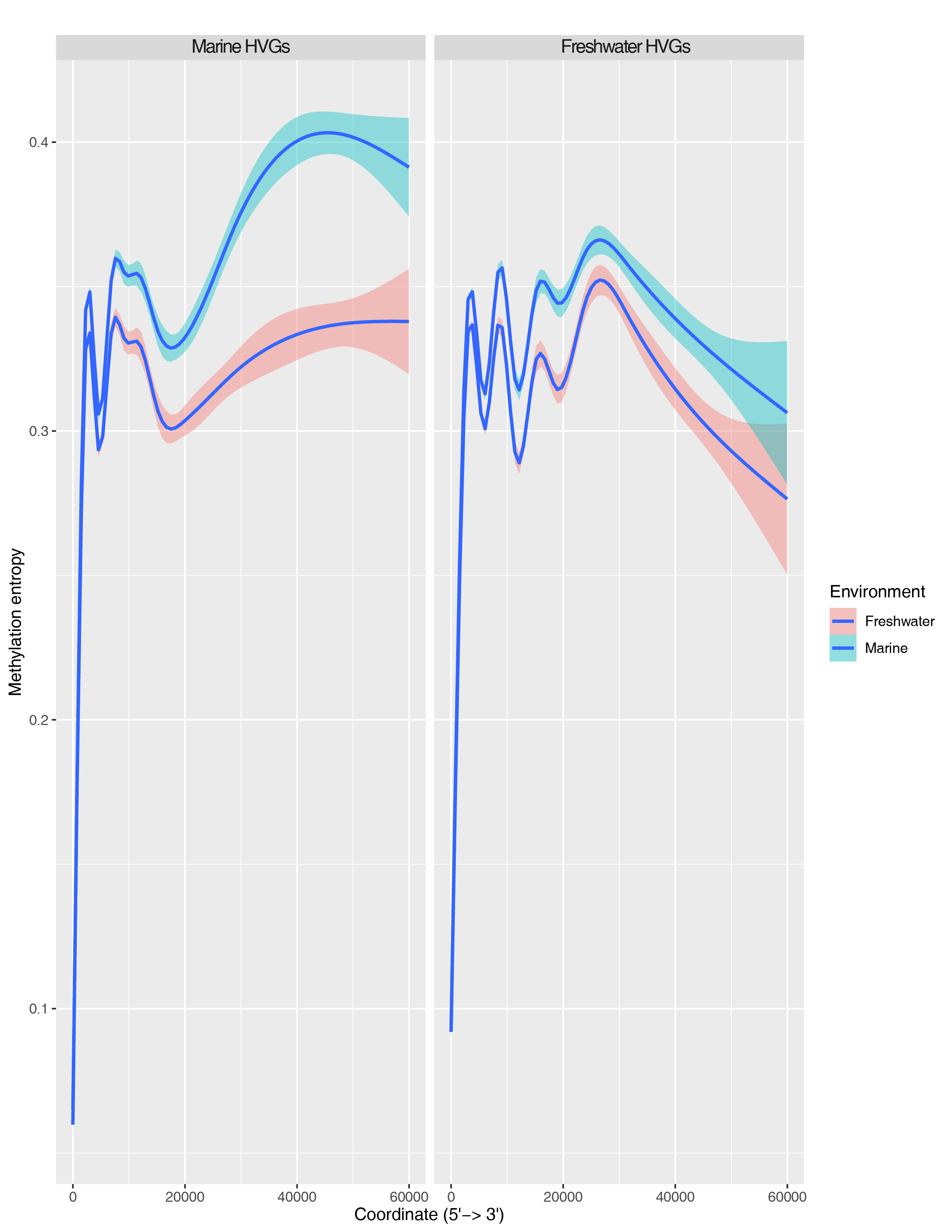

**Suppl. Fig. 33.** Methylation entropy in bodies of differentially variable genes (nucleotide scaling). Two types of genes were used: showing increased transcriptional variance in marine (marine HVGs, left) and freshwater (freshwater HVGs, right) environments.

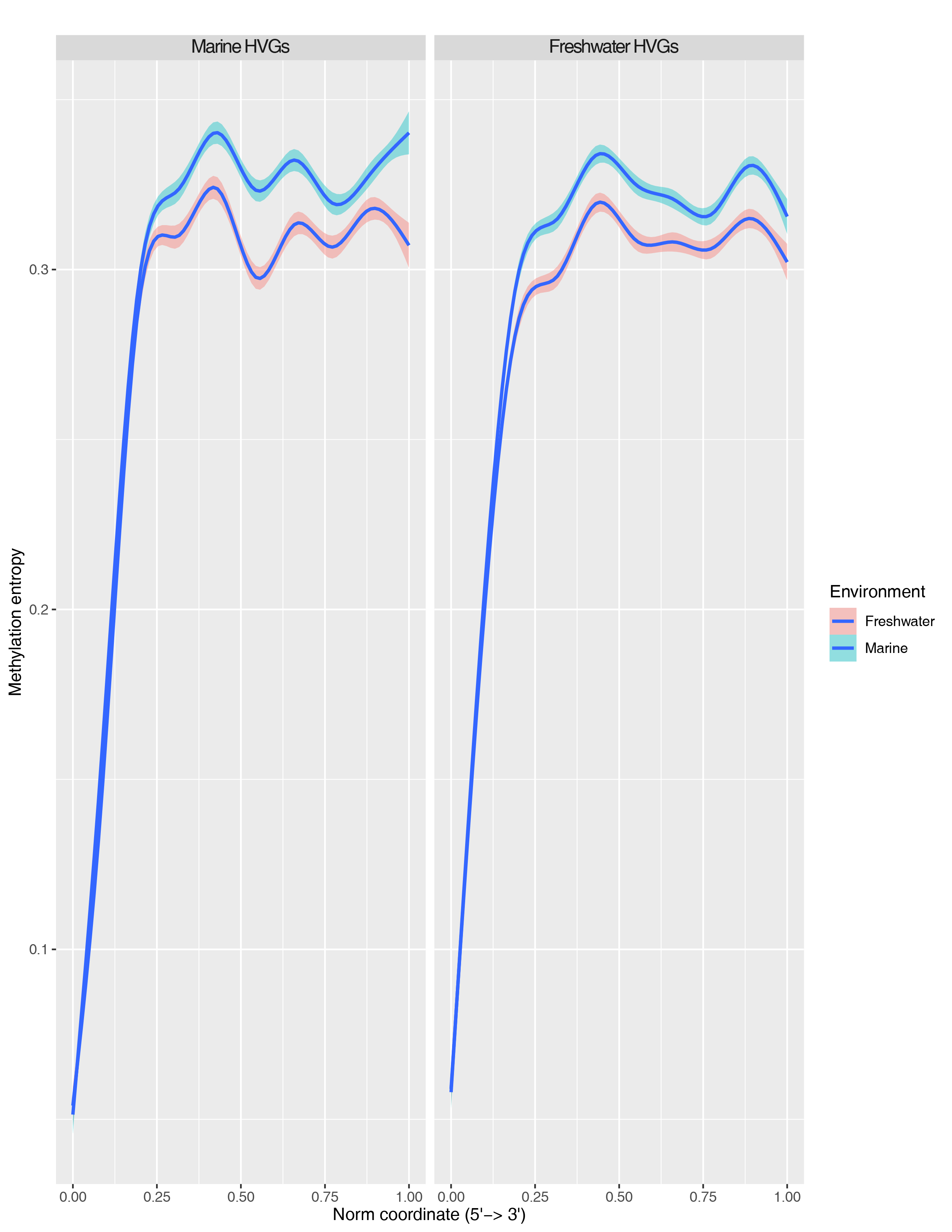

**Suppl. Fig. 34.** Methylation entropy in bodies of differentially variable genes (relative scaling). Two types of genes were used: showing increased transcriptional variance in marine (marine HVGs, left) and freshwater (freshwater HVGs, right) environments.

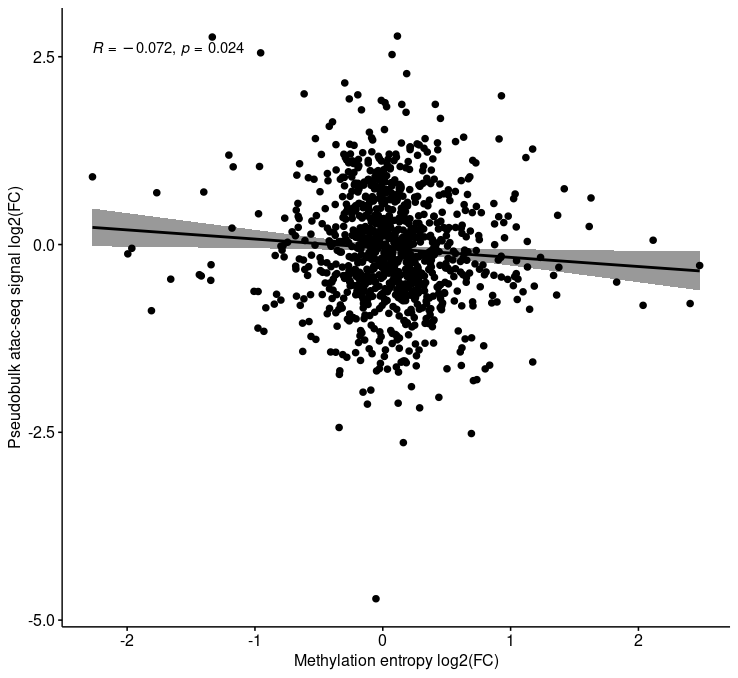

**Suppl. Fig. 35.** Correlation between pseudobulk atac-seq fold change and fold change of the averaged methylation entropy values. The peaks were selected randomly.

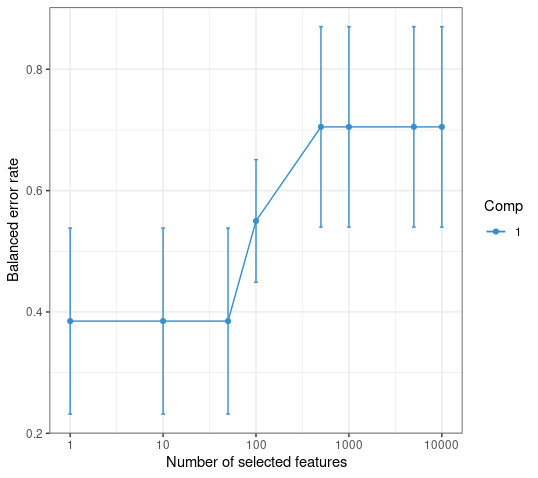

**Suppl. Fig. 36.** Determining of the variables (peaks) number for the most accurate division into classes. A 5-fold cross-validation with 50 repeats was performed.

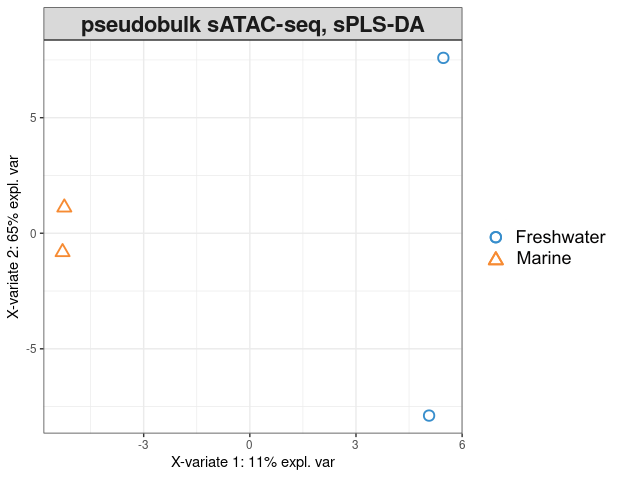

**Suppl. Fig. 37.** sPLS-DA final model with top 50 selected peaks.

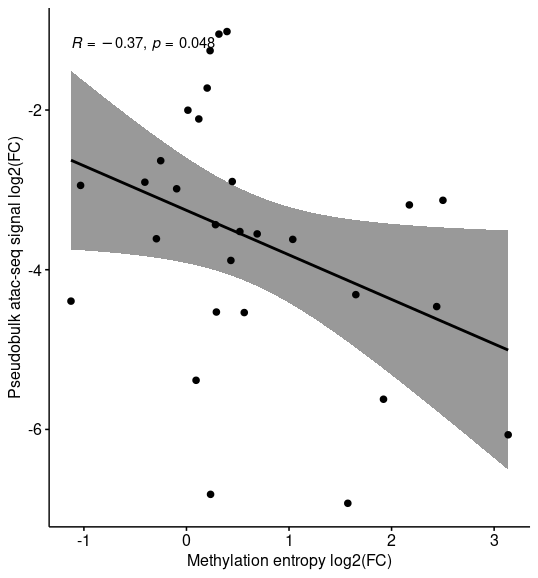

**Suppl. Fig. 38.** Correlation between pseudobulk atac-seq fold change and fold change of the averaged methylation entropy values. The top 50 peaks were obtained by sPLS-DA.

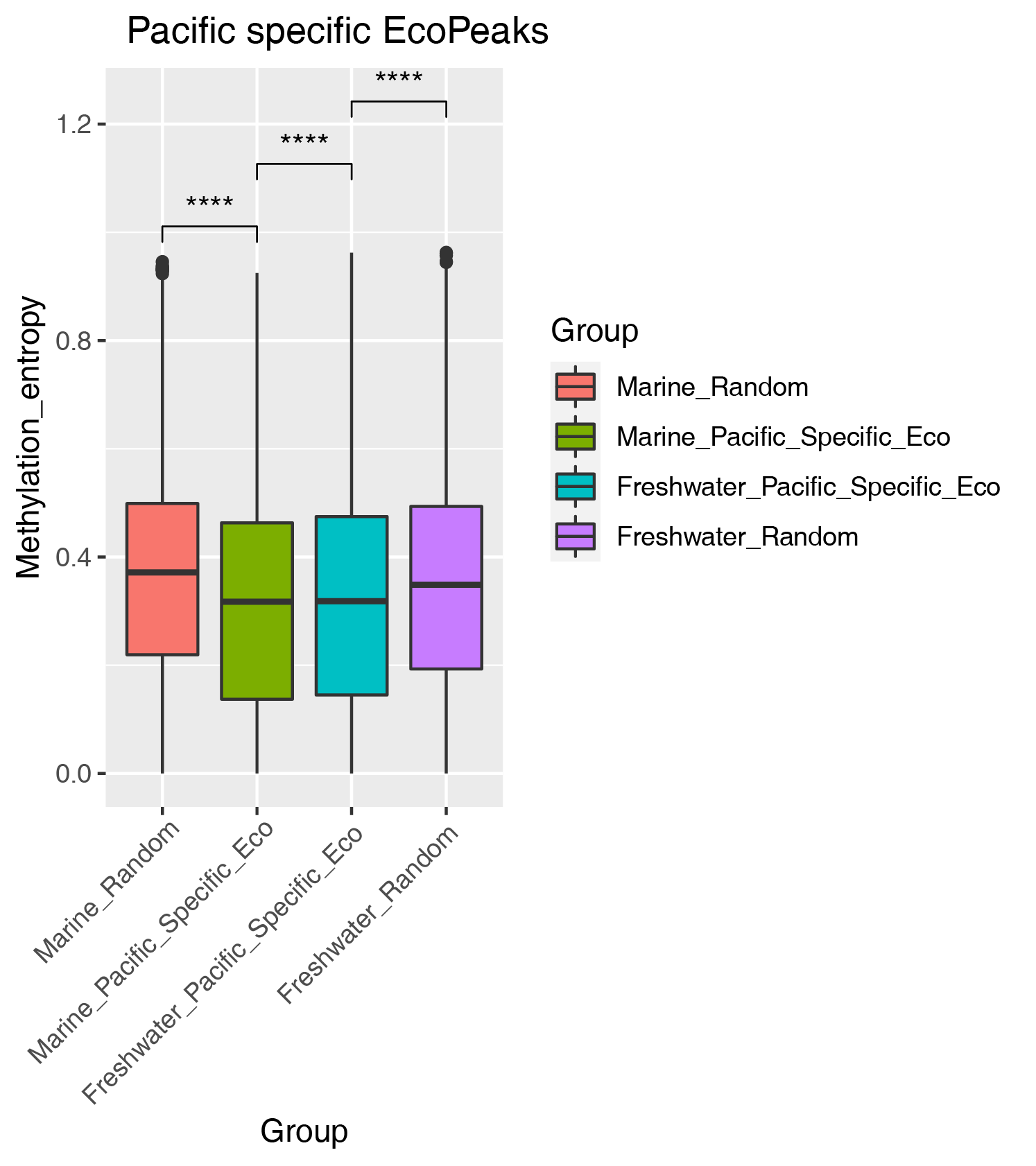

**Suppl. Fig. 39.** DNA methylation entropy in the Pacific specific EcoPeaks.

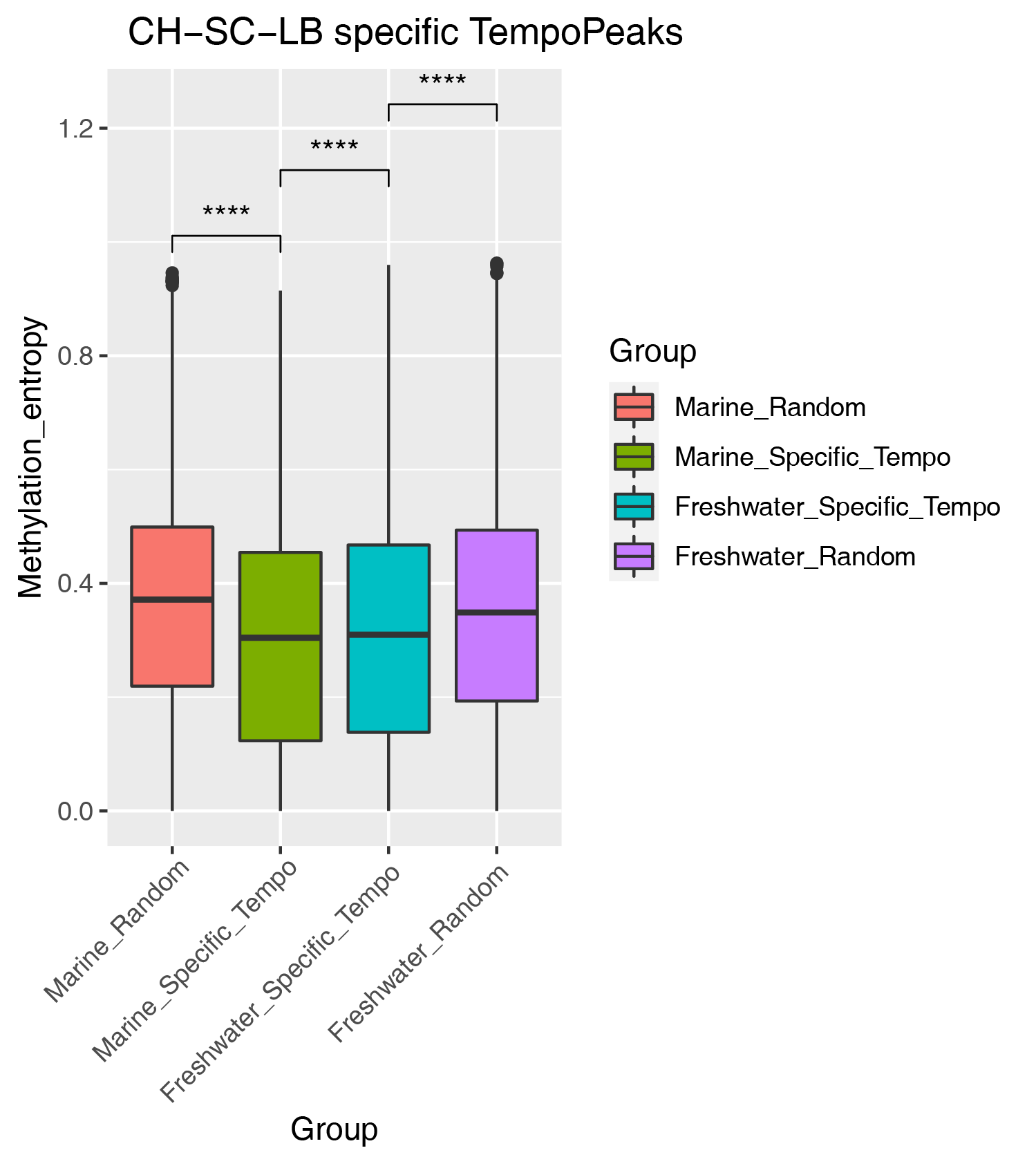

**Suppl. Fig. 40.** DNA methylation entropy in the specific TempoPeaks.

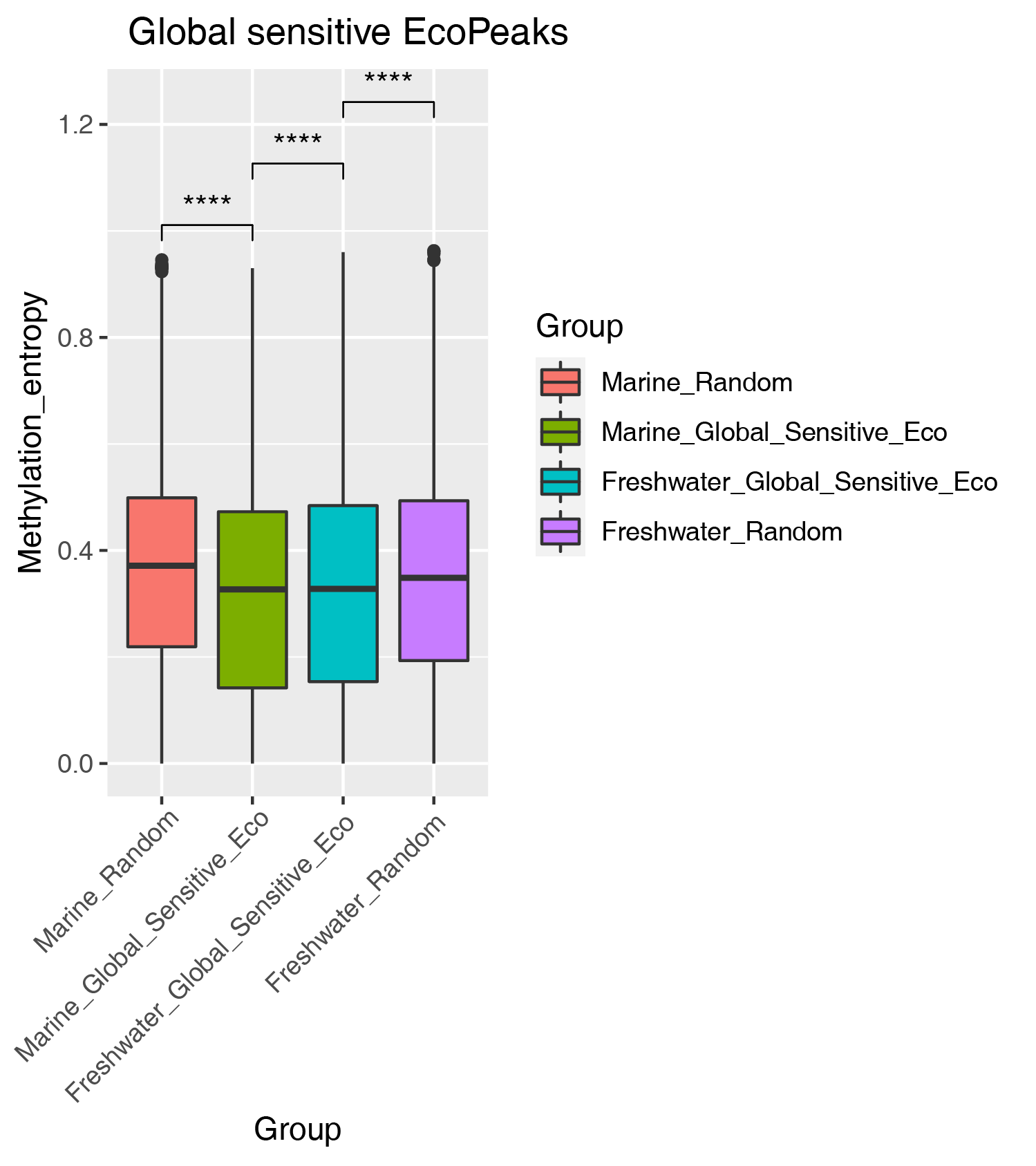

**Suppl. Fig. 41.** DNA methylation entropy in the Global sensitive EcoPeaks.

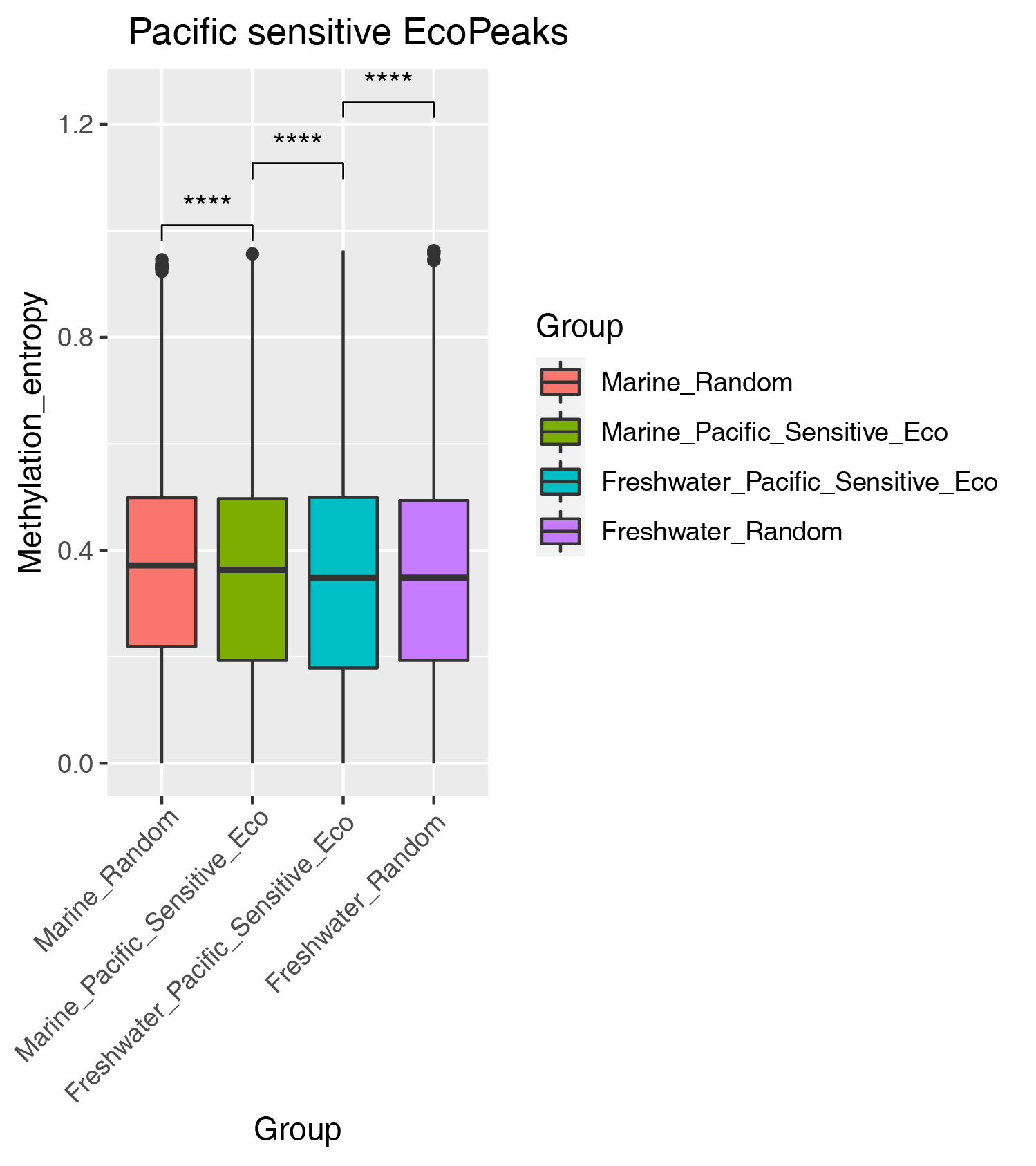

**Suppl. Fig. 42.** DNA methylation entropy in the Pacific sensitive EcoPeaks.

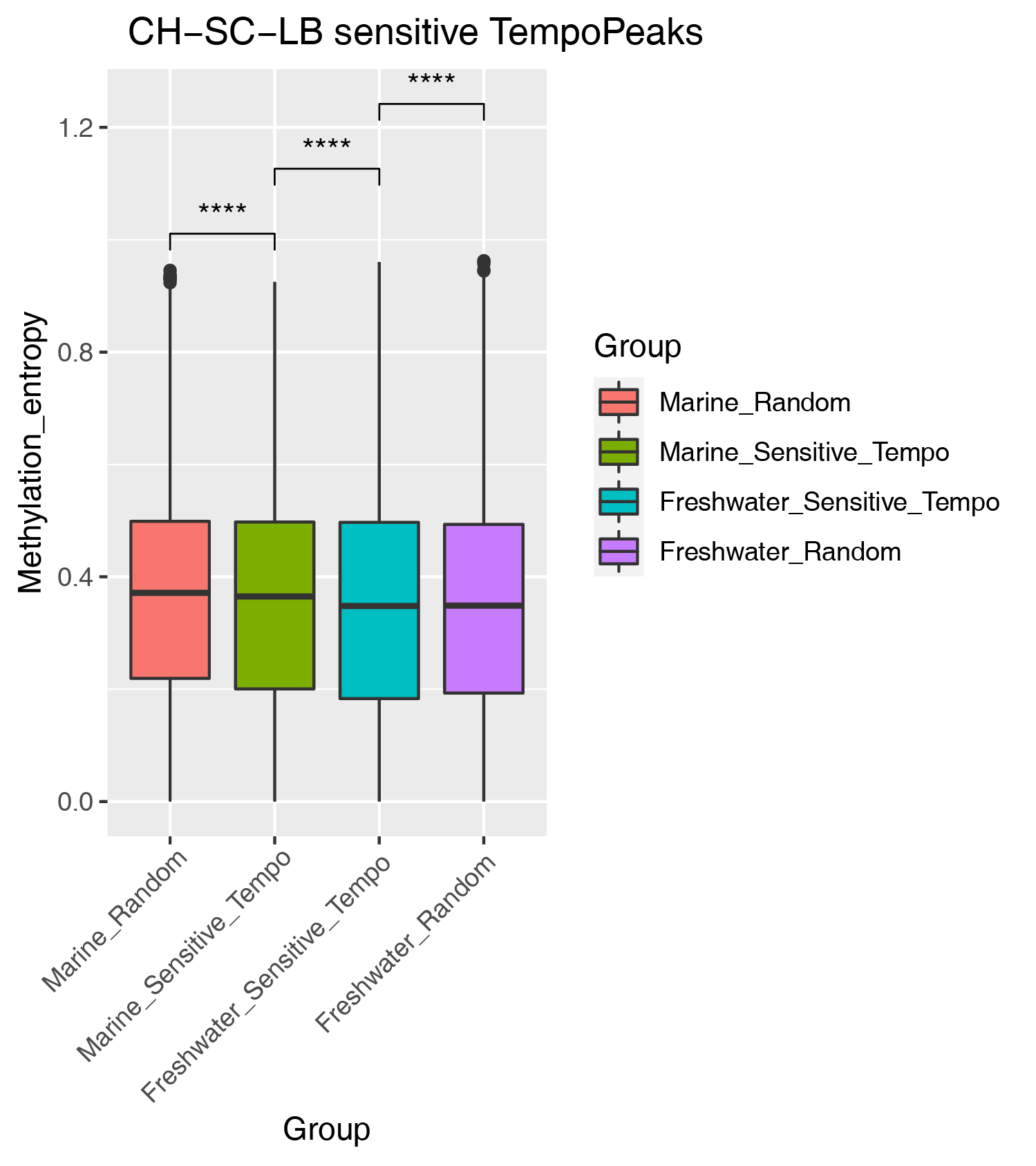

**Suppl. Fig. 43.** DNA methylation entropy in the sensitive TempoPeaks.

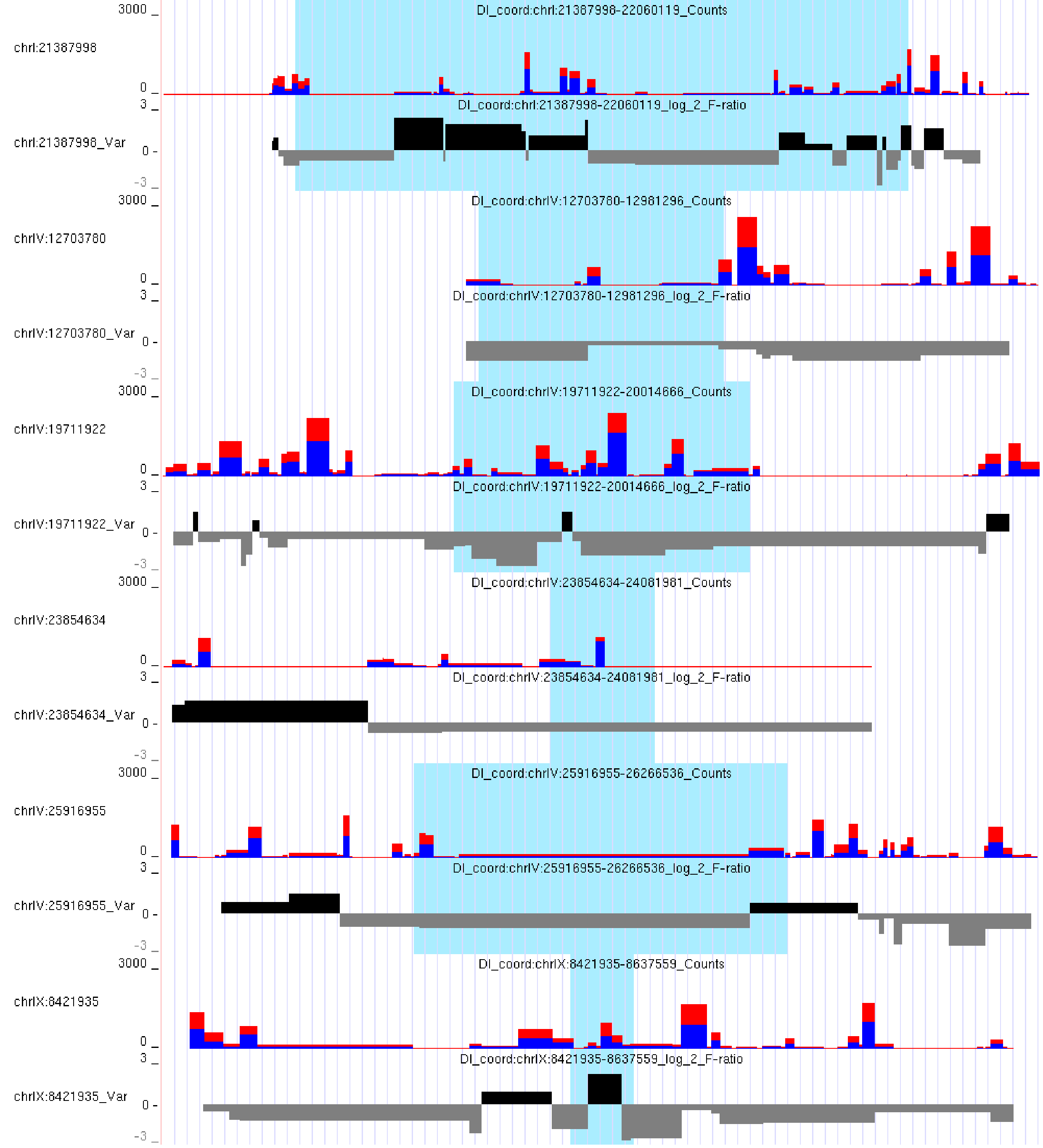

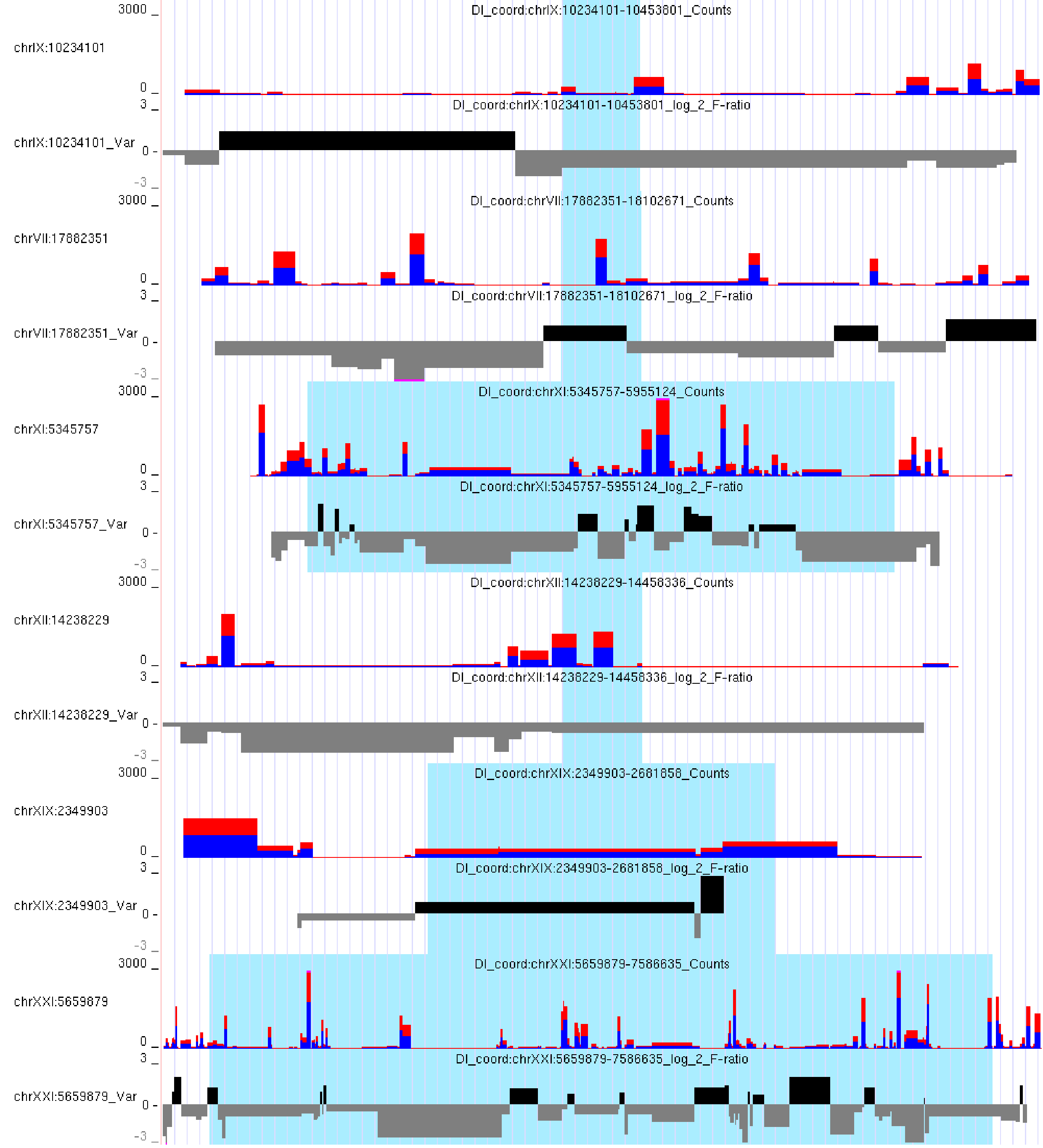

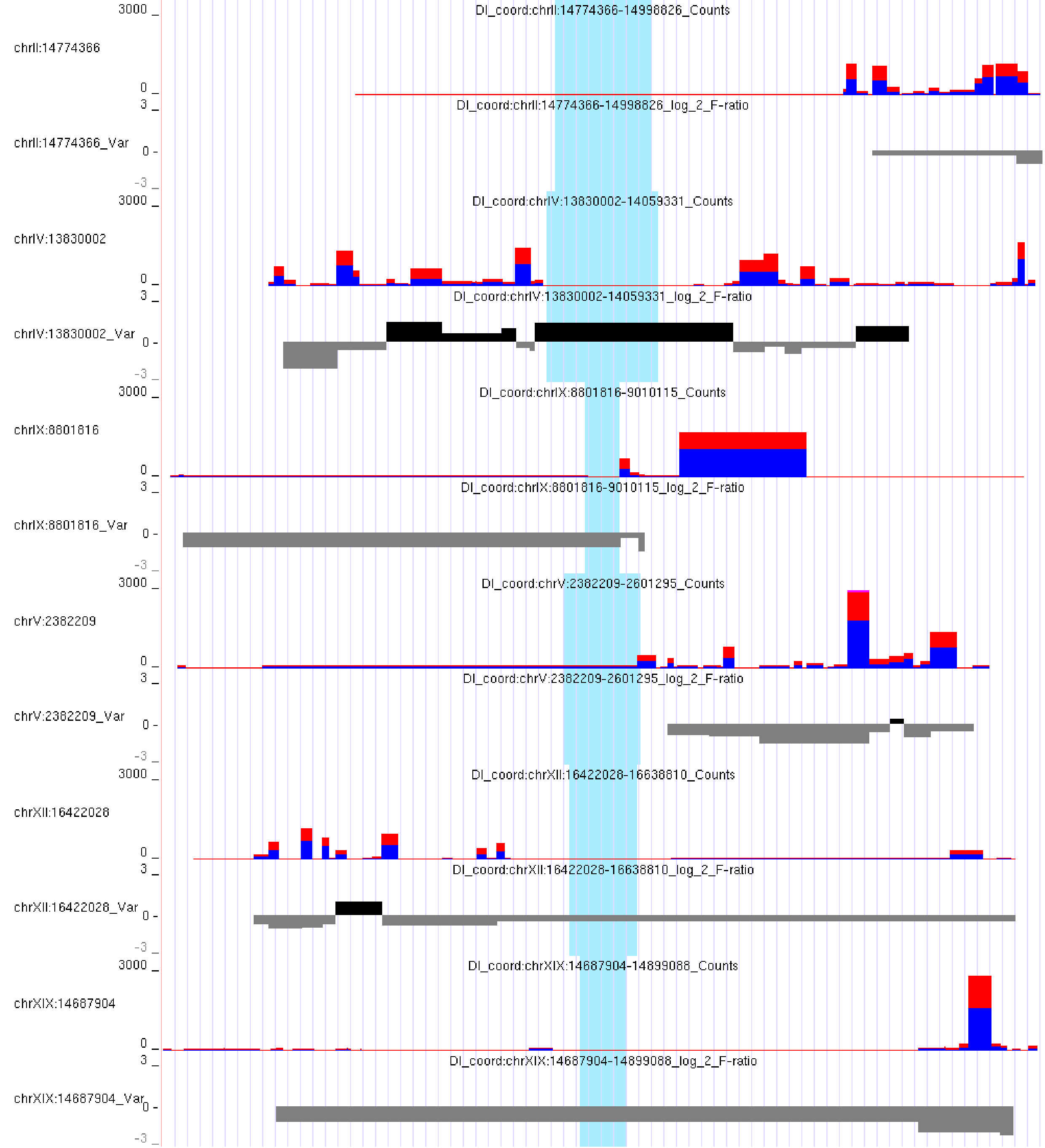

**Suppl. Fig. 44.** Chromatin accessibility in DIs. Red and blue bars represent the coverage of ATAC-seq peaks in freshwater and marine environments. Black and gray bars represent changes in the variance of ATAC-seq peaks (F-ratio = freshwater variance / marine variance).

### **SUPPLEMENTARY TABLES**

**Suppl. Table 1.** Cell Ranger run summary for the scRNA-seq dataset.

**Suppl. Table 2.** Cluster markers of the whole scRNA-seq dataset after clustering with resolution = 0.1.

**Suppl. Table 3.** Markers of 804 cells from the scRNA-seq dataset without erythrocytes after clustering with resolution = 0.3 and cluster annotation.

**Suppl. Table 4.** F-ratio analysis output for HVG genes corresponding to figure 2A.

**Suppl. Table 5.** HVG genes used for the F-ratio analysis in figure 2A.

**Suppl. Table 6.** Cell Ranger run summary for the scATAC-seq dataset.

**Suppl. Table 7.** Parameters for QC filtration steps in Signac pipeline for each scATAC-seq sample.

**Suppl. Table 8.** Nuclei-targeted and stock concentrations during isolation of nuclei for scRNA-seq and scATAC-seq protocols.

**Suppl. Table 9.** Enrichment of known motifs for scATAC-seq peaks in the vicinity of marker SNPs.
